# Distributed network flows generate localized category selectivity in human visual cortex

**DOI:** 10.1101/2022.02.19.481103

**Authors:** Carrisa V. Cocuzza, Ruben. Sanchez-Romero, Takuya. Ito, Ravi D. Mill, Brian P. Keane, Michael W. Cole

## Abstract

A central goal of neuroscience is to understand how function-relevant brain activations are generated. Here we test the hypothesis that function-relevant brain activations are generated primarily by distributed network flows. We focused on visual processing in human cortex, given the long-standing literature supporting the functional relevance of brain activations in visual cortex regions exhibiting visual category selectivity. We began by using fMRI data from N=352 human participants to identify category-specific responses in visual cortex for images of faces, places, body parts, and tools. We then systematically tested the hypothesis that distributed network flows can generate these localized visual category selective responses. This was accomplished using a recently developed approach for simulating – in a highly empirically constrained manner – the generation of task-evoked brain activations by modeling activity flowing over intrinsic brain connections. We next tested refinements to our hypothesis, focusing on how stimulus-driven network interactions initialized in V1 generate downstream visual category selectivity. We found evidence that network flows directly from V1 were sufficient for generating visual category selectivity, but that additional, globally distributed (whole-cortex) network flows increased category selectivity further. Using null network architectures we also found that each region’s unique intrinsic “connectivity fingerprint” was key to the generation of category selectivity. These results generalized across regions associated with all four visual categories tested (bodies, faces, places, and tools), and provide evidence that the human brain’s intrinsic network organization plays a prominent role in the generation of functionally relevant, localized responses.

**Author Summary:** A fundamental question in neuroscience has persisted for over a century: to what extent do distributed processes drive brain function? The existence of category-selective regions within visual cortex provides long-standing evidence supporting localized computations, wherein specialized functions (e.g., selective responsiveness to face images) are thought to be primarily generated by within-region processes. This account was recently updated to include category selectivity dispersed across visual cortex, in the absence of category-selective regions. Here we provide groundwork evidence demonstrating that locally-exhibited visual-category-selective responses can be accurately generated via distributed activity flowing over globally connected systems. These processes were simulated via empirically-based computational models initialized by stimulus-evoked activity patterns and empirical connectivity matching each category-selective region’s unique intrinsic functional connectivity fingerprint. Results demonstrate that activity flowing over the human brain’s distributed network architecture can account for the generation of category selectivity in visual cortex regions.

## Introduction

Fundamental progress in cognitive neuroscience will require understanding how functionally-relevant, localized activations are established in the distributed and networked brain systems within which they are embedded. Most modern neuroscience assumes that both local (i.e., spatially restricted) and distributed (i.e., spatially dispersed across connected systems) processes coexist in the brain [1], but the extent to which (distributed) brain connectivity patterns shape localized activations remains unclear. There is extensive evidence for specific cortical regions exhibiting highly selective responses for visual categories such as bodies [2,3], faces [4], places [5], and tools [6]. For instance, selectivity for face images in the fusiform face area (FFA) has been validated with human neuroimaging [7], neural recordings in non-human primates [8], human lesion studies [9], and brain stimulation in humans [10]. In parallel, a landmark study provided evidence that visual categories could be decoded from dispersed and overlapping activity patterns in ventral temporal cortex [11]. Contemporaneous work proposed that a cortical region’s functioning is largely determined by its connectivity fingerprint (not to be confused with individual difference “connectome fingerprinting” [12]) – a region’s pattern of connections that specify what information that region receives and conveys [13]. Additionally, prior work has shown that a brain region’s response can be predicted by a model of its connective field, given by spatially-modeled (Gaussian) activity patterns in other parts of the brain [14]. The key hypothesis that this collective history of research suggests is that connectivity fingerprints are not just a byproduct of a region’s function, but a key determinant [15].

Over recent decades, the functional relevance of brain network interactions has been borne out by a growing body of network neuroscience [16] studies assessing structural, functional, and effective connections [17,18] during rest and task [19]. A core theoretical position across network neuroscience is that neurocognitive information is propagated across structurally and/or functionally organized brain systems. Building on this, we propose that whilst propagating, information is transformed not only by local computations, but by features of connectomic organization as well. In this framework, brain systems do not solely act as routes along which locally-computed information can be conveyed, but as additional means of shaping localized activations along the way [20]. Information processing is thus a complex operation that is at least partially specified by brain network interaction patterns. The critical gap in the literature is systematically quantifying the extent that network interactions specify or account for functionally-relevant processes during tasks. Supporting a crucial role for connectivity fingerprints in visual processing, work by Osher et al. [21] found that structural connectivity fingerprints can predict visual category responses in select ventral temporal cortex regions. However, because statistical associations were used to make those predictions (rather than a generative and/or mechanistic model), it remains unknown why connectivity is predictive of visual category selectivity. Moreover, it is unclear whether the brain’s intrinsic (i.e., resting-state [22–25]) functional network architecture specifies the interactions required to support such a mechanism. Thus, building on work by Osher et al. [21] (as well as [26]), quantifying the extent that intrinsic functional network interactions shape visual category selectivity is a crucial next step in understanding how function-relevant, localized activations are generated in the human brain.

Our recent work demonstrated the importance of activity flow processes – the movement of activity between neural populations – in generating neurocognitive processes [27,28]. The activity flow mapping approach can be conceptualized as generating computational models from empirical connectivity data that are then tested empirically using task-evoked activations [20]. This computational modeling approach is unique in its contact with empirical data, allowing especially rigorous conclusions regarding the empirical validity of hypothesized model features in generating cognitive processes (e.g., visual category selectivity). Specifically, activity flow mapping makes contact with empirical data in three ways: (1) empirical brain connectivity is used to build an activity flow model, (2) inputs to the model are empirical brain activations in “source” (e.g., sensory input) regions, and (3) model-generated cognitive activations are tested as predicted mappings of “target” empirical brain activations. The first two steps keep the proposed activity flow mechanisms close to empirical reality, while the last step empirically tests the validity and causal sufficiency of the proposed activity flow mechanisms for the generation of the cognitive processes of interest.

In the present study, we hypothesized that visual category selectivity exhibited by visual cortex regions is generated by task-evoked activity flowing over each of these region’s unique connectivity fingerprints. By systematically testing several models of such activity flow processes (Fig 1), we establish the extent that activity flowing over the brain’s distributed resting-state network architecture can account for localized visual category selectivity. Comparing process models in this manner not only provides a quantitative estimate for how well distal activity flowing over connectivity patterns account for task-evoked activations (Fig 1A), but also the impact of known visual system features (Fig 1B-D) amidst the overall process. Importantly, this supports a framework for uncovering the functional importance of both localized and distributed processes, concurrently. We propose that distributed activity, selected by a given region’s connectivity fingerprint, converges on that region, conferring it with locally specialized functioning (e.g., category selectivity).

**Fig 1.**
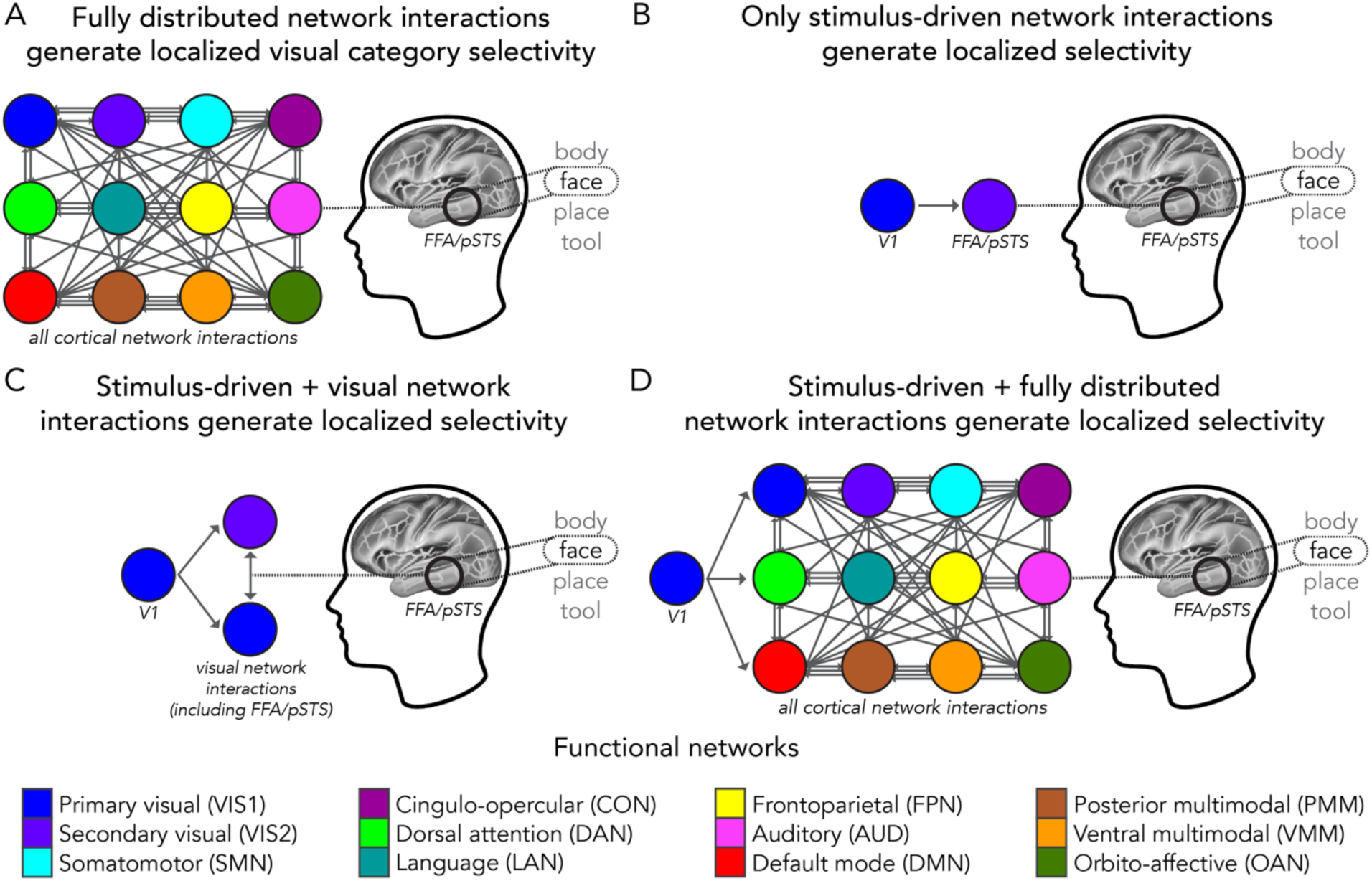
Network interaction models that could generate localized visual category selectivity. In all panels, a simplified model schematic of face selectivity in the fusiform face area and posterior superior temporal sulcus (FFA/pSTS, Methods) is depicted as a localized outcome of network interactions. We tested these models by applying activity flow mapping from Cole et al. [27,28] to select held-out “target” regions (including FFA/pSTS). Circles: regions in a network (color legend: bottom [25]). Gray arrows: activity flow processes (Methods), which are weighted (by each region’s connectivity fingerprint) but shown as uniform for visualization purposes. (**A**) This model predicts that all cortical network interactions (“fully distributed”) converge to generate localized face selectivity. (**B**) This model predicts that only stimulus-driven interactions with V1 generate face selectivity. We tested this by restricting activity flow mapping initialization to vertex-level data from V1. (**C**) Extending panel B: This model predicts that stimulus-driven interactions are further shaped by activity flow processes in visual networks. We tested this by simulating the flow of V1-initialized stimulus-evoked activity through the entire visual system. (**D**) This model predicts that fully distributed interactions (i.e., beyond just the visual system in panel C) are important for generating localized visual selectivity. We tested this by simulating the flow of V1-initialized stimulus-evoked activity through the entire set of cortical brain regions, including recurrent/feedback activity flow. Using mappings initialized in V1 (as in B-C), we assessed how well selectivity was generated when the fully distributed set of network interactions was initialized by stimulus-driven activity flow processes.

One likely purpose of this localized convergence (e.g., in the FFA) of network interactions is to represent related information (e.g., faces) distributed throughout the brain in a common space for efficient competition between alternative activity patterns. In this account, FFA may facilitate face recognition via implementing a winner-take-all computation (via lateral inhibition; [29]) between alternate faces represented in FFA, which is consistent with many computational models of local neural functions [30]. Importantly, we propose that the selective activity observed in functionally specialized regions is generated primarily by distributed activity flowing over each region’s connectivity fingerprint. Our hypotheses suggest that connectivity patterns play an active, generative role in localized region-level activity, and predict that estimating activity flow over a region’s unique connectivity fingerprint would account for the majority of that region’s localized task-evoked selectivity. Importantly, evidence that distributed activity flow processes provide the dominant influence on visual-category-selective responses does not discount the importance of local computations, but rather provides the groundwork for understanding how network interactions can sufficiently generate localized activations that may be mistaken for local computations.

Activity flow mapping [27,28] is well-suited to test these hypotheses (relative to other approaches, such as standard encoding models and artificial neural networks) because all parameters are explicitly based on empirical quantities, rather than abstractions [31,20]. Brain connectivity parameters (i.e., a region’s unique connectivity fingerprint) are key to characterizing the network-based processes that converge to generate localized responses. Moreover, we explicitly test whether activity flow mapping based on intrinsic connectivity fingerprints can generate localized visual category selectivity, which provides novel evidence for the functional relevance of the brain’s resting-state network architecture within the context of well-established localized visual responses.

We used resting-state functional connections within our models based on extensive evidence that they are highly similar to functional connections during a wide variety of task states [32–34]. Estimating a single set of connection weights for our models, rather than a separate set of connection weights for each task condition, allowed us to use a parsimonious set of general (as opposed to stimulus-specific) visual processing models. The intrinsic (i.e., state-general) nature of resting-state functional connectivity partially reflects constraints from structural connectivity on resting-state functional connections [35]. Importantly, our use of multiple regression (rather than the more standard Pearson correlation) to estimate functional connectivity is known to make the relationship to underlying structural connectivity stronger [36,37]. This is due to multiple regression fitting all time-series simultaneously, substantially reducing confounded and indirect functional connections to more closely reflect structural connections [27,38]. Together, these prior studies suggest our use of resting-state functional connectivity estimated using multiple regression resulted in highly stable estimates of underlying connectivity that generalize well to a variety of brain states (e.g., all visual processing conditions included here).

We tested our hypotheses (Fig 1) — an initial step toward understanding how localized activations emerge from distributed network interactions in the scope of visual category selectivity — utilizing human functional magnetic resonance imaging (fMRI) resting- and task-state data from the Human Connectome Project (HCP) [39] young adult release. We focused on category-specific representations of bodies, faces, places, and tools embedded in a working memory task. These visual categories correspond to functionally specialized visual cortex regions that are highly replicated in the literature, such as the fusiform face area (FFA) and the parahippocampal place area (PPA). In sets (or complexes) of functional ROIs in the visual cortex, we quantified actual and activity-flow-mapped visual category selectivity and estimated the degree to which fully distributed (i.e., whole-cortex) network interactions (Fig 1A) contributed to this selectivity. We probed the differential contributions of large-scale functional networks to activity flow processes and used substitution null models to corroborate that a given ROI’s connectivity fingerprint significantly shaped its visual category selectivity. Given that visual input from the retina is known to initially arrive (via the lateral geniculate nucleus) in cortex primarily via V1, along with evidence that fMRI activity in V1 represents a retinotopic map of visual features [40,41], we also tested refined hypotheses (Fig 1B-1D) that specify stimulus-driven activity flow processes as initialized in V1. In adapting the activity flow mapping procedure to address stimulus-driven hypotheses, we were also able to reduce the potential impact of causal circularity by constraining empirical parameters to network interactions originating in V1 only. Systematically testing how well visual category selectivity can be generated via empirically-constrained activity flow mapping constitutes an important first step in developing a network-interaction-based framework for how localized functionality emerges within cortex.

## Materials and methods

### Data acquisition and participants

Data used in the present study were collected by the Washington University-Minnesota Consortium of the Human Connectome Project (HCP) [39] as part of the young adult (S1200) release. Participants all provided written informed consent in accordance with protocols approved by the ethics review board of Washington University. This dataset included resting- and task-state functional neuroimaging data (see following Methods sections for more details on each). We obtained the minimally preprocessed [42] neuroimaging data for N=352 healthy young adults. Following the pipeline outlined Ito et al. [43], this N=352 subset of the HCP participants were selected to ensure high quality scan data and no familial relations. All study procedures were approved by the Institutional Review Board of Rutgers University-Newark. Further details on participant recruitment can be found in Van Essen et al. [39].

We used a split-sample validation approach to minimize false discovery rate [44]. The cohort of HCP participants in the present study (N=352) were quasi-randomly allocated to either a discovery (n=176) or replication (n=176) dataset. Participants were chosen randomly, but the sample size of both discovery and replication datasets were set at n=176 to ensure equal split-halves. Each dataset was analyzed identically but independently. Participants in the discovery dataset (77 men, 99 women) had the following age ranges: 22-25 years (26.14%), 26-30 years (41.48%), 31-35 years (30.68%), and 36+ years (1.7%). Participants in the replication dataset (93 men, 83 women) had the following age ranges: 22-25 years (22.16%), 26-30 years (43.18%), 31-35 years (34.09%), and 36+ years (0.57%). Results presented in figures refer to the discovery dataset, with replication dataset results reported in-text.

### Task paradigm

In a subset of analyses (see “Network analyses: Response profile network contributions”), we used data corresponding to seven HCP tasks and 24 task conditions that sampled diverse cognitive domains: emotion, gambling reward, language, motor, relational reasoning, social cognition, and working memory. In all other analyses we focused on the working memory/category-specific representation n-back task (see [45] for more details on all tasks). Prior studies suggest that the n-back task can be used to test hypotheses regarding the function(s) of specific brain areas [46]. Further, manipulating how far back a participant must remember (0-back versus 2-back) allows for the assessment of working memory maintenance (along with other contrasts, such as stimulus type). While working memory is not the main interest of the present study, specifically manipulating working memory has the added benefit of promoting task engagement.

The cognitive domain of interest in the present study was the processing of semantic categories [11,47,48]. Category-specific representations were embedded in the n-back task via blocks of trials that presented images of four reliably studied and distinct visual semantic categories [6,21]: (1) bodies [2,3]; (2) faces [4,49,50] ; (3) places [5,51]; and (4) tools [47,52,53].

The specifics of the n-back task are detailed in Barch et al. [45]. Briefly, the n-back task included 2 runs with blocks of visual category trials (8 blocks, 10 trials each, 2.5 seconds each) and fixation blocks (4 total, 15 seconds each). Half of the blocks were 0-back and half were 2-back. During 0-back working memory tasks, participants determined via button-press whether presented stimuli (images of either bodies, faces, places, or tools) matched a constant target shown in the initial cue screen. During 2-back working memory tasks, participants were asked to indicate if the present stimulus matched a target from two screens back. Given that working memory load was not the main interest of the present study, results presented here reflect an average across 0-back and 2-back conditions (i.e., fMRI data was averaged after all preprocessing steps), except for in analyses examining “response profiles” (i.e., responsiveness of regions-of-interest to 24 HCP conditions, assessed independently). Additionally, preliminary analyses found no evidence that visual category selectivity was variable across the two working memory load conditions.

### MRI parameters

All MRI data were collected at Washington University in St. Louis, Missouri, USA in a customized Siemens 3T “Connectome Skyra” scanner. Whole-brain, multiband, and echo-planar images (EPI) were acquired with a 32-channel head coil. The repetition time (TR) was 720 ms; the echo time (TE) was 33.1 ms; the flip angle was 52 degrees; the bandwidth (BW) was 2290 Hz/Px; in-plane field-of-view (FoV) was 208 x 180 mm; 72 slices; 2.0 mm isotropic voxels; and the multiband acceleration factor was 8. Whole-brain and high-resolution T1-weighted and T2-weighted anatomical scans were also acquired, with an isotropic voxel resolution of 0.7 mm. All imaging data were collected over two days for each participant. Each day included two resting-state fMRI scans, each lasting 14.4 minutes (for a total of approximately 29 minutes of rest data per day). During rest, participants were instructed to keep their eyes open and fixated. Task fMRI (30 minutes of task data per day) followed resting-state fMRI scans on each day. Each of the 7 tasks were performed over two consecutive runs. Further details on MRI protocols and parameters can be found in Van Essen et al. [39].

### fMRI preprocessing

We acquired minimally preprocessed data [42] from the HCP database (www.humanconnectome.org). The minimal preprocessing pipeline is extensively detailed in Glasser et al. [42] and is openly available (https://github.com/Washington-University/HCPpipelines). In-brief, minimal preprocessing included: anatomic reconstruction and segmentation, motion correction, intensity normalization, and EPI reconstruction to a standard (surface-based) template. We also implemented HCP MSM-All registration (multimodal surface matching registration based on areal features from multiple sources, including: cortical folding/sulcal depth, myelin maps, resting-state networks, and visuotopic maps; [54]). Preprocessing was performed equivalently but independently on all functional runs, including resting-states and the 24 task-state conditions. The resulting data were in CIFTI grayordinate (i.e., vertex) space. We performed additional preprocessing steps on cortical vertices (59,412 vertices across the two hemispheres). On both resting- and task-state data, we removed the first five frames of each fMRI run, demeaned and detrended the timeseries, and performed nuisance regression. We followed a variant [43] on methods suggested by Ciric et al. [55] to model and regress out nuisance parameters, including motion and physiological noise. We performed nuisance regression to remove six motion parameters, their derivatives, and quadratics (24 total). Anatomical CompCor (aCompCor; [56]) was used on white matter and ventricle timeseries to model physiological noise. The first five principal components from white matter and ventricles were independently extracted. These parameters, their derivatives, and the quadratics of all regressors were removed (40 total). Altogether, 64 nuisance parameters were modeled, and associated variance removed from the data.

Global signal was not removed given evidence that it can artificially introduce negative relationships [57,58] and is non-standard for task activation analyses. However, the nuisance parameters modeled and removed with aCompCor contain components similar to the proposed global signal, with the added benefit of not removing gray matter signals [43,59].

### Task-state activity estimation

Cortical activations related to task-state fMRI conditions (n-back visual semantic categories) were estimated by a standard general linear model (GLM), which convolved task timing (per block-design condition) with the SPM canonical hemodynamic response function [60]. This was performed per cortical vertex, each belonging to a parcel (or region) of the Glasser et al. [54] multimodal parcellation (MMP) atlas. Average coefficients of the GLMs were utilized for each region or functional complex (described further below in “Identification of functional complexes”) as estimates of visual category evoked brain activity. Note that the fully-distributed model (Fig. 1A) was performed on the region-level data; the stimulus-driven (Fig. 1B) and stimulus-driven plus visual network interactions (Fig. 1C) models were performed on vertex-level data before averaging into functional complexes; and the stimulus-driven plus fully-distributed model (Fig. 1D) was performed on vertex-initialized data in the first step and region-level data in the second step. The steps and rationale of each of these models are extensively detailed below.

### Identification of functional complexes

To address the question of whether distributed network interactions can map [19,27,28] localized, category-selective functional responses, we focused on the functional brain regions identified in Osher et al. [21] given the similarity of the four visual semantic categories in that study to the HCP n-back task studied herein. Additionally, as suggested by Poldrack et al. [61], variables that will be used in predictive analyses should not be chosen in an empirically-driven manner (e.g., from post hoc cross validation and/or optimization procedures) because this has the potential to artificially inflate accuracy due to circularity. Thus, selecting functional regions of interest a priori has the benefit of controlling false positives in accuracy, particularly with our focus on the well-established visual category literature. Upon further review of the literature (at the time of originally conducting this study and compiling this manuscript: 2021-2022) as well as the regions belonging to the Glasser et al. [54] MMP atlas, we observed that these functional regions: (1) contained multiple MMP regions (e.g., the parahippocampal place area or PPA, which contained 6 MMP regions as in [51]); (2) did not have an exact terminological match in the MMP atlas; and/or (3) included two discrete locations (e.g., the retrosplenial complex, or RSC, often studied in conjunction with the PPA). To address this we first performed a literature search of each of the Osher et al. [21] functional regions with Google scholar (https://scholar.google.com/) and Neurosynth (https://neurosynth.org/), filtering for relevance, year, and citation count (when possible) (see Table 1 references in right-most column). We focused further on publications that had corresponding volumetric coordinates (or MMP surface region labels when possible) and generated a list of Montreal Neurological Institute (MNI) x/y/z volumetric coordinates for each functional region (with a minimum of 10 entries per region). We then used the average x/y/z coordinate set as a reference to identify corresponding MMP regions (Table 2).

**Table 1.** Specifications of functional brain areas.

| Visual category | Region (acronym, L/R indices) | Network | Functional complex | References |
| --- | --- | --- | --- | --- |
| Bodies |  |  |  |  |
|  | MST, 2, 182 | VIS2 | EBA | [2, 3, 45, 62-65*, 66-72] |
|  | PH, 138, 318 | VIS2 | EBA | [2, 3, 45, 62-72, 73*, 74*] |
|  | V4t, 156, 336 | VIS2 | EBA | [2, 3, 45, 62-63, 64*, 65*, 66-72, 75*] |
|  | FST, 157, 337 | VIS2 | EBA | [2, 3, 45, 62-63, 64*, 65*, 66-72] |
|  | TE2p, 136, 316 | DAN | FBA | [3, 67, 70-72, 73*, 74*, 76-81] |
| Faces |  |  |  |  |
|  | FFC, 18, 198 | VIS2 | FFA | [4*, 45, 66, 67, 80, 82-84*, 85-92] |
|  | STSdp, 129, 309 | LAN | pSTS | [69-71, 81, 87, 88, 90-96] |
|  | STSvp, 130, 310 | DMN | pSTS | [69-71, 81, 87, 88, 90-96] |
| Places |  |  |  |  |
|  | PHA1, 126, 306 | DMN | PPA | [5, 45, 66, 67, 97-104] |
|  | PHA2, 155, 335 | DMN | PPA | [5, 45, 66, 67, 97-104] |
|  | PHA3, 127, 307 | DAN | PPA | [5, 45, 66, 67, 97-104] |
|  | VMV1, 153, 333 | VIS2 | PPA | [5, 45, 66, 67, 97-104, 105*, 106] |
|  | VMV2, 160, 340 | VIS2 | PPA | [5, 45, 66, 67, 97-104, 105*, 106*] |
|  | VMV3, 154, 334 | VIS2 | PPA | [5, 41*, 45, 66, 67, 97-104] |
|  | POS1, 31, 211 | DMN | RSC | [5, 82*, 97, 98, 101, 102, 104, 107-109] |
| Tools |  |  |  |  |
|  | V4, 6, 186 | VIS2 | LOC | [65*, 67, 72, 86, 99, 110, 111*, 112-122] |
|  | V8, 7, 187 | VIS2 | LOC | [65*, 67, 72, 86, 99, 110-122, 123*] |
|  | LO1, 20, 200 | VIS2 | LOC | [65*, 67, 72, 75*, 86, 99, 110, 111*, 112-122] |
|  | LO2, 21, 201 | VIS2 | LOC | [65*, 67, 72, 75*, 86, 99, 110, 111*, 112-122] |
|  | PIT, 22, 202 | VIS2 | LOC | [8*, 11*, 64*, 65*, 67, 72, 86, 99, 110-122] |
|  | V3CD, 158, 338 | VIS2 | LOC | [65*, 67, 72, 86, 99, 110-122] |

**Table 2.** Average MNI coordinates for each functional complex from the literature.

| Functional Complex | Left hemisphere |  |  | Right hemisphere |  |  |
| --- | --- | --- | --- | --- | --- | --- |
|  | x<br>mean (SD) | y<br>mean (SD) | z<br>mean (SD) | x<br>mean (SD) | y<br>mean (SD) | z<br>mean (SD) |
| EBA | -48.9 (6.6) | -73 (4.6) | 2.7 (8.3) | 45.7 (6.5) | -69.5 (3.5) | -0.7 (3.9) |
| FBA | -39.7 (4) | -44.2 (8.2) | -20.2 (1.6) | 41.9 (3.5) | -47 (6.2) | -21.5 (3.7) |
| FFA | -41.2 (3.6) | -53.6 (7.2) | -18.6 (3.9) | 40.6 (3.5) | -51.7 (3.6) | -18.6 (5.2) |
| pSTS | -51.8 (6.5) | -49.9 (9.9) | 8 (6) | 51.3 (4.1) | -49 (8.1) | 8.6 (4.4) |
| PPA | -27.2 (4.9) | -44.6 (6.2) | -9 (4.7) | 25.6 (4.4) | -42.4 (5.5) | -10.3 (5.1) |
| RSC | -13.3 (4.2) | -56.5 (6) | 10 (6.2) | 14.3 (5) | -54 (5.6) | 9.3 (7.8) |
| LOC | -45.7 (7.1) | -70.4 (8.5) | -8.5 (6.3) | 42.5 (6.4) | -69.4 (11.1) | -9 (7.2) |

To convert from volumetric space to region space, we employed the following steps (using Connectome workbench, FreeSurfer, and AFNI/SUMA commands, per hemisphere): (1) convert the MMP atlas to FreeSurfer’s fsaverage space; (2) convert fsaverage to the surface-mesh MNI-152 template (which is also included in AFNI); (3) convert surface MNI-152 template to volumetric MNI-template and obtain coordinates. Additional commands were used to ensure that the templates aligned as best as possible, including zero-padding, thicker ribbons (that were later filled), and adding MMP sphere labels. To obtain x/y/z coordinates, we divided the template into equal portions perpendicular to the long axis of a given sphere label, identified the middle division, then calculated the centroid. We then found the MMP regions that were physically closest to the reference coordinate set for each functional region. We then referred to the Glasser et al. [54] supplemental results (as well as publications they based the MMP atlas upon; see references with asterisks in Table 1) to exclude regions that were physically close but functionally irrelevant (e.g., auditory regions near the pSTS, which was a potential ROI for face processing).

The final set of regions, including their proposed category selectivity, network affiliations, and references are listed in Table 1. The term “functional complex” indicates firstly that more than one MMP region comprised each complex, and secondly serves to distinguish an MMP atlas region from a category-selective (based on prior literature) complex in the present study. Further, in some cases there were two functionally relevant complexes in the literature (e.g., PPA and RSC, both relevant to place processing, as in [21]), therefore both were included. When appropriate, we applied the Cole-Anticevic brain-wide network partition (CAB-NP, [25], Fig 3C), which was based on HCP resting-state fMRI data and was generated from a Louvain community detection algorithm that assigned MMP regions to 12 functional networks (labeled via distinct colors in Fig 3C). Table 1 lists the network assignments of each region, which is further reflected in Results figures by color labeling matching Fig 3C. The reference sets of volumetric coordinates (x/y/z) are listed in Table 2.

### Resting-state functional connectivity estimation

Functional connectivity (FC) was modeled as a series of target dependent variables (i.e., each brain region), each with multiple source independent variables (i.e., other brain regions) (see following section “Activity flow mapping: a generative model”). As in Cole et al. [27], source vertices excluded all vertices within a 10 mm dilated mask of the target region (as well as excluding vertices of the target region itself). This reduced potential circularity related to spatial autocorrelation by avoiding nearby vertices in predicting the activity of the target. Modified source sets of vertex-level timeseries were then averaged together based on their Glasser et al. [54] MMP region assignment. FC was then estimated with a method proposed by Sanchez-Romero and Cole [125] termed combinedFC. CombinedFC integrates the standard methods of bivariate correlation, partial correlation, and multiple regression together to account for causal confounds better than each method alone (as in Results Fig 3A). In Sanchez-Romero and Cole [125], combinedFC improved precision in a series of simulated networks, and retained robust network structure with empirical fMRI data (see [125] for a detailed report of this method). Briefly, each participant’s resting-state data (pre-processed) was firstly assessed with partial correlation (amongst each pair of regions, conditioning on all other regions) to find conditional associations, which were examined for statistical significance. This step addressed potential confounders and causal chains [126] and built an initial connectivity architecture by adding in significantly correlated edges only. Then, bivariate correlation estimates (i.e., Pearson correlation) for each pair of connected regions (from the initial partial correlation step) were used to remove spurious edges resulting from conditioning on colliders (specifically, edges where r was statistically equal to zero). Lastly, a multiple regression procedure was performed to scale the estimated edge weights. For each target region functioning as a response variable, all other connected source regions functioned as regressors to estimate the final combinedFC weight.

### Activity flow mapping: a generative model

Activity flow mapping is a generative modeling approach that maps held-out brain activity based upon the inherent link between activation patterns and functional connectivity [20,32,127,128]] (i.e., an empirically-parameterized extension of generative network models outlined in Betzel and Bassett [129]). Activity flow was developed by Cole et al. [27] (see Cocuzza et al. [28] for further details), and is described by Formula 1 (Fig 2):

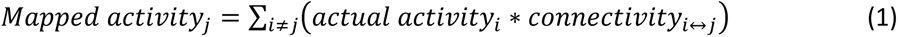

**Fig 2.**
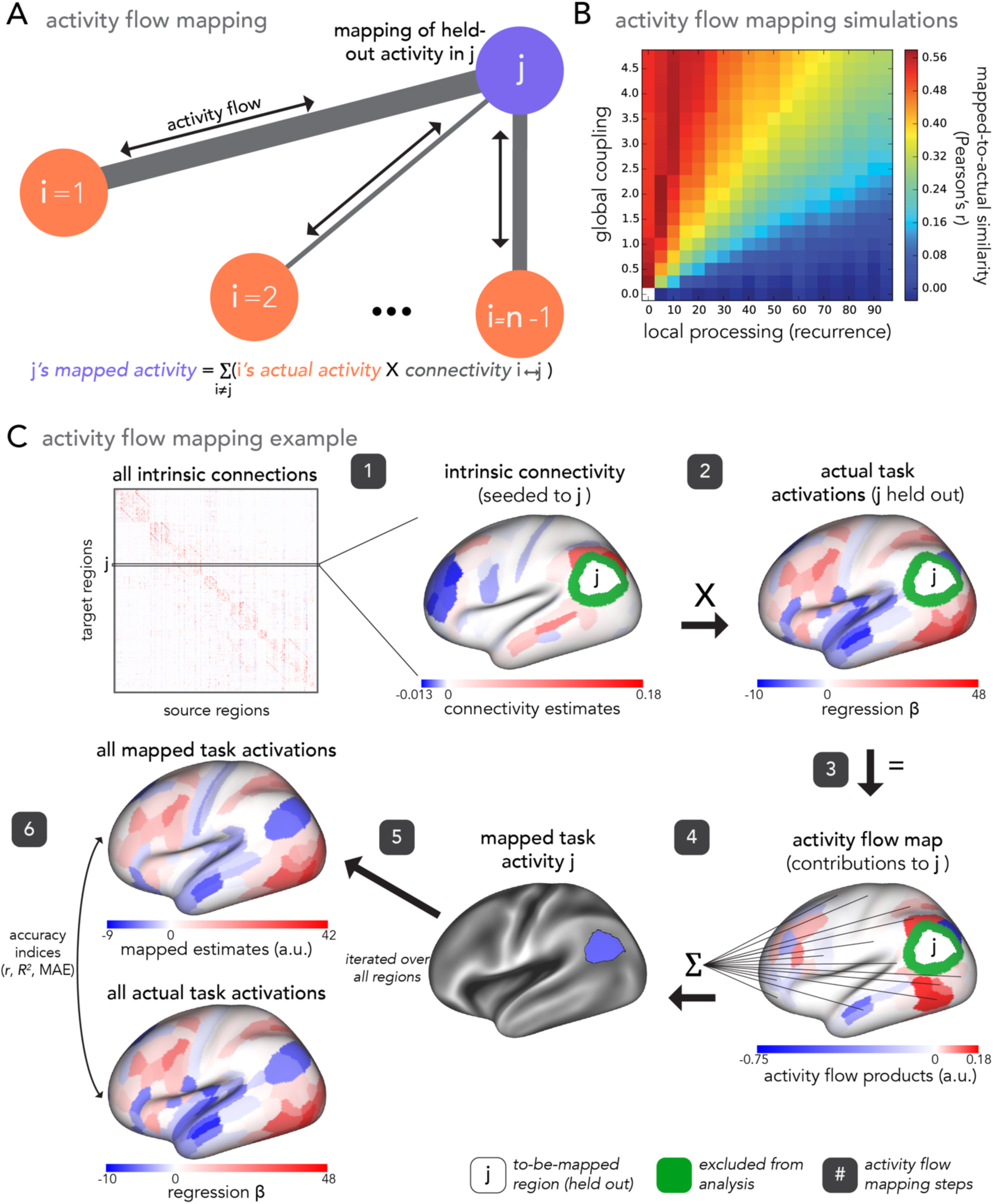
Activity flow procedure to map task activity in held out brain regions. **(A**) Activity flow mapping toy diagram and formula. Task activity for held-out region *j* (purple) is mapped as the sum of task activity of all other cortical regions, *i* (coral) (*n* = total number of regions), weighted by their connectivity estimates with *j* (gray). (**B**) Activity flow simulation results (reproduced with permission from [27]) demonstrating that activity flow mapping is most successful when distributed processing mechanisms are high and localized processing mechanisms are low. (**C**) Example of activity flow mapping with empirical data (steps 1-6) (reproduced with permission from [28]). For a given target region *j*, estimates of intrinsic (e.g., resting-state) connectivity between *j* and all other source regions (step 1) are multiplied by all other regions’ actual task activations (step 2). This yields an activity flows map quantifying the contribution of all other regions’ activity flow upon the held-out region, *j*, for the task of interest (step 3). These are summed to equal the mapped task activity of *j* (step 4). This procedure is iterated over all regions to generate activity-flow-mapped task activations across the whole cortex (step 5). This is compared with the actual whole-cortex map of task activations via Pearson’s *r*, MAE, and R^2^ to estimate accuracy (step 6). Importantly, this approach is flexible to different estimates of connectivity. Source vertices not included in the analyses (10 mm from the target region *j*; see Methods) are depicted in green.

Where j is a held-out brain region, and i indexes all other regions in a given parcellation. Note that the standard application of activity flow mapping given by Formula 1 is consistent with the “fully distributed” mapping of Fig 1A (i.e., mapping that is used to generate visual category selectivity, see “Category selectivity” below) because source regions (i) refer to all other cortical regions. Variations on this are outlined in later Methods sections (see the following sections below titled “Stimulus-driven visual category selectivity” and “Category selectivity from stimulus-driven network interactions further shaped by fully distributed network interactions”). This method induces the bias that brain activations are propagated and shaped via distributed network interactions. Throughout the present study, the term “distributed” means globally connected brain systems that are specified by functional brain network organization (i.e., functional connectomes). Applied to research questions on functional processing, activity flow maps task-evoked (functionally relevant) activity in held-out target regions as the sum of the task-evoked activity in all other source regions scaled by FC estimates between the target region and corresponding source regions (Formula 1). The activity flow mapping procedure explicitly tests the hypothesis that the sum of empirically-based source region activity, parameterized by empirical patterns of brain connectivity, is equal to target region activity. Using a simulated model of neural processing, Cole et al. [27] found that activity flow mapping was most accurate when a global coupling parameter was high and the strength of self-coupling (local) connections was low (Fig 2B). Critically, this suggests that the successful application of fully distributed activity flow mapping (Fig 1A) to empirical brain data reflects similar high global coupling properties being preserved in real (rather than simulated) data. Thus, the success of activity flow mapping can be taken as a proxy for how much signals over distributed network interactions contribute to a given activation (see “Category selectivity” below for more details).

To map category-selective responses to bodies, faces, places, and tools we applied the activity flow approach to each of these categories independently. Actual task activations used were beta estimates generated by task GLMs (described above in “Task-state activity estimation”), with a focus on the regions comprising functional complexes as the held-out target regions (described above in “Identification of functional complexes”). Functional connectivity estimates between targets (functional complex regions) and sources (all other connected regions) were based upon resting-state timeseries and estimated via combinedFC (described above in “Resting-state functional connectivity”). To assess accuracy, activity-flow-mapped category responses were compared to actual (i.e., empirical) category responses via Pearson r correlation coefficient (NumPy corrcoef function in Python), mean absolute error (MAE), and the coefficient of determination (R^2^) (sklearn r2_score function in Python).

Each of these indices quantified a different aspect of accuracy, altogether providing a thorough evaluation of activity flow mapping. In brief, Pearson’s r estimated the linear correlation between actual and mapped brain activations, with a potential range from -1 to 1 and the property of scale invariance. MAE measured the arithmetic mean of absolute errors between actual and mapped brain activations, with a sensitivity to scale. The coefficient of determination (R^2^) estimated the percent of unscaled variance in the actual brain activations that was accounted for in the mapped activations. Note that R^2^ maintains a range of negative infinity to 1; from 0 to 1 R^2^ can be interpreted as 0 to 100% variance, but negative values indicate incorrectly scaled predictions. Thus, each index had clear advantages and disadvantages, but altogether provided a complementary, full characterization of mapping accuracy. In all cases, these comparison statistics were taken per participant, then averaged across participants to provide a random effects estimate.

### Category selectivity

To assess how selective each functional complex of interest was for a given visual category, we measured the ratio of category activation amplitude divided by non-category activation amplitude, for both activity-flow-mapped data and actual data, per participant. For example, for the functional complex of the extrastriate body area (EBA) and fusiform body area (FBA) (see above: “Identification of functional complexes”), we quantified body responses divided by non-body responses. Importantly, non-category responses included only other n-back conditions, and not any other HCP task condition, such that the estimation of visual category selectivity only considers comparable task manipulations. Per participant and functional complex, we used Formula 2:

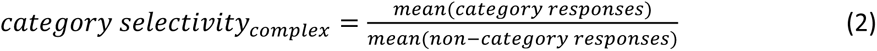

We will henceforth refer to the ratio in Formula 2 as “category selectivity”, which can be interpreted as how many times larger category responses were than non-category responses. For example, if a given participant’s body-category selectivity score (for the EBA and FBA complex) was 1.5, then their activations to body images were 1.5 times larger than their activations to non-body images (i.e., face, place, and tool images in the n-back task). A category selectivity score of 1.0 reflected the hypothesized null mean given that it quantifies category activations as equal to non-category activations. The main advantage of utilizing a ratio versus a difference score is that it is less sensitive to beta value scale differences across brain regions.

Prior to calculating the category selectivity score (Formula 2) it was necessary to normalize the range of activation values. This was necessary for two reasons: (1) avoiding negative numbers (which would limit interpretability of the selectivity values) and (2) rescaling activation values such that every participant’s data was in the same range (which allows for group level comparisons). We used a standard normalization procedure termed min-max normalization (also termed feature scaling). We used Formula 3:

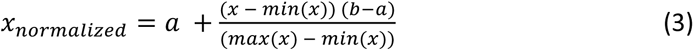

Where a and b are the lower and upper bounds of the rescaled data, respectively. We used a=0 and b=1.0, which were selected to increase interpretability of the category selectivity ratio since all ratios were positive and included a possible minimum of 0. The minimum and maximum values in Formula 3 were calculated based on the actual activations of a given region set (e.g., EBA and FBA across-complex average for body selectivity) across all 24 task conditions (x) for each participant separately. Note that results were highly similar if minimum and maximum values were calculated separately for mapped and actual activations. Additionally, we report the cross-complex average results herein given that no functional complex was substantially driven by a particular region (i.e., regions in a given functional complex had similar category selectivity scores). We additionally identified and removed outlier participants (in category selectivity) following the methods described by Leys et al. [130] (median absolute deviation), using a highly conservative threshold of -/+ 5 deviations relative to the median. Given this high threshold, outlier removal excluded a small number of participants (an average of n=5 of 176 were removed across analyses, or approximately 2.8% of participants) with category selectivity scores that were not representative of the group average, which is how category selectivity was reported (which had a large sample size of n=176 per discovery and replication datasets). Note that statistical significance (see “Experimental design and statistical analysis”) was not impacted by removal of outlier participants.

In assessments of the fully distributed network interaction scheme (Fig 1A), we estimated the contribution of distributed network interactions to a given category-selective response by computing the percent of actual category selectivity (distributed plus localized processes) captured by activity-flow-mapped category selectivity (i.e., a model based upon distributed processes only, Fig 2B). We propose that the percent leftover (i.e., not captured via activity flow mapping) is due to either local processes (not isolated in the model) or other sources of model error (e.g., noise in the data). We used Formula 4:

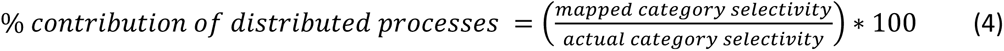

If the percentage given by Formula 4 was statistically greater than 50%, we inferred that category selectivity in that functional complex was generated primarily via distributed activity flow processes. Note that these estimates reflect a lower bound on the contribution of distributed processes, since they can be driven lower not only by local processes but by other sources of model error.

### Network analyses

In assessments of the fully distributed network interaction scheme of Fig 1A, we tested the contributions of each large-scale CAB-NP functional network ([25] Fig 3B) to activity-flow-mapped responses. We did this at both the levels of: (1) mapped category-specific responses, and (2) mapped response profile.

**Fig 3.**
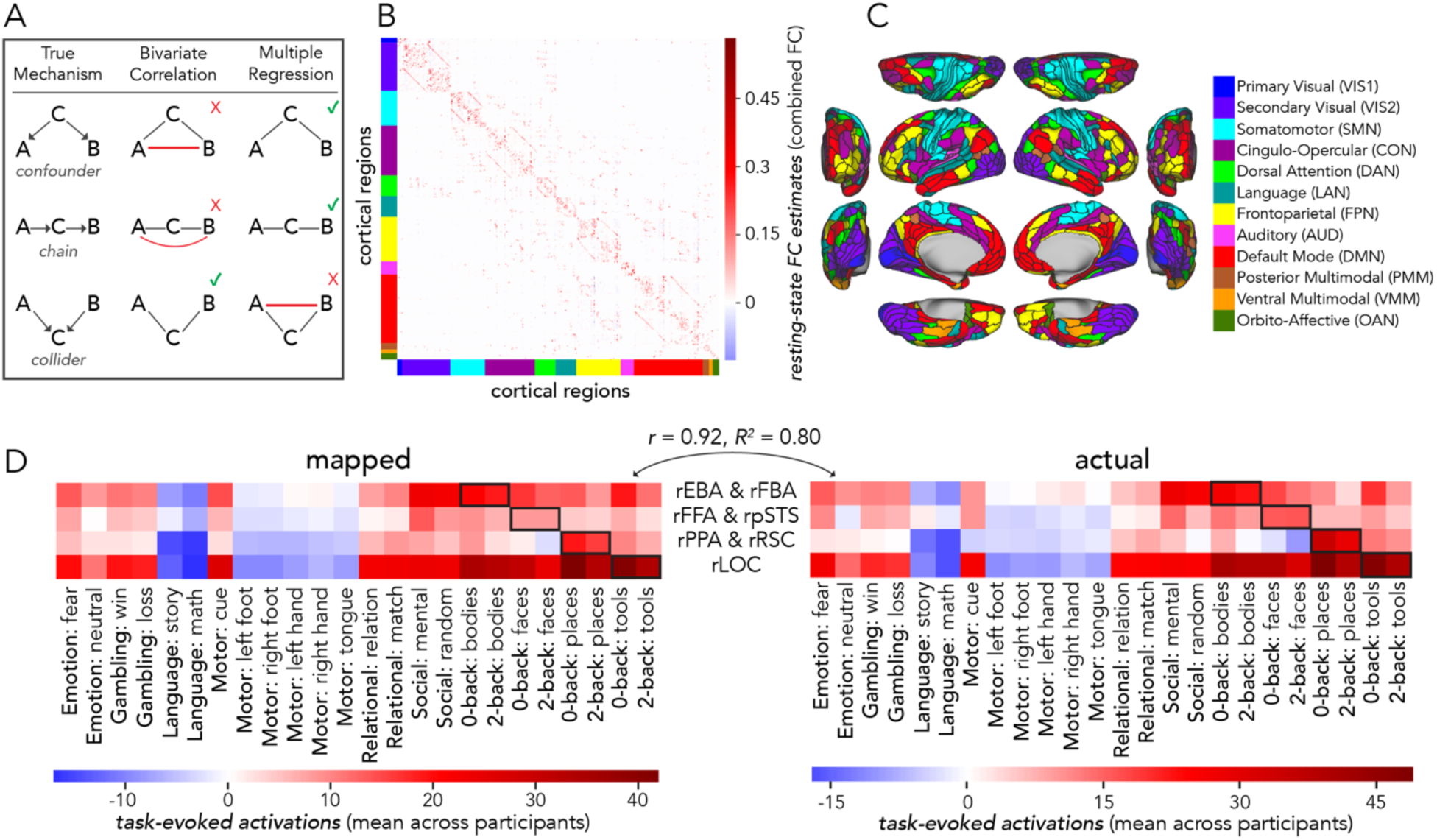
Mapped activations across all HCP conditions yielded response profiles for four functional complexes of interest. (**A**) Causally confounding graph patterns in standard FC estimation methods (adapted with permission from [125]). CombinedFC incorporates both bivariate and multiple regression measures such that confounders, chains, and colliders are accounted for. (**B**) The cross-participant (*n*=176) average resting-state connectivity matrix (estimated via combinedFC) of 360 MMP regions [54], ordered along each axis per the Cole Anticevic brain-wide network partition (CAB-NP [25]; color-coded on each axis to match panel C). This was the functional network organization utilized in the present study for activity flow mapping. Note that our implementation of combinedFC used multiple regression as the final step, and therefore FC estimates were given by beta coefficients (see Methods). (**C**) Cortical schematic of the CAB-NP and its 12 functional networks from Ji et al. [25], reproduced with permission. (**D**) Response profiles (across all 24 HCP conditions) of four complexes of interest to the present study (indicated along the y-axis; note that “r” stands for right hemisphere); mapped versus actual (left and right respectively; mean across participants depicted in each panel). Black boxes highlight the n-back conditions that maintained visual semantic category embeddings germane to a given functional complex (e.g., 0-back bodies and 2-back bodies for the right EBA and right FBA). Activity-flow-mapped response profiles were highly accurate, suggesting that mapped activation patterns of the functional complexes of interest were reliable across multiple cognitive domains.

#### Category-specific network contributions

The activity flow map (Fig 2C, step 3: “activity flow products”, or just “flows”) quantified how much activity of a given source region was weighted by functional connectivity with the held-out target region (i.e., before summing in Formula 1). We used the network-based averages of these flow values for each functional complex and its respective visual processing category to probe category-specific network contributions to the activity flow map. Source regions were averaged together based upon their CAB-NP assignments. These contributions were each statistically compared to one another (see “Experimental design and statistical analysis”) to discover the networks that contributed the most to a given activity flow map.

#### Response profile network contributions

To measure network contributions to activity flow mapping at the response profile level (i.e., across all 24 HCP conditions), we conducted a dominance analysis [131]. We first re-computed activity flow mapping (across all conditions for a given functional complex) with source regions iteratively restricted by CAB-NP network assignment. Note that these values exactly summed to the whole-cortex, activity flow mapping, thus this approach simply parsed a given mapping into its network components. Then, we conducted multiple linear regression (per participant, across conditions), where network-level mapped activations were the predictor variables (12 in total), and actual activity was the response variable. Following dominance analysis, this yielded the “full model” explained variance (R^2^), which in the present study refers to the proportion of variance in the actual response profile explained by the network-based, mapped response profile. An iterative subset of multiple regression models were then performed, with all combinations of predictors tested. These can be thought of as subset models because they were iterated on choosing one through eleven predictors, and performing multiple regression on all combinations therein (procedural details are openly available, and can be found here: github.com/dominance-analysis/dominance-analysis). This followed a standard combination calculation, nCr, where n was the full number of predictors, or 12, and r was iterated between one and eleven (note that r = 12 describes the full model). In the present study, there were a total of 4095 subsets of models for each functional complex assessed. The incremental contribution of each predictor to R^2^ was quantified for each of these subsets of models. Then, the cross-model average of these incremental contributions was taken as the partial R^2^ of each network, which added up to equal the full model R^2^. Given this property of dominance analysis (that the partial R^2^ values add up to the full R^2^), the percentage of relative importance of each network was calculated as the partial R^2^ divided by the full R^2^, times 100. The dominance analysis approach has the benefit of being robust to multicollinear predictor variables [132] (i.e., estimating unmixed partial R^2^).

### Control analyses

#### Assessing the impact of resting-state connectivity fingerprints

To assess how critical each functional complex’s unique connectivity fingerprint (as in [13,15]) was to respective visual category selectivity results, we performed an FC substitution experiment. For each target functional complex (and per participant), we substituted each complex’s connectivity fingerprint with each other functional complex’s fingerprint, then recomputed category selectivity (see above, “Category selectivity”). This meant that for each of the four original, or true, mappings (bodies, faces, places, and tools), there were three null model comparisons. For example: original body complex rsFC substituted with (1) face complex rsFC, (2) place complex rsFC, and (3) tool complex rsFC (repeated likewise for each original model). To preserve an essential property of the connectivity graphs used in the present study, we set self-connections to zero (e.g., right EBA/FBA target to right EBA/FBA source) (for more on constraint considerations for network null models, see [133,134]). Additionally, to account for the zero FC weights that were in the null functional complex’s self-connection locations, we set connectivity estimates between the original target complex and the null model complex with the transposed estimate, per hemisphere (e.g., right EBA/FBA target to right FFA/pSTS source set with right FFA/pSTS target to right EBA/FBA source). We performed paired samples t-testing to test the alternative hypothesis that the true mapped category selectivity was greater than the null mapped category selectivity (for each of the four visual categories and in each of the three respective FC substitution null models).

#### Assessing the impact of other category-responsive regions

We aimed to assess the potentially circular influence of regions excluded from functional complexes (see “Identification of functional complexes” above) that still exhibit responsiveness to visual categories in the literature. Such regions could, in principle, become the source of category selectivity for our target regions, rather than activity flows from non-category-selective regions. Note that the V1-initiated models (Fig. 1B-D; described in the next sections) were robust to this issue, given that all category selectivities had to originate (directly or indirectly) from activity flows originating in V1. The following regions were considered: (1) processing of body images: the dorsal visuomotor stream [135]; (2) processing of face images: the occipital face area [136–139], the inferior frontal gyrus [136,140,141], and the orbitofrontal cortex [142,143]; (3) place processing: the transverse occipital sulcus [144,145]; and (4) tool processing: the middle frontal gyrus [146,147] and the intraparietal sulcus [148,149]. In each case we converted MNI coordinates to surface space (as described in “Identification of functional complexes”) when available, or used topographic information provided by Glasser et al. [54] when regions were already identified within the MMP atlas (e.g., the orbitofrontal cortex). Note that an important exclusion criterion that removed these regions from the original definition of a functional complex was limited representation in the literature. Therefore, category selectivity in these regions is inherently less validated than the regions we did include. Regardless, category selectivity generated by the fully-distributed activity flow processes (Fig 1A) may still have been driven by these regions being part of the source region sets for corresponding functional complexes. For example: does the occipital face area drive activity-flow-mapped face selectivity in the FFA/pSTS? For each participant, functional complex, and hemisphere, we re-ran every step of the fully-distributed model with these regions held out from respective source sets. Then, we conducted an individual differences regression on the original fully-distributed category selectivity results against these source-set-limited category selectivity results. The more correspondence between these (measured by coefficient of determination, R^2^), the less likely held-out source regions were impacting the fully-distributed category selectivity estimates.

### Stimulus-driven visual category selectivity

We sought to test the possibility that stimulus-driven activity flow processes conferred by V1 are capable of generating visual category selectivity (Fig 1B). Note that analyses that examined stimulus-driven (described first below; Fig. 1B) and stimulus-driven plus visual network interactions (described second below; Fig. 1C) were performed with vertex-level data, then averaged into functional complexes after estimating category selectivity. The final model, stimulus-driven plus fully-distributed network interactions (Fig. 1D) were initialized with vertex-level data in a first step, then assessed across region-level averaged data, as detailed in the next section titled “Category selectivity generated by stimulus-driven network interactions further shaped by fully distributed network interactions”. Firstly, we estimated rsFC between all visual system (i.e., VIS1 and VIS2 network vertices from Ji et al. [25], see “Network analyses” above) vertices, and refer to this visual sub-system as “VIS” throughout corresponding Results and Figures. Note that we included all functional complexes in this VIS sub-system. FC was estimated with combinedFC [125] as in prior analyses (see “Resting-state functional connectivity estimation”), except regularized regression was used instead of multiple regression to scale edge weights, given that there were more vertices than time points in vertex-level resting-state data. For each participant and target vertex (i.e., dependent variable or vertex timeseries that connections were to be estimated with), we used L1 regularized lasso regression with cross-validation across resting-state runs. The penalty term was chosen from a wide sweep of 100 log-spaced terms. We then conducted a multi-step activity flow mapping procedure. We report step 1 as the V1, stimulus-driven model, and steps 2+ as the extended VIS model.

In step 1, only activation patterns of source vertices from V1 (both hemispheres), shaped by their connectivity fingerprints with each VIS vertex (see “Activity flow generative model” and Formula 1), were used to map the task-evoked activation patterns of each VIS vertex (Fig 1B). Resulting maps for functional complex vertices were the focus of the step 1 analysis, but the entire VIS sub-system was mapped because it was later used in steps 2+. Note that cortical region V1 was identified based on the V1 region identified by Glasser et al. [54] using multimodal methods. Cortical V1 vertices were all those inside of the V1 border, with 10 mm dilated masking applied to activity flow mapping (with VIS) as described previously (see “Resting state functional connectivity estimation” and “Activity flow generative model”, Fig 2) to control for the effects of spatial autocorrelation. When mapping held-out target vertices of each functional complex, the cross-vertex (i.e., across V1) mean activation value for the corresponding category-specific condition was subtracted from each V1 source vertex to account for the (unlikely but possible) potential confound that there may still be category-specific information fed back to V1 from the complex of interest (at the temporal resolution of fMRI). These mappings were then used to generate activity in each functional complex, which was then tested for visual category selectivity (see “Category selectivity” and Formula 2). Note that at this point in the analysis, category selectivity (actual and activity-flow-mapped) was estimated for each vertex in each functional complex, then averaged (i.e., average selectivity of vertices in each complex). Thus, actual category selectivity in corresponding Results slightly varied from the region-level estimates based on fully distributed mappings (Fig. 1A) (which was estimated for each region, then averaged). Additionally, we removed outliers (using a conservative threshold of 5 median deviations) as before (see “Category selectivity”). The degrees of freedom are indicated in corresponding Results that report hypothesis testing statistics.

We also performed a control analysis testing whether each functional complex’s unique connectivity fingerprint with V1 determined its visual category selectivity better than a null rsFC network model. Here, we randomly permuted the connectivity architectures (per participant) 100 times but maintained edge strength and degree [134,150,151]. Note that this was used instead of rsFC substitution (see “Control analyses” above) because there was a variable number of vertices in each functional complex, and thus they could not be used to substitute each other’s connectivity fingerprints. Nonparametric estimates of statistical significance given by Max-T (see “Experimental design and statistical analyses”) were used to compare real mapped category selectivity to the null network mapped category selectivity.

Then, in steps 2+ (extended VIS) we used the mapped activation patterns from step 1 as the source activation patterns in activity flow mapping (Formula 1) and extended this to include all other VIS sub-system source vertices (Fig 1C). We repeated this until a “settling threshold” was reached. This was to simulate potential feedback and/or recurrent processes (initialized with stimulus-driven processes) that may occur over VIS network interactions. We defined a settling threshold as the step when the mapped activation patterns no longer changed within four decimal point precision, which can be thought of as the beginning of a horizontal asymptote in mapped activation values. We then performed all analyses as we did in step 1, to assess whether stimulus-driven activity flow processes further shaped by VIS network interactions improve the generation of visual category selectivity.

### Category selectivity generated by stimulus-driven network interactions further shaped by fully distributed network interactions

Expanding upon the analyses in the previous section (“Stimulus driven category selectivity”), we sought to test whether V1-initialized activity flow mappings (step 1) that are then processed over all cortical network interactions (step 2) (Fig 1D) – as opposed to a step 2 that incorporates only visual subsystems – would accurately generate visual category selectivity. We henceforth refer to this as the “stimulus-driven + fully distributed” mapping. Here, modulations conferred by all other functional networks (e.g., possible top-down effects from source regions in the dorsal attention network; DAN) were allowed to contribute to activity flow mapping in a second step. As before, we firstly initialized activity flow mapping with stimulus-driven interactions with V1 (as in “Stimulus-driven visual category selectivity” above). This first step of activity flow mapping was initialized with vertex-level data as in “Stimulus-driven visual category selectivity” above. Then we conducted an additional step that included all cortical source regions. Given the computational cost of vertex-level regularized regression (with cross-validation) across the whole cortex, this second step was performed at the region level. Following prior analyses, to control for multiple comparisons we used max-T nonparametric permutation testing (see “Experimental design and statistical analysis”). We used 100,000 permutations (i.e., p of 0.00001) and report statistical significance in terms of *p* < 0.00001.

### Experimental design and statistical analysis

Each functional complex that was proposed to be highly responsive to the visual categories of bodies, faces, places, and tools (see above: “Identification of functional complexes”) had estimates for actual activity and activity-flow-mapped activity. For each of these estimates and each visual category, we contrasted category responses to non-category responses (e.g., body vs non-body), hypothesizing that a functional complex would exhibit higher activity (in actual and mapped data) to the category vs the non-category. To test these hypotheses, we used one-tailed, paired samples t-tests on cross-participant data. For each category of interest, non-category activity was the average of the three remaining categories. For example, if body was the category of interest, responses to faces, places, and tools were averaged for non-body activations. This provided a consistent contrast across all four sets of analyses (i.e., always category vs non-category).

In every set of analyses involving a comparison, we used the Max-T nonparametric permutation approach (shuffling conditions over 100,000 permutations – i.e., p-values of 0.00001 – unless otherwise stated) to correct for multiple comparisons [152]. For example, the Max-T approach was used when testing whether category selectivity scores (Formula 2) revealed for each functional complex were greater than 1.0 (the null mean), as well as testing whether estimates of distributed processing contributions (Formula 4) were greater than 50% (see “Category selectivity” for more on these measures). We likewise used Max-T (here with 10,000 permutations for computational tractability) to address multiple comparisons in the network-based analyses (i.e., comparing each network-level result to every other network-level result). When comparing estimates based on each network to every other network, the max-T threshold statistic that was surpassed for significance is reported, instead of a lengthy list of observed t-statistics. The max-T procedure was used in any analysis where we could not assume a normal distribution a priori. Note that for category selectivity and percent distributed processing analyses, degrees of freedom were based on sample size with outlier participants removed (see “Category selectivity”). To assess accuracy of the activity-flow-mapped (Formula 1) results, we compared the actual and activity-flow-mapped activations in each of the four visual category analyses with Pearson’s r, MAE, and R^2^ (Fig 2). All findings are reported for cortical regions in the right hemisphere for the discovery dataset. Left hemisphere and replication dataset results are reported in Supporting information.

### Code, software, and data accessibility

All MATLAB, Python, demonstration code, and software are publicly available on GitHub. The main repository for the present study can be found here: https://github.com/ColeLab/ActFlowCategories. The Activity Flow Toolbox and related software [27,28] can be found here: https://github.com/ColeLab/ActflowToolbox. Further, the HCP maintains open access imaging data at various levels of processing here: https://www.humanconnectome.org/. Data at other levels of processing or analysis pertinent to the present study are available upon request.

## Results

### Localized response profiles are captured by activity flow processes

Our preliminary hypothesis was that activity flow processes could accurately map responses in extensively investigated category-selective visual cortex regions. Firstly, we extended prior work [20,27,28] by implementing a resting-state functional connectivity (rsFC) method [125] that estimates a more causally plausible network (Fig 3A and 3B), and by analyzing a large cohort (N=352) split into discovery and replication datasets. These updates serve to improve the inferential validity of activity flow mapping results [18]. Secondly, we quantified response profiles (also termed population receptive fields) of functional complexes identified as integral to four visual categories (Tables 1 and 2). A response profile (rows of Fig 3D) was defined as the task-evoked activity levels across all 24 HCP conditions. Establishing response profiles allowed us to verify that each complex was responsive to their respective visual categories (outlined in black: Fig 3D, right) and verify that activity flow mapping (Fig 3D, left) was reliable across a variety of cognitive domains.

Activity-flow-mapped response profiles maintained high accuracy in both hemispheres (right: r = 0.92, MAE = 3.93, R^2^ = 0.80; left: r = 0.92, MAE = 3.96, R^2^ = 0.79; replication dataset: S1 Table), which corroborates prior findings that activity flow processes (specified by connectivity patterns) can accurately map task-evoked activity in held out brain regions across diverse cognitive domains [27]. Moreover, this suggests that intrinsic (i.e., resting-state) functional network organization (Fig 3C) plays a prominent role in generating localized response profiles. Subsequent analyses examined the processes involved in category-selective responses (i.e., focusing on black-outlined conditions in Fig 3D), as generated by activity flow mapping.

### Localized visual category selective responses are generated via empirically-derived models of fully distributed network interactions

We next tested the hypothesis that the classically observed selectivity for visual categories in select visual cortex regions is shaped by their unique connectivity fingerprints distributed across cortex (Fig 1A). An emerging literature suggests that network interactions and/or distributed activation patterns are integral to visual category processing [11,14,91,127]. However, the nature of the network interactions underlying the generation of visual category selectivity remains unclear, as well as the extent to which those distributed interactions account for localized category selectivity (relative to local, within-region processes). Here, we sought to test the generative capacity of distributed network interactions by modeling activity flow processes over intrinsic functional connections [27]. The four visual category models are reported in separate sections below with similar formatting to aid cross-comparison. The functional complexes identified based on prior literature (Fig 4, Tables 1 and 2) were anatomically positioned along occipital (anterior and lateral to V1) and temporal (tending posterior and ventral to A1) cortices, which reflects the associative processing demands of visual category computations [47,48]. Functional complex regions belonged primarily to the secondary visual (VIS2) network, with additional locations in dorsal attention (DAN), default mode (DMN), and language networks (LAN) (Fig 4) [25].

**Fig 4.**
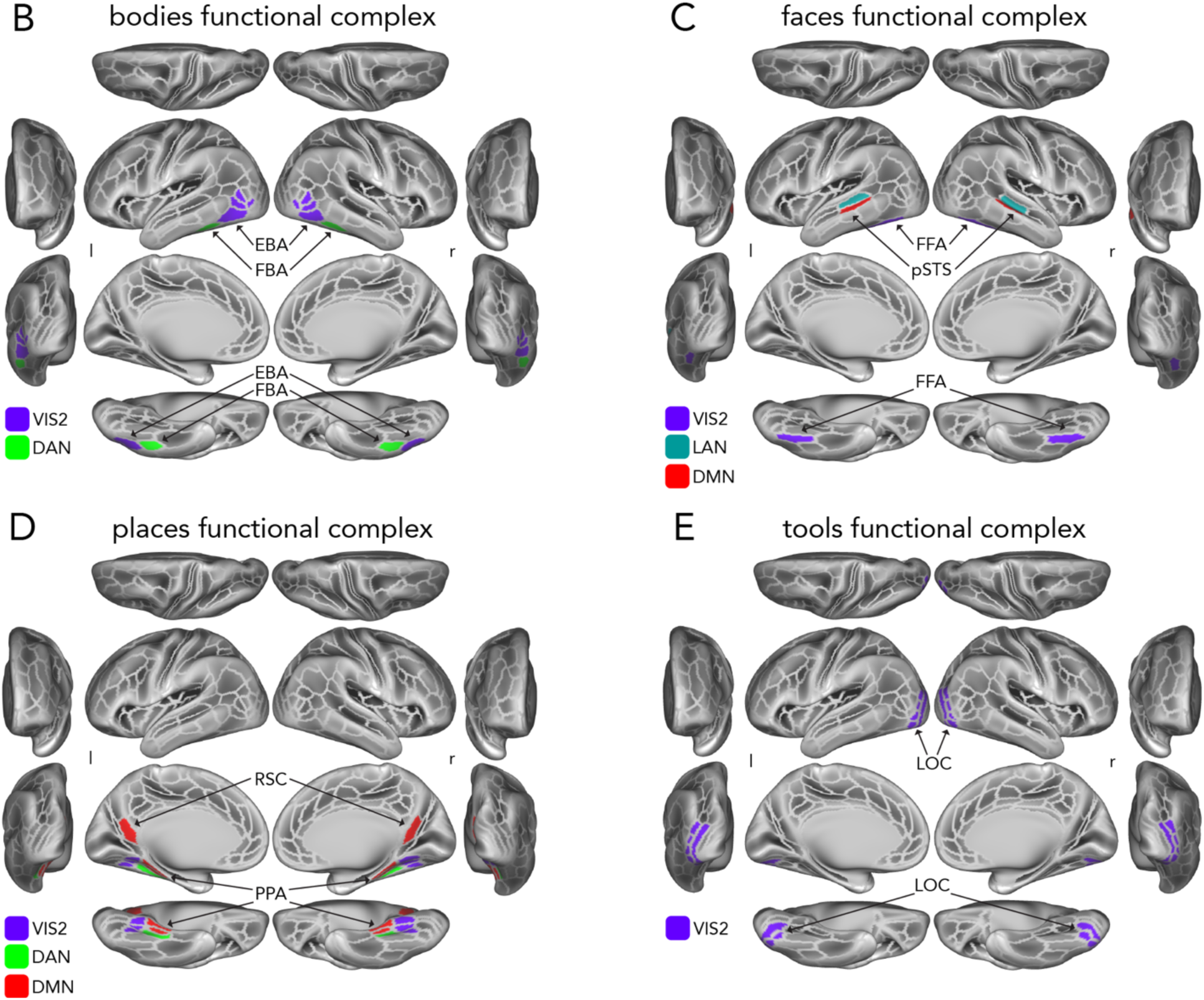
Visual categories and identified functional complexes. (**A**) [Please note, panel A has been excluded from the preprint due to privacy concerns.] Exemplar stimuli from each of the four visual semantic categories shown to participants across the n-back task (bodies, faces, places, and tools). Please note that the face example is a photograph of the author of this manuscript, meant to be representative (but not an exact match) of the original face stimuli while protecting privacy concerns of the original models. In each category, stimuli varied to sample a wide array of representations. Body images included whole bodies (no faces) and isolated body parts (excluding nudity); face images included diverse ages and facial expressions; place images included indoor and outdoor scenes (and combinations, e.g., a patio); and tool images included distinct items (e.g., plier versus drill) as well as similar variants (e.g., two drills: one white, one blue). See Barch et al. [45] and www.humanconnectome.org for further details. (**B**) A schematic of the cortical surface in three gross orientations, with MMP regions [54]. outlined in silver and functional complexes relevant to processing body features (see Tables 1 and 2, Methods) colored in based on CAB-NP network affiliations ([25], Fig 1C) and identified with text labels and arrows. EBA = extrastriate body area; FBA = fusiform body area. VIS2 = secondary visual network; DAN = dorsal attention network. “r” denotes right hemisphere; “l” denotes left hemisphere. (**C**) Functional complexes related to face processing. FFA = fusiform body area; pSTS = posterior superior temporal sulcus; LAN = language network; DMN = default mode network. (**D**) Functional complexes related to places and scenes. RSC = retrosplenial complex; PPA = parahippocampal place area. (**E**) Functional complexes related to processing images of tools and objects. LOC = lateral occipital complex.

We tested the hypothesis that fully distributed activity flow processes (Fig 1A) – shaped by the intrinsic functional connectivity fingerprint of each functional complex (Fig 4) – are sufficient to generate category-selective responses without the need for additional within-region processing. Before this, we began with two confirmations essential for subsequent tests of this hypothesis. We first constrained prior activity flow mappings [27,28] to body, face, place, and tool conditions and compared actual versus activity-flow-mapped responses across the whole cortex (S1 Fig). We then focused on activations in each functional complex (Fig 4) and in each hemisphere and quantified “benchmark” contrasts to corroborate the literature by comparing the category of interest versus non-category responses (e.g., body versus non-body) in the actual and activity-flow-mapped data (note that category selectivity is reported subsequently). Whole-cortex, activity-flow-mapped responses to visual categories were highly accurate (bodies: r = 0.89, MAE = 5.27, R^2^ = 0.78; faces: r = 0.86, MAE = 5.83, R^2^ = 0.72; places: r = 0.88, MAE = 5.85, R^2^= 0.77; tools: r = 0.89, MAE = 5.62, R^2^ = 0.78; S1 Fig). Significant benchmark contrasts in each functional complex were also observed in both the actual (bodies: t(175) = 26.65, Cohen’s d = 2.01, faces: t(175) = 20.37, Cohen’s d = 1.54; places: t(175) = 39.53, Cohen’s d = 2.89; tools: t(175) = 8.49, Cohen’s d = 0.64; all p < 0.0001) and activity-flow-mapped data (bodies: t(175) = 17.91, Cohen’s d = 1.35; faces: t(175) = 10.06, Cohen’s d = 0.76; places: t(175) = 26.06, Cohen’s d = 1.88; tools: t(175) = 9.94, Cohen’s d = 0.75; all p < 0.0001) (S2 Fig) (left hemisphere and replication: S1 and S2 Tables). In each set of benchmark contrast analyses, multiple comparisons correction was performed with nonparametric permutation testing (i.e., the “max-T” procedure described in the Methods; here, with 10,000 permutations for each set of analyses).

#### Distributed activity flowing over intrinsic connectivity generates localized body category selectivity

Processing of human body images (Fig 4A) is associated with the fusiform body area (FBA) and the extrastriate body area (EBA) [2,3,71], which are located in the lateral occipito-temporal cortex (Tables 1 and 2, Fig 4B). EBA computations are thought to discriminate between bodily attributes and parts [153] and provide postural information to frontoparietal regions [72], indicating a pivotal role in action planning. The FBA is thought to process images of whole bodies [3] and pairs of body parts [154]. Extending the fully distributed hypothesis to the processing of body images, we hypothesized that activity flowing via the intrinsic connectivity fingerprint of the EBA and FBA (henceforth: EBA/FBA) (Figs 5A and 5B) — its unique pattern of distributed cortical connections — is likely the primary determinant of its body-selective responses.

**Fig 5.**
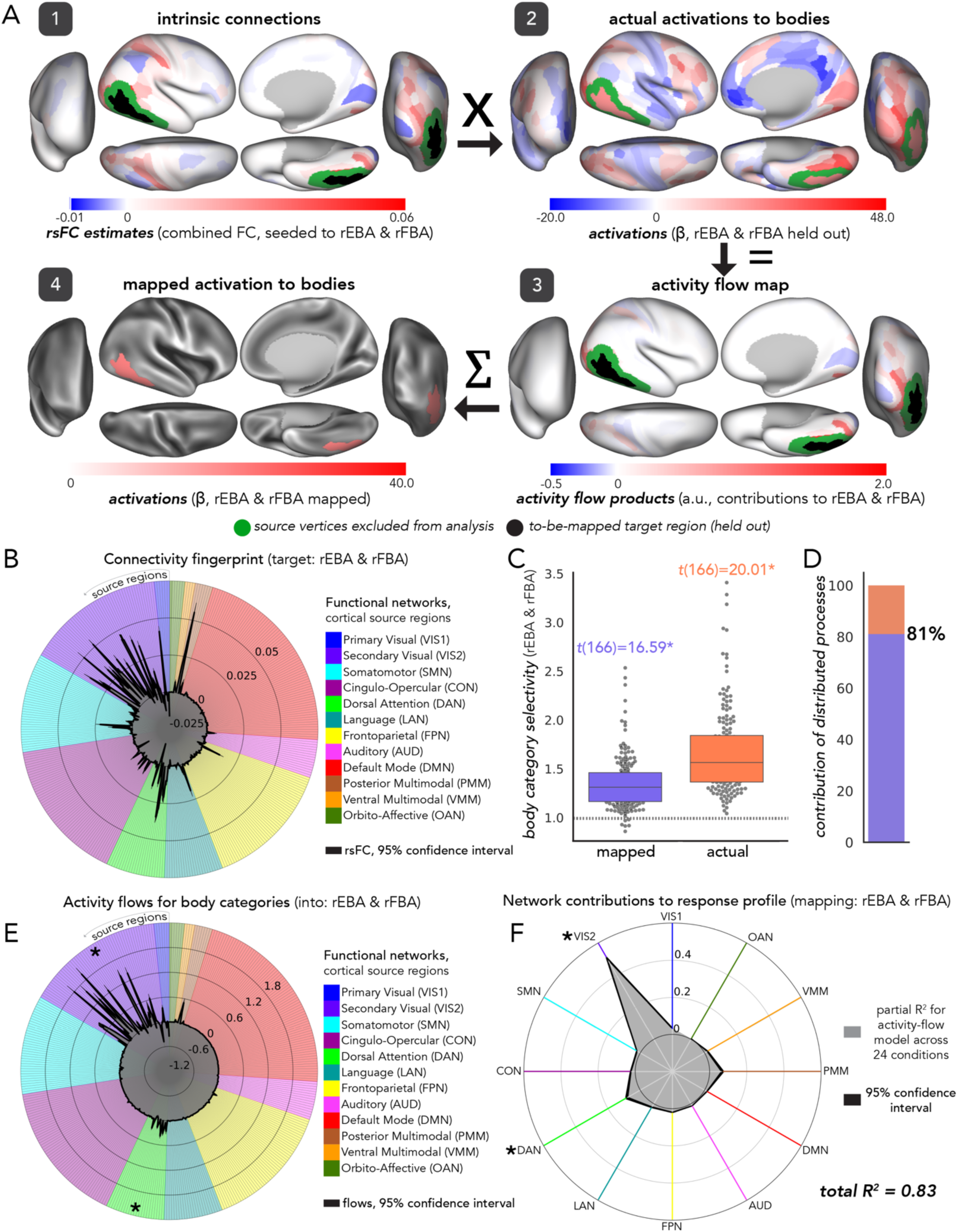
Distributed activity flows account for the majority of localized selectivity to bodies. (**A**) The activity flow mapping procedure (steps match Fig 2C) generating body responses in EBA/FBA (black), projected onto cortical schematics (right hemisphere). Green: source vertices excluded from analyses. Step 2 was not blacked out for visual comparison with step 4, however it was held out in-analysis. Step 4 color scale shows maximum of all regions’ mapped activations to body images for visual comparison with step 2. (**B**) The connectivity fingerprint [13,15] of the right EBA/FBA via rsFC (black lines). Radial lines: source regions connected to EBA/FBA, clustered by functional network assignments [25] (colored per legend; Fig 3C). 95% confidence interval: across participants. (**C**) Right EBA/FBA body selectivity: activity-flow-mapped (purple) and actual (coral). Gray dots: individual participants’ scores. Dashed line: no selectivity (1.0); used for comparison in statistical tests. Significant t-statistics are indicated with an asterisk (p<0.00001; see Methods). (**D**) Estimated contribution of distributed network interactions to body selective responses in EBA/FBA. (**E**) The activity flows (as in A3) of each source region contributing to mapped EBA/FBA responses to body images (statistical significance asterisks at network-mean level). (**F**) Variance explained by each network-restricted activity flow model (unmixed partial R^2^ in gray) of EBA/FBA’s response profile. Black lines: 95% confidence interval across participants. Asterisks: statistical significance versus each other network. E-F suggests that activity flowing over interactions with VIS2 and DAN represents a general network coding mechanism for EBA/FBA. See Methods for full details of each analysis.

We assessed *body selectivity* in the EBA/FBA — quantified as the ratio of body to non-body category activations — in both activity-flow-mapped (Fig 5C, purple) and actual (Fig 5C, coral) data. As expected, mapped and actual body selectivity were statistically significant relative to a selectivity ratio of 1.0 (i.e., greater than the null hypothesis of equivalent responses to bodies and non-bodies; *p* < 0.00001; see Methods): activity-flow-mapped mean body selectivity = 1.35, *t*(166) =16.59, Cohen’s *d* = 1.29; actual mean body selectivity = 1.67, *t*(166) = 20.01,Cohen’s *d* = 1.55; Fig 5C). We next estimated the contribution of distributed activity flow processes to body selectivity via the percentage of actual body selectivity captured by mapped body selectivity (Fig 5D). Given that the fully distributed activity flow approach maps task-evoked activity based on distributed features ([27]; Fig 2B), this ratio approximates the extent that distributed network interactions contribute to the generation of body selectivity in the EBA/FBA (relative to local, within-complex processes and/or error). As expected, the estimated contribution of distributed activity flow processes was 81%, which was statistically greater than 50% (*p* < 0.00001; see Methods) (Cohen’s *d* = 2.52, *t*(166) = 32.49, Fig 5D) (left hemisphere and replication: S3 Table).

Next, we examined how each large-scale functional network (Fig 3C [25]) contributed to the observation that activity flowing over a fully distributed intrinsic network architecture (connectivity fingerprint: Fig 5B) shaped body selectivity in the EBA/FBA. The activity flows map (Fig 5A, step 3) — where source activity was weighted by rsFC with the (held out) target region — was an entry point to assess such network contributions. Firstly, we averaged the estimated activity flows based on network assignment (Fig 5E). Secondly, we used dominance analysis to identify each functional network’s unmixed contribution (partial R^2^, see Methods) to the activity-flow-mapped response profile (i.e., activations across all 24 conditions, Fig 3D) of the EBA/FBA (Fig 5F). Secondary visual network (VIS2) activity flows exhibited the largest contribution (versus each other network) to EBA/FBA’s body-evoked activations (Fig 5E, all *t*-statistics above the max-T threshold (175) = 3.39, *p* < 0.0001; left hemisphere and replication: S4 Table). DAN activity flows were also significantly higher (same max-T thresholds, see Methods) than the other networks, except VIS2 and posterior multimodal network (PMM). VIS2 also accounted for most of the explained variance in EBA/FBA’s response profile (partial R^2^ = 0.51, 61% of the full model R^2^, Fig 5F). DAN accounted for the next highest amount of explained variance (partial R^2^ = 0.09, 10% of the full model R^2^, Fig 5F; left hemisphere and replication: S5-S7 Tables). Thus, for both body-specific conditions and across all conditions (i.e., response profile), activity flowing over VIS2 and DAN exhibited substantial contributions to EBA/FBA activation patterns.

As hypothesized by the fully distributed network interaction model (Fig 1A), we found evidence that activity flowing over resting-state functional connections distributed throughout cortex shaped body selectivity in the EBA/FBA. Results also suggested that distributed processes chiefly specified localized body-selective responses in the EBA/FBA. The total explained variance in the EBA/FBA’s 24-condition response profile was significantly greater than 50% (total R^2^ = 0.83, versus 0.5: *t*(175) = 66.07, *p* = 1.14 x 10^-125^; left hemisphere and replication: S5-S7 Tables), suggesting that distributed activity flow processes predominantly influence EBA/FBA responses across a variety of cognitive domains.

#### Distributed activity flowing over resting-state connectivity generates localized face category selectivity

The fusiform face area (FFA) is a hallmark region exhibiting localized category selectivity given its well-replicated relationship with face processing [4, 49]. The posterior superior temporal sulcus (pSTS) is thought to be a multisensory processing region responsive to the changeable aspects of faces [21,155]. The FFA and pSTS are loci of specialization in a two-pathway model [87,91], representing facial identity (relatively invariant) and facial expression (dynamic), respectively [11,92]. While both the FFA and pSTS are part of the ventral temporal cortex, the pSTS is more dorsal and lies adjacent to the temporoparietal junction; the FFA is more posterior and typically identified on the ventral aspect of the cortex (i.e., the fusiform gyrus) (Tables 1 and 2, Fig 4C). Extending the fully distributed hypothesis (Fig 1A) to the processing of face images, we hypothesized that activity flow processes via the whole-cortex, resting-state connectivity fingerprint of the FFA and pSTS (Figs 6A and 6B) (henceforth: FFA/pSTS) is sufficient to determine its face selective responses.

**Fig 6.**
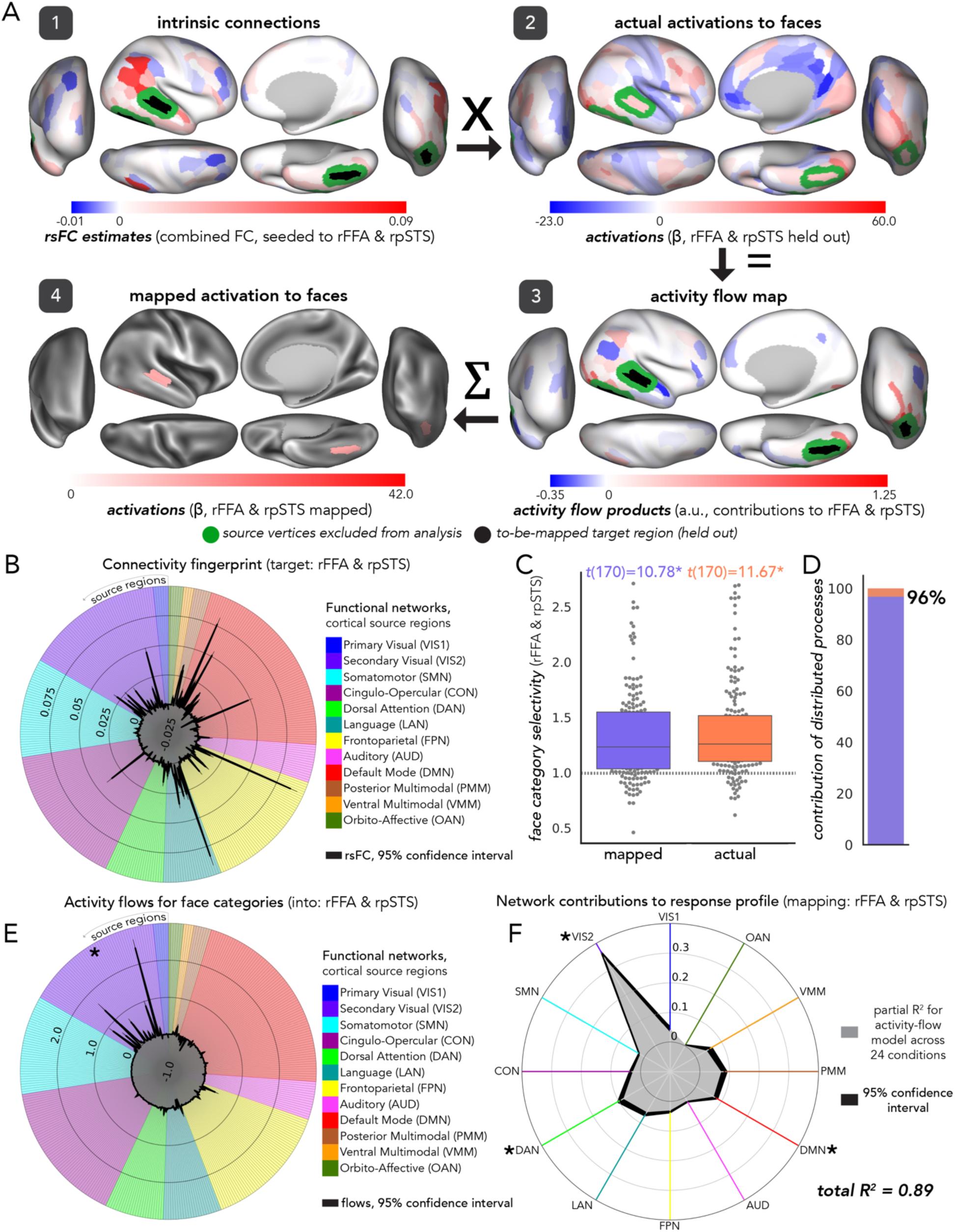
Distributed activity flows account for the majority of localized selectivity to faces. All figure specifications follow Fig 5. (**A**) The activity flow procedure mapping activations to face categories, projected onto cortical schematics (right hemisphere). Right FFA/pSTS was the held-out target complex. (**B**) The connectivity fingerprint [13,15] of the right FFA/pSTS via whole-cortex rsFC (black lines). Radial lines: source regions connected to the FFA/pSTS, clustered by functional network assignments [25] (colored per legend and Fig 3C). (**C**) Face category selectivity exhibited by the right FFA/pSTS. Significant t-statistics are indicated with an asterisk (p<0.00001; see Methods). (**D**) Estimated contribution of distributed activity flow processes to face selectivity exhibited by the right FFA/pSTS. (**E**) Activity flows (as in A step 3) of each source region contributing to the mapping of FFA/pSTS responses to face images. VIS2 regions contributed most to the FFA/pSTS mapped activation magnitude to faces. (**F**) Variance explained by each network-restricted activity flow model (unmixed partial R^2^ via dominance analysis; Methods) of the right FFA/pSTS’ response profile. VIS2 accounted for the most variance, altogether suggesting that activity flowing over VIS2 regions represents a general network coding mechanism for FFA/pSTS processing. DAN and DMN regions also accounted for a nontrivial amount of variance at the response-profile level suggesting that, across diverse cognitive domains, FFA/pSTS processing is impacted by activity flowing over DAN and DMN regions, in addition to VIS2 (in the face-specific case).

We followed the foregoing pipeline for body selectivity exactly, except analyzing face images and the FFA/pSTS (Fig 6A, black regions). Overall, as expected, the pattern of results observed for body processing (EBA/FBA) extended to face processing (FFA/pSTS). Significant face selectivity was observed in both the actual and activity-flow-mapped data (mapped mean face selectivity = 1.33, *t*(170) = 10.78, Cohen’s *d* = 0.83; actual mean face selectivity = 1.37, *t*(170) = 11.67, Cohen’s *d* = 0.89; *p* < 0.00001; Fig 6C), and the estimated contribution of distributed activity flow processes was 96% (*t*(170) = 29.61, Cohen’s *d* = 2.27, *p* < 0.00001; Fig 6D) (left hemisphere and replication: S3 Table). We found that face activations in the FFA/pSTS were most influenced by activity flows over VIS2 connections (max-T threshold (175) = 3.39, *p* < 0.0001; Fig 6E; left hemisphere and replication: S4 Table). VIS2 also accounted for the most variance in FFA/pSTS processing at the response profile level (40%; left hemisphere and replication: S5-S7 Tables).

These results suggest that face selectivity in the FFA/pSTS was significantly shaped by activity flow processes over a fully distributed intrinsic network architecture. These processes were most prominently influenced by activity flowing over VIS2, similar to the EBA/FBA. Across all conditions, activity flowing over dorsal attention (DAN) and default mode (DMN) network connections were predictive as well (DAN: 11%, DMN: 9%). The total explained variance in this response profile model (Fig 6F) was R^2^ = 0.89 (versus 0.5: *t*(175) = 92.61, *p* = 1.32 x 10^-150^; left hemisphere and replication: S5-S7 Tables), suggesting that distributed processes chiefly influenced FFA/pSTS responses across many cognitive domains.

#### Distributed activity flowing over intrinsic connectivity generates localized place category selectivity

We next focused on place images (Fig 4A), sometimes termed scenes, environment, or topography. Place-specific regions include the parahippocampal place area (PPA) and the retrosplenial cortex (RSC) (Tables 1 and 2, Fig 4D), which are thought to act cooperatively toward a cohesive percept [5,97,101]. The PPA is thought to compute viewpoint-specific discrimination [156], as well as mediating (or binding) contextual associations pertinent for episodic memory [157]. The RSC is thought to provide the medial temporal lobe with visuospatial information [158], and to integrate viewpoint-invariant information for navigation and learning [5]. The PPA consisted of previously identified regions [51], and the dorsal RSC corresponded to Brodmann areas 29 and 30 [159] (Fig 4D). Extending the fully distributed hypothesis (Fig 1A) to the processing of place images, we hypothesized that activity flow via the whole-cortex, resting-state connectivity fingerprint of the PPA and RSC (Figs 7A and 7B) (henceforth: PPA/RSC) significantly determines its place-selective responses.

**Fig 7.**
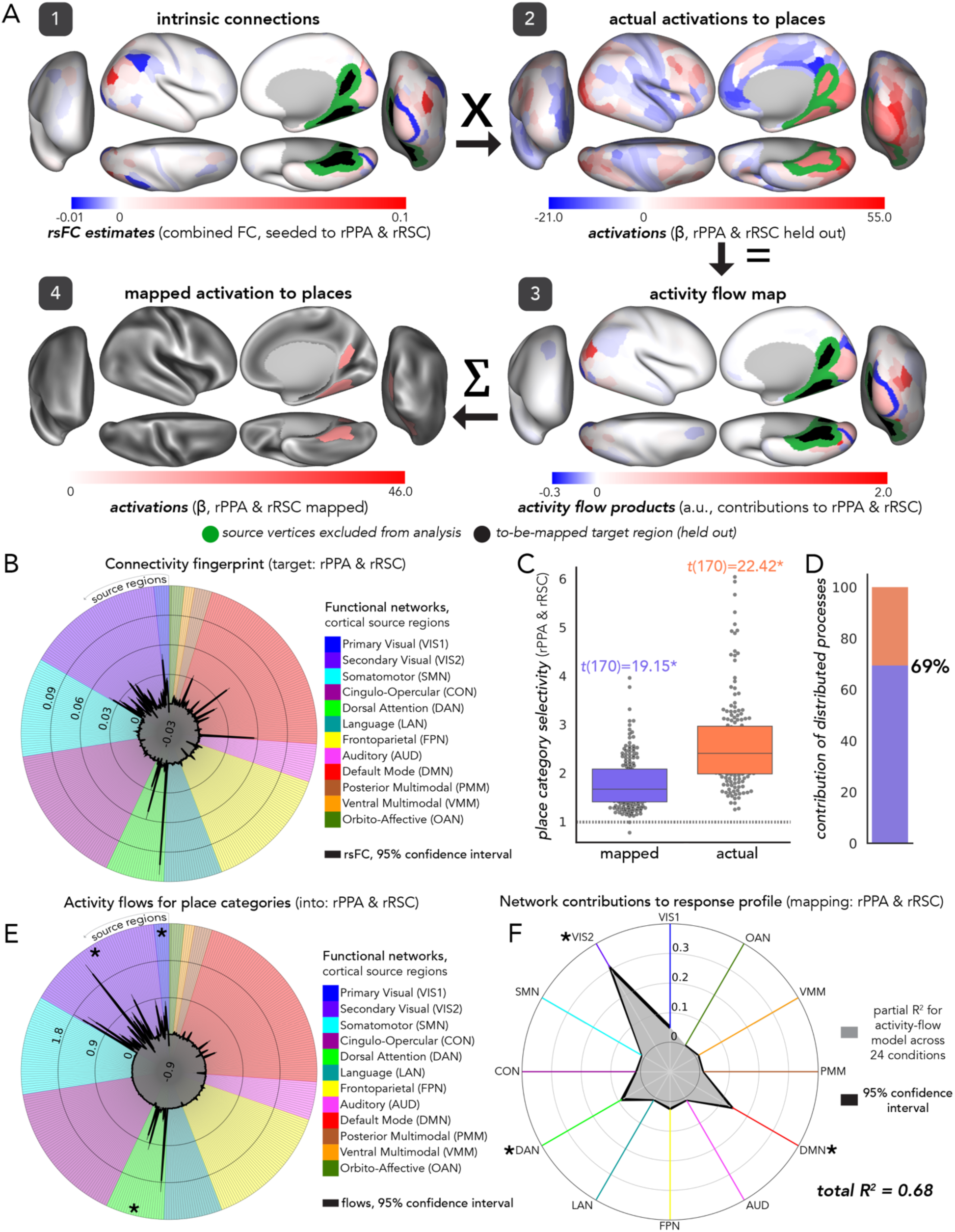
Distributed activity flows account for the majority of localized selectivity to places. All formatting and figure specifications follow Fig 5. (**A**) Activity flow mapping of activations to place categories, projected onto cortical schematics (right hemisphere only). Right PPA/RSC was the held-out target complex. (**B**) The connectivity fingerprint of the right PPA/RSC, as in Passingham et al. [13] except with whole-cortex rsFC (black lines). Radial lines: source regions connected to the PPA/RSC, clustered by CAB-NP functional network assignments [25] (colored per legend and Fig 3C). (**C**) Place category selectivity exhibited by the PPA and RSC in the right hemisphere. Significant t-statistics are indicated with an asterisk (p<0.00001; see Methods). (**D**) Estimated contribution of distributed activity flow processes to the emergence of place selective responses in the right PPA/RSC. (**E**) Activity flows (as in A step 3) of each source region contributing to the mapping of PPA/RSC responses to place images. VIS1, VIS2 and DAN contributed most to the right PPA/RSC mapped activation magnitude to place categories. Note that VIS1, VIS2, and DAN were all statistically greater than each other network, except for each other. (**F**) Variance explained by each network-restricted activity flow model (partial R^2^ via dominance analysis; Methods) of the right PPA/RSC’s response profile. VIS2, DAN, and DMN accounted for the most variance, suggesting that activity flowing over regions in these networks represents a general network coding mechanism for PPA and RSC processing, while VIS1 contributes to place-specific responses.

Significant place selectivity was observed in the PPA/RSC (mapped mean place selectivity = 1.8, *t*(170) = 19.15, Cohen’s *d* = 1.49; actual mean place selectivity = 2.59, *t*(170) = 22.42, Cohen’s *d* = 1.74; *p* < 0.00001; Fig 7C), and the estimated contribution of distributed processes was 69% (versus 50%: *t*(170) = 14.87, Cohen’s *d* = 1.15, *p* < 0.00001; Fig 7D) (left hemisphere and replication: S3 Table). Activity flowing over VIS1, VIS2, and DAN provided the largest contributions to PPA/RSC’s place activations (all significant except when compared to each other: max-T threshold (175) = 3.4, *p* < 0.0001; Fig 7E) (left hemisphere and replication: S4 Table). A similar set of networks accounted for the most variance at the response profile level (Fig 7F), including VIS2 (44%), DAN (13%) and DMN contributions (21%) (left hemisphere and replication: S5-S7 Tables).

As hypothesized by the fully distributed model of Fig 1A, the whole-cortex connectivity fingerprint of the PPA/RSC significantly shaped its place selectivity. PPA/RSC responses were influenced by activity flowing over VIS2 and DAN. Additionally, VIS1 was particularly important for place-specific activity; and DMN for cross-domain activity (Figs 7E and 7F), suggesting that PPA/RSC’s activity flow processes were more heterogeneous than prior models. The total variance in the PPA/RSC response profile explained by activity flow processes (Fig 7F) was greater than 50% (total R^2^ = 0.68; *t*(175) = 23.93, *p* = 4.51 x 10^-57^; left hemisphere and replication: S5-S7 Tables), which provides evidence that distributed processes predominantly influenced PPA/RSC activations to a variety of cognitive domains.

#### Distributed activity flowing over intrinsic connectivity generates localized tool category selectivity

The final visual category included tool images (Fig 4A), sometimes termed inanimate objects. Following an extensive literature on object recognition [52,160], we hypothesized that tool selectivity is exhibited by the lateral occipital complex (LOC) [21,161], which is posteriorly located and wraps around the cortex in the ventromedial direction (Fig 4E). The LOC is thought to represent higher-level information of objects, as opposed to low level visual features [162], suggesting a role at the category level of visual processing. However, reports vary on the degree of semantic content processed by the LOC [163]. Thus, the link between the LOC and tool selectivity was the least clear (of all four models) *a priori*. Extending the fully distributed hypothesis (Fig 1A) to the processing of tool images, we hypothesized that activity flow via the whole-cortex, resting-state connectivity fingerprint of the LOC (Figs 8A and 8B) significantly determines its tool selective responses.

**Fig 8.**
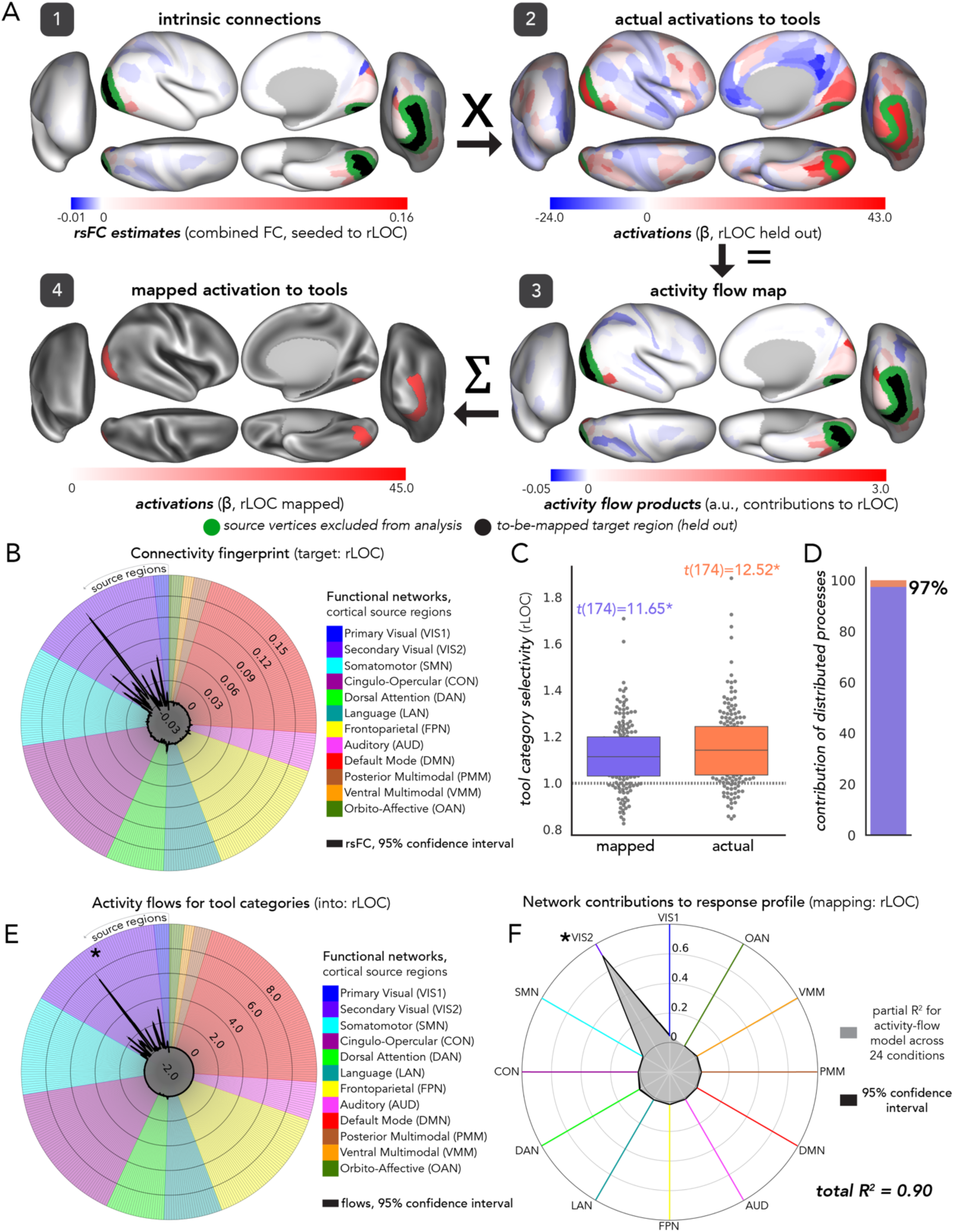
Distributed activity flows account for the majority of localized selectivity to tools. All formatting and figure specifications as in Fig 5. (**A**) Activity flow mapping of activations to tool categories in the held-out target - the right LOC - projected onto cortical schematics (right hemisphere). (**B**) The connectivity fingerprint of the right LOC, as in Passingham et al. [13] except with whole-cortex rsFC (black lines). Radial lines: source regions connected to the LOC, clustered by CAB-NP functional network assignments [25] (colored per legend and Fig 3C). (**C**) Tool category selectivity exhibited by the LOC in the right hemisphere. Significant t-statistics are indicated with an asterisk (p<0.00001; see Methods). (**D**) Estimated contribution of distributed activity flow processes to the emergence of tool selective responses in the right LOC. (**E**) Activity flows (as in A step 3) of each source region contributing to the mapping of LOC responses to tool images. VIS2 contributed most to the right LOC mapped activation magnitude to tool categories (**F**) Variance explained by each network-restricted activity flow model (partial R^2^ via dominance analysis; Methods) of the right LOC’s response profile. VIS2 accounted for the most variance, suggesting that activity flowing over regions in these networks represents a network coding mechanism for LOC processing.

Tool selectivity in the LOC was statistically significant (mapped tool selectivity = 1.12, *t*(174) = 11.65, Cohen’s *d* = 0.88; actual tool selectivity = 1.15, *t*(174) = 12.52, Cohen’s *d* = 0.95; *p* < 0.00001; Fig 8C), and the estimated contribution of distributed activity flow processes to tool selectivity in the LOC was particularly high at 97% (*t*(174) =111.44 ,Cohen’s *d* = 8.45; Fig 8D) (left hemisphere and replication: S3 Table; given evidence in the literature that tool selectivity is left lateralized, future work building on activity-flow-mapped tool selectivity should pay special attention to the tool processing results reported in the Supplement). VIS2 activity flows demonstrated a strikingly high contribution to LOC’s tool-specific responses (max-T threshold (175) = 3.4, *p* < 0.0001; Fig 8E) (left hemisphere and replication: S4 Table) and accounted for the majority of the variance in LOC’s response profile (79%; Fig 8F) (left hemisphere and replication: S5-S7 Tables).

As hypothesized, tool selectivity was strongly influenced by distributed activity flow processes (97-98%). As in all other models (bodies, faces, and places) the total explained variance that activity flow mapping captured for the LOC response profile was greater than 50% (total R^2^ = 0.90 vs 0.5: *t*(175) = 115.55, *p* = 3.71 x 10^-167^). Given that the standard application of activity flow mapping [27,28] (i.e., following the fully distributed network interaction scheme of Fig 1A) is a distributed processing model (Fig 2B), this suggests that distributed processes may be the primary mechanism generating category-selective LOC activations.

### Control analyses

#### Null connectivity fingerprints reduce visual category selectivity

To corroborate that each functional complex’s unique whole-cortex resting-state connectivity fingerprint (Figs 5B-8B) — its placement in the brain’s intrinsic network architecture — determined its visual category selectivity (Figs 5C-8C), we built null models based on FC substitution. We hypothesized that a null connectivity fingerprint – here defined as a fingerprint based on the wrong functional complex – would confer significantly lower activity-flow-mapped category selectivity.

As hypothesized, visual category selectivities were significantly greater when based upon the true (i.e., empirically-based) whole-cortex connectivity fingerprints (Table 3, reporting right hemisphere functional complexes and the discovery dataset). This assessment is more stringent than a null model that randomly scrambles FC because there is likely some similarity in visual processing (and therefore activity flow patterns) across the four functional complex’s fully distributed network interactions. However, no substituted connectivity fingerprint was sufficiently similar to the true fingerprint to generate the original mapped category selectivity. Each of these results were corroborated in the left hemisphere and in the replication dataset (S8 and S9 Tables). These results corroborate that the fully distributed, resting-state connectivity fingerprint unique to each functional complex significantly shaped its visual category selective response. Given that connectivity fingerprints are the bases of activity flow processes (Fig 1A, Formula 1) that were used to generate mapped category selectivity, these results further support the proposition that fully distributed network interactions are sufficient to generate localized visual category selectivity in all four functional complexes tested.

**Table 3.** Empirical (true) resting-state connectivity fingerprints generate visual category selectivity significantly better than null model connectivity fingerprints.

| Comparison | *t*(175) | *p*-value | Cohen's *d* |
| --- | --- | --- | --- |
| True model body selectivity > null model body selectivity when rEBA/rFBA rsFC was substituted with: |  |  |  |
| face complex connectivity fingerprint: rFFA/rpSTS rsFC | 6.36 | $1.7 \times 10^{-9}$ | 0.61 |
| place complex connectivity fingerprint: rPPA/rRSC rsFC | 3.53 | 0.0005 | 0.37 |
| tool complex connectivity fingerprint: rLOC rsFC | 11.34 | $1.1 \times 10^{-22}$ | 1.16 |
| True model face selectivity > null model face selectivity when rFFA/rpSTS rsFC was substituted with: |  |  |  |
| body complex connectivity fingerprint: rEBA/rFBA rsFC | 8.21 | $4.7 \times 10^{-14}$ | 0.81 |
| place complex connectivity fingerprint: rPPA/rRSC rsFC | 21.45 | $7.1 \times 10^{-51}$ | 2.11 |
| tool complex connectivity fingerprint: rLOC rsFC | 14.95 | $4.4 \times 10^{-33}$ | 1.48 |
| True model place selectivity > null model place selectivity when rPPA/rRSC rsFC was substituted with: |  |  |  |
| body complex connectivity fingerprint: rEBA/rFBA rsFC | 14.02 | $2.1 \times 10^{-30}$ | 1.50 |
| face complex connectivity fingerprint: rFFA/rpSTS rsFC | 14.46 | $1.1 \times 10^{-31}$ | 1.44 |
| tool complex connectivity fingerprint: rLOC rsFC | 11.87 | $2.3 \times 10^{-24}$ | 1.20 |
| True model tool selectivity > null model tool selectivity when rLOC rsFC was substituted with: |  |  |  |
| body complex connectivity fingerprint: rEBA/rFBA rsFC | 2.74 | 0.007 | 0.23 |
| face complex connectivity fingerprint: rFFA/rpSTS rsFC | 9.97 | $7.8 \times 10^{-19}$ | 0.91 |
| place complex connectivity fingerprint: rPPA/RSC rsFC | 2.06 | 0.04 | 0.22 |

*The prefix “r” indicates the right hemisphere. rsFC = resting-state functional connectivity*

#### Activity-flow-mapped category selectivity was not driven by category responsiveness in other parts of the brain

A potential confound in our analysis pipeline was that the criteria for identifying functional complexes (see Methods) excluded some brain regions that also exhibit a degree of visual category selectivity and/or responsiveness (Fig. 9A). Therefore, it is possible that the findings in the preceding sections – that fully distributed activity flow processes are capable of generating localized visual category selectivity – were driven by these other, excluded regions (which remained as sources of task-evoked activity). To address this, we held out select regions (Fig. 9A) from the source sets of each visual category selectivity analysis and compared such source-set-controlled findings to the fully-distributed model findings (Fig. 1A; Figs. 5-8). Individual differences in selectivity were preserved across all visual categories (Fig. 9B), suggesting that activity flow processes were capable of generating visual category selectivity in each functional complex without the influence of other visual category responsive regions. Note that the following sections on stimulus-driven network interactions more fully address this circularity concern, while also testing refined versions of our core hypothesis.

**Fig 9.**
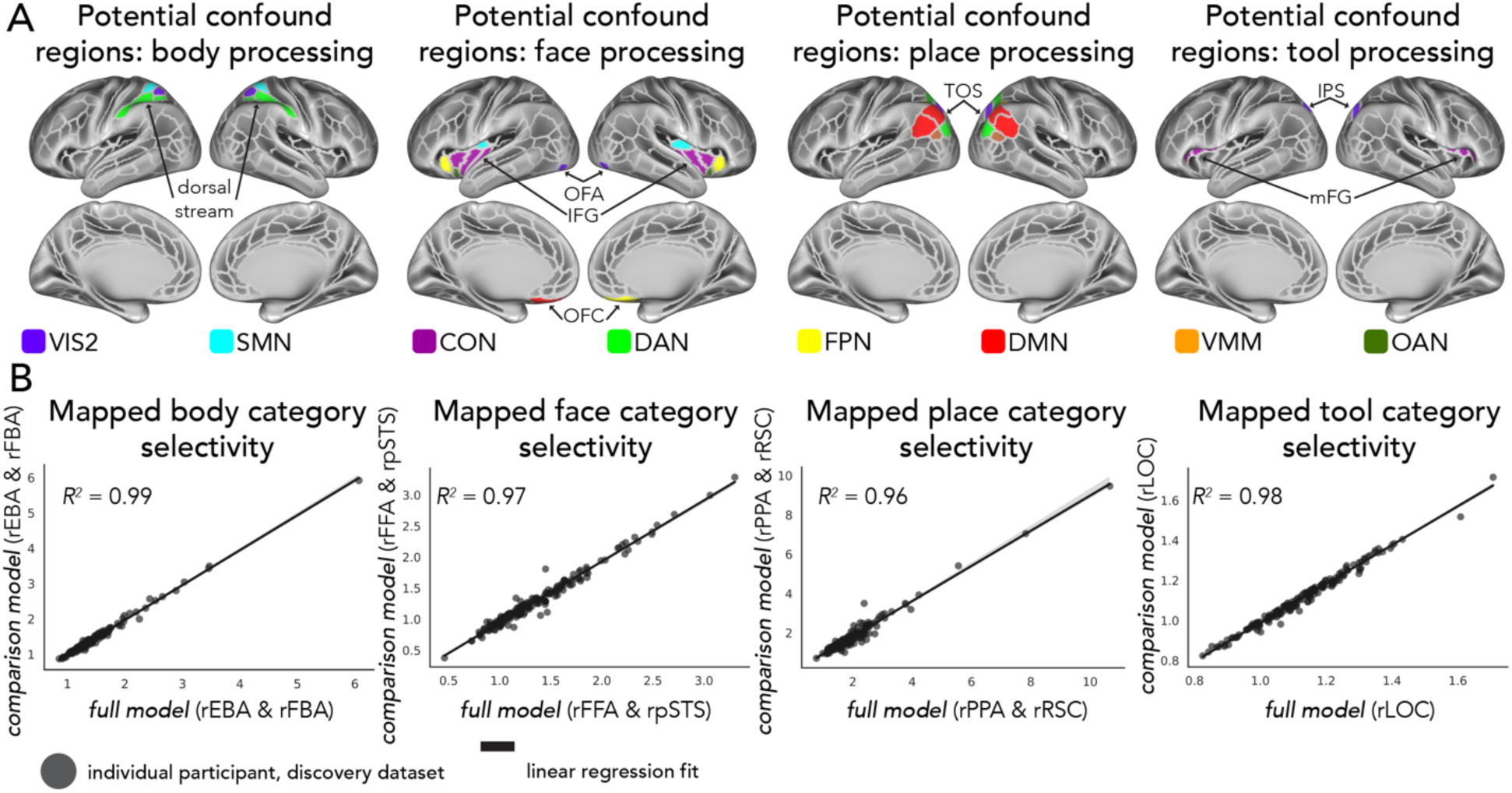
Other regions associated with visual category responses in the literature do not drive the fully-distributed model results. (**A**) While our procedure for identifying functional complexes from the literature was systematic (see Methods), it was possible that excluded, category-responsive regions were driving the results in Figs. 5-8. Such regions were excluded from functional complexes because they were either: (1) not supported by 10 or more peer-reviewed studies at the time of preparing this manuscript, (2) studies used experimental stimuli too distinct from the n-back visual categories, or (3) studies did not provide spatial information systematically consistent with standard volumetric and/or surface-based topography. These regions, held-out from source sets in this control analysis, were as follows (from left to right in panel A): the dorsal visuomotor stream for body selectivity; the occipital face area (OFA), inferior frontal gyrus (IFG), and orbitofrontal cortex (OFC) for face selectivity; the transverse occipital sulcus (TOS) for place selectivity; the intraparietal sulcus (IPS) and medial frontal gyrus (mFG) for tool selectivity. Colors of regions are consistent with the functional network assignment used throughout all other figures. (**B**) For each body, face, place, and tool (left to right) selectivity analysis, individual participant’s (dots) visual selectivity scores were maintained between the results in Figs. 5-8 (x-axes) and the same analyses with the regions in panel A held-out from the source set. This suggests that other category responsive regions in the literature did not drive the fully-distributed model findings.

### Stimulus-driven activity flow processes directly from V1 are sufficient for generating category selectivity

We next sought to test the refined hypothesis that stimulus-driven activity flowing over a functional complex’s connectivity fingerprint with V1 only is sufficient to generate localized visual category selectivity (Figs 1B and 10A). This does not negate the “fully distributed” hypotheses tested above (Fig 1A), but instead specifies that processes instantiated in V1 are key to our prior observations that fully distributed activity flows are sufficient for generating localized visual category selectivity. Alternatively, it is possible that network interactions with V1 account for only a small proportion of variance in visual category selectivity generated by the fully distributed network interaction models (Figs 5-8). For example, it remains possible that the category selectivities observed in the fully distributed model were generated via (1) local (e.g., within-region) computations in regions outside the functional complexes, which (2) together flow into the functional complexes, concentrating selectivity within the complexes. Another possibility is that visual category selectivity is only minimally influenced by direct stimulus-driven inputs from V1, given evidence that visual information processing (which may not be strictly hierarchical; see [164, 165]) is modulated by attention [166–168], prior expectations [169], task goals/context [170–172], and synthesis-related processes (i.e., continuously maintaining a percept from a noisy or ambiguous visual scene) [173–175]. However, it is unclear if these top-down modulations (and other non-visual contributions, such as emotional modulation; as in [176]) are indeed just modulations, or requirements for visual category selectivity.

**Fig 10.**
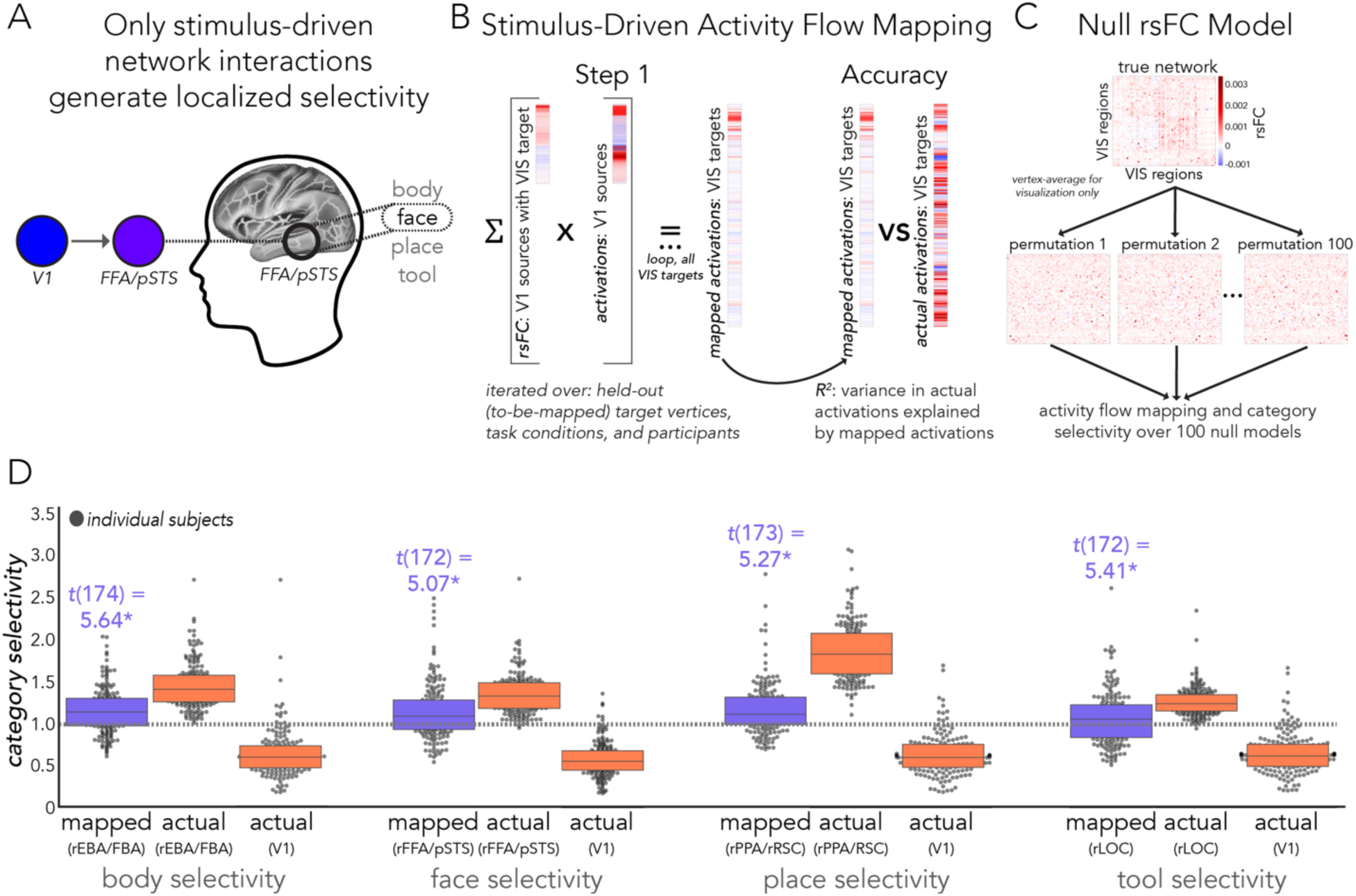
Activity flows directly from V1 are sufficient to generate visual category selectivity. (**A**) Theoretical schematic of stimulus-driven activity flow processes generating visual category selectivity (as in Fig 1B). Given prior literature, mapped activity flow processes (gray arrow) have a refined inference: from V1 to later visual regions, we inferred that activity flow processes were primarily stimulus driven. This contrasts with the prior whole-cortex models, which also included top-down and likely recurrent influences. (**B**) Activity flow mapping procedure for the stimulus-driven model, conducted at the vertex level. V1 sources were used to map targets across VIS. Note that the usage of “step 1” serves as a prelude to later steps tested in an extended visual system model (Fig 11). (**C**) The null connectivity fingerprint (rsFC) model used for control analyses. The top depicts the true VIS network, and the bottom depicts pseudo-random (edge degree and strength preserved) network architectures over 100 permutations. For visualization purposes, networks are shown at the region level, but analyses were conducted at the vertex level. (**D**) Actual (coral) and mapped (purple) visual category selectivity exhibited by the right EBA/FBA, FFA/pSTS, PPA/RSC, and LOC (left to right). Category selectivity exhibited by V1 (for each respective image category) is shown to demonstrate that activation patterns in V1 alone do not account for mapped visual category selectivities (V1 selectivity scores were all nonsignificant; see main text for statistics). Most importantly, these results demonstrate that the activity flow mapping process increased the category selectivity of every functional complex relative to V1 activity patterns, despite source activity originating solely from V1. Category selectivity generated by V1-initialized activity flow processes were significant in each functional complex (see main text for statistics). Significant t-statistics are indicated with an asterisk (p<0.00001; see Methods); see Figs. 5-8 for significance of actual category selectivity of functional complexes.

We additionally sought to address a theoretical limitation borne out of whole-cortex, undirected network estimation from fMRI data. Namely, a limited ability to isolate whether activity flow processes contributing to a given complex’s category-selective response also contain some information from that complex itself, or from other brain regions responsive to visual categories. For example, do fully distributed activity flows generating FFA/pSTS face selectivity (Fig 6) contain some information (potentially via feedback from areas downstream of FFA/pSTS) from the FFA/pSTS itself? This type of causal circularity is a theoretical concern for the ongoing goal of delineating the extent that network interactions contribute to the generation of localized processes with refined detail. However, given that regions within each functional complex (e.g., the FFA and pSTS for the FFA/pSTS complex) were excluded as activity flow sources, the most likely culprits of this sort of feedback circularity were already accounted for. Thus, it is likely that the fully distributed network models (Figs 5-8) accurately reflect the generation of category selectivity, just at a broad scale and with potential inferential limits.

To these ends, we developed an intrinsic-connectivity-based, generative model of category-selective responses that reduces the impact of (potential) causal circularity by constraining activity flows to V1 (sources) and functional complexes (targets) (Figs 10A and 10B). Given robust evidence that activity in V1 represents retinotopic mapping and simple visual feature detection [40,41], this model better ensures that mapped activation patterns are primarily stimulus-driven. We first estimated rsFC between all vertices in the visual system (including both the VIS1 and VIS2 networks and all functional complexes) (Fig 10C, top). We will henceforth refer to this sub-system as “VIS”. In the following steps of this analysis, we focused on the interactions between V1 sources and functional complex targets, but it was important to include the entire visual system in the initial rsFC estimation step. This allowed us to infer – based on our use of multiple regression for rsFC estimation – the likely direct connectivity between each V1 vertex and each functional complex vertex [27, 125].

We found that for each functional complex, stimulus-driven, activity-flow-mapped category selectivity was significantly above 1.0 (*p* < 0.00001 for all complexes; see Methods): right hemisphere EBA/FBA mean body selectivity = 1.12, *t*(174) = 5.64, Cohen’s *d* = 0.45); right FFA/pSTS mean face selectivity = 1.14, *t*(172) = 5.07, Cohen’s *d* = 0.41); right PPA/RSC mean place selectivity = 1.13, *t*(173) = 5.27, Cohen’s *d* = 0.42); right LOC mean tool selectivity = 1.07, *t*(172) = 5.41, Cohen’s *d* = 0.3) (Fig 10D; left hemisphere and replication dataset: S10 Table). In each of these four models, mapped category selectivity was statistically significant, suggesting direct activity flows from V1 (the primary visual input to cortex) are sufficient to produce selectivity for these four visual categories. Note, however, that selectivity in these four models was lower than when mapped via network interactions of the whole cortex. This suggests that additional selectivity is generated via distributed activity flow processes over the whole cortex (Figs 5C-8C).

As in prior control analyses (Table 3), we tested whether each functional complex’s unique resting-state connectivity fingerprint (here, with V1) determined its visual category selectivity by using a null rsFC model. Note that V1 itself did not exhibit significant selectivity for any of the four visual categories (V1 body selectivity = 0.63, *t*(175) = -17.3, *p* = 1, Cohen’s *d* = -1.32; face selectivity = 0.87, *t*(175) = -12.9, *p* = 1, Cohen’s *d* = -0.99; place selectivity = 0.64, *t*(175) = -13.6, *p* = 1, Cohen’s *d* = -1.04; tool selectivity = 0.66, *t*(175) = -12.8, *p* = 1, Cohen’s *d* = -0.97) (Fig 10D; replication dataset: S11 Table), thus downstream selectivity could not be driven by activation patterns in V1 without additional connectivity-based transformation (also supported by [113,177–179]). Given the varied number of vertices in each complex, we used randomly permuted connectivity architectures (Fig 10C) – maintaining edge strength and degree [134,150,151] – instead of rsFC substitution. Body, face, and place selectivity were significantly greater than the aggregate of 100 null rsFC models (body: t(175) = 6.4, *p* = 6.6 x 10^-10^, Cohen’s *d* = 0.49; face: t(175) = 11.6, *p* = 8.2 x 10^-24^, Cohen’s *d* = 0.89; place: t(175) = 3.5, *p* = 2.5 x 10^-4^, Cohen’s *d* = 0.27), but not tool selectivity (t(175) = -1.15, *p* = 0.8, Cohen’s *d* = -0.09) (left hemisphere and replication dataset: S12 Table). This suggests that stimulus-driven category selectivity exhibited by EBA/FBA, FFA/pSTS, and PPA/RSC were significantly shaped by their unique connectivity patterns with V1.

These results suggest that direct stimulus-driven activity flows from V1 are a key step in the generation of visual category selectivity in visual cortex (Fig 1B). For some functional complexes, such as the LOC, early information patterns across V1 can select category-selectivity-generating activity flow processes in a manner less dependent on its connectivity fingerprint (but note that LOC’s connectivity fingerprint with V1 appears more critical in the whole-cortex model; Table 3). For other complexes such as the FFA/pSTS, the stimulus-driven account of category selectivity appears to be influenced to a large extent by its connectivity fingerprint with V1, given its strong statistical significance over the null rsFC model.

### Multi-step activity flows across the visual system improves response profile accuracies, but not category selectivity

Given evidence that higher-level visual representations (such as category selectivity) involve information processing along the ventral visual stream (Fig 11A) [86,180,181], we sought to extend our stimulus-driven model and test the hypothesis that category selectivity mappings (which were significant but below empirically-observed levels in the V1 model) improve when accounting for all VIS network interactions (Fig 1C). We conducted this analysis using the mapped activation patterns from the stimulus-driven V1 model (Figs 10B and 11B) – weighted by rsFC across VIS – to model later visual cortex activity flow processes (Fig 11B, Step 2). This step was repeated to model potential bidirectional (i.e., reciprocal) and/or recurrent processes within the visual system [165,181] (Fig 11B, Steps 3+), until a settling threshold was reached (i.e., where mapped activation patterns no longer changed, Fig 11C; also see Methods).

**Fig 11.**
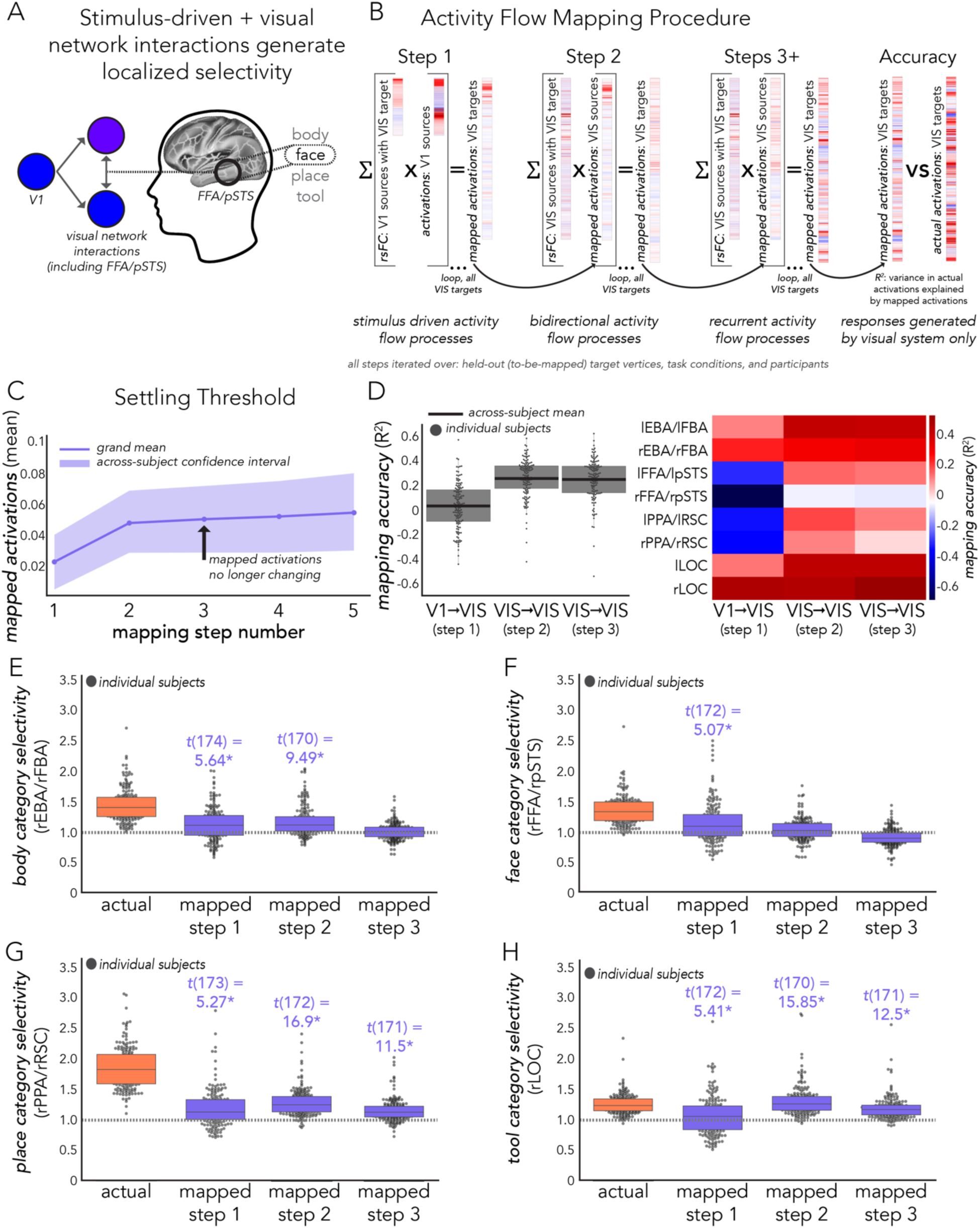
Visual category selectivity generated by activity flow processes over the extended visual system. (**A**) Stimulus driven activity flow processes, further shaped by later visual interactions, generate localized visual category selectivity (Fig 1C). From V1 to later visual regions, we inferred that activity flow processes are stimulus driven (Figs 1B and 10). Within the visual system, we inferred that activity flow processes are bidirectional and/or recurrent. (**B**) Activity flow mapping procedure for the extended visual system model. All steps were conducted at the vertex level. ***Step 1***: V1 sources were used to map targets across the visual cortex (VIS) (Fig 10). ***Step 2***: mapped VIS activation patterns from Step 1 (weighted by connectivity estimates as in all activity flow models; see Methods) were used as sources to map held-out targets across visual cortex. ***Steps 3+***: step 2 was repeated until a settling threshold was reached – or the point at which mapped values stopped changing (see Methods). (**C**) The settling threshold was reached at step 3. All further analyses only included steps 1-3 from panel B. (**D**) Accuracy of mapped activation patterns across all conditions (left: an average of all visual regions; right: functional complexes studied herein). Across visual cortex, explained variance tended to increase with each step, indicating that the extended visual system model was improving mapped response profiles across the cortex. This pattern was also observed across functional complexes, with some exceptions, such as the PPA/RSC (which appears most accurate at step 2). (**E-H**) Actual (coral) and visual-system-mapped (purple) category selectivity (see Methods) exhibited by the right EBA/FBA (E), FFA/pSTS (F), PPA/RSC (G), and LOC (H). Mapped category selectivity is shown for each step in panel B. Activity-flow-mapped category selectivity estimates were not consistently significant (see main text for statistics), suggesting that extending V1-initialized activity flow processes across the visual system alone does not stably generate localized visual category selectivity. In panels E-F: significant t-statistics are indicated with an asterisk (p<0.00001; see Methods); see Figs. 5-8 for significance of actual category selectivity of functional complexes.

Cross-condition (i.e., response profile) mapping accuracy (of all VIS vertices) increased with each step (Fig 11D), suggesting that this extended model improved the specification of visual system responses. However, category selectivity was not improved consistently across functional complexes of interest. Mapped body selectivity in the right EBA/FBA was improved in step two (body selectivity = 1.18, *t*(170) = 9.49, *p* < 0.00001, Cohen’s *d* = 0.75) but became nonsignificant in step three (body selectivity = 1.01, *t*(174) = 0.58, Cohen’s *d* = 0.03) (Fig 11E). Face selectivity diminished in step 2 (face selectivity = 1.03, *t*(174) = 1.82, not significant, Cohen’s *d* = 0.14) and step 3 (face selectivity = 0.9, *t*(167) = -9.23, not significant, Cohen’s *d* = -0.71) (Fig 11F), suggesting that within the confines of the visual system, stimulus-driven activity flows from V1 (step 1) were the best predictors of right FFA/pSTS face selectivity. Place selectivity in the right PPA/RSC improved in step 2 (place selectivity = 1.28, *t*(172) = 16.9, *p* < 0.00001, Cohen’s *d* = 1.43), but decreased to the original level in step 3 (place selectivity = 1.15, *t*(171) = 11.5, *p* < 0.00001, Cohen’s *d* = 0.88) (Fig 11G). Tool selectivity in the right LOC improved in step 2 (tool selectivity = 1.28, *t*(170) = 15.85, *p* < 0.00001, Cohen’s *d* = 1.5), but decreased in step 3, albeit to a level higher than the initial V1 step (tool selectivity = 1.16, *t*(171) = 12.5, *p* < 0.00001, Cohen’s *d* = 1.23) (Fig 11H; left hemisphere and replication dataset: S13 and S14 Tables). Interestingly, tool selectivity generated by step 2 was greater than the actual tool selectivity exhibited by the LOC (Fig 11H, coral), suggesting that extended visual system modulations to activity flow processes initially over-specified tool selectivity, and that step 3 modeled the best-performing network interactions for mapping LOC tool selectivity. Thus, for two complexes (EBA/FBA and PPA/RSC), one additional step of extended visual system network interactions best specified category selectivity. For one complex (FFA/pSTS), the V1 stimulus-driven (step 1) model was best, and for another complex (LOC) three steps were best.

Therefore, even though mapped response profiles tended to improve when modeling bidirectional and/or recurrent processes across the visual system, visual-system-mapped category selectivities did not consistently improve beyond what was specified by stimulus-driven activity flows from V1. Additionally, given that all observations were most robust in the fully distributed model (Fig 1A) (including response profile accuracies, which explained a maximum of 45% variance in the extended visual model, Fig 11D, and up to 92% in the whole-cortex model, Fig 3D), the visual system is likely further modulated by other systems (e.g., DAN interactions; also supported by Keane et al. [182]) to provide the full set of distributed activity flow processes that stably generate localized category selective responses (as in Figs 5-8).

### Adding fully distributed network interactions to stimulus-driven activity flows further enhances category selectivity

Given evidence that stimulus-driven network interactions with V1 generated visual category selectivity (Fig 10), but to a lesser extent than the fully distributed mappings (Figs 5-8), and that this inconsistently improved when extended to later visual system interactions (Fig 11), we next sought to extend the stimulus-driven model to fully distributed network interactions (i.e., a “stimulus-driven + fully distributed” mapping) (Figs 1D and 12A). As in prior analyses, fully distributed network interactions refer to activity flow processes over all cortical source regions (Formula 1). Note that the stimulus-driven + fully distributed model differs from the initial cortex-wide model (Fig 1A) by constraining inputs to V1 – reducing the chance for causal circularity (e.g., from recurrent feedback to each functional complex) and excluding possible temporally extended activity (in non-V1 regions) originating prior to stimulus onset (e.g., attentional or task set top-down biases). First, using the extended mapping approach as in Fig 11B, we initialized activity flow mapping with V1 network interactions, then incorporated all other network interactions in a second step (i.e., with regions beyond just VIS, see Methods for full details). Across all functional complexes, we found that activity-flow-mapped category selectivity was not only statistically significant but was also remarkably close to selectivity generated by fully distributed network interactions (Figs 5-8), and in some cases exhibited an improvement. Further, selectivity was significantly improved for all functional complexes relative to the stimulus-driven V1-initiated model, demonstrating the importance of additional brain-wide activity flows in increasing category selectivity beyond direct-from-V1 activity flows.

**Fig 12.**
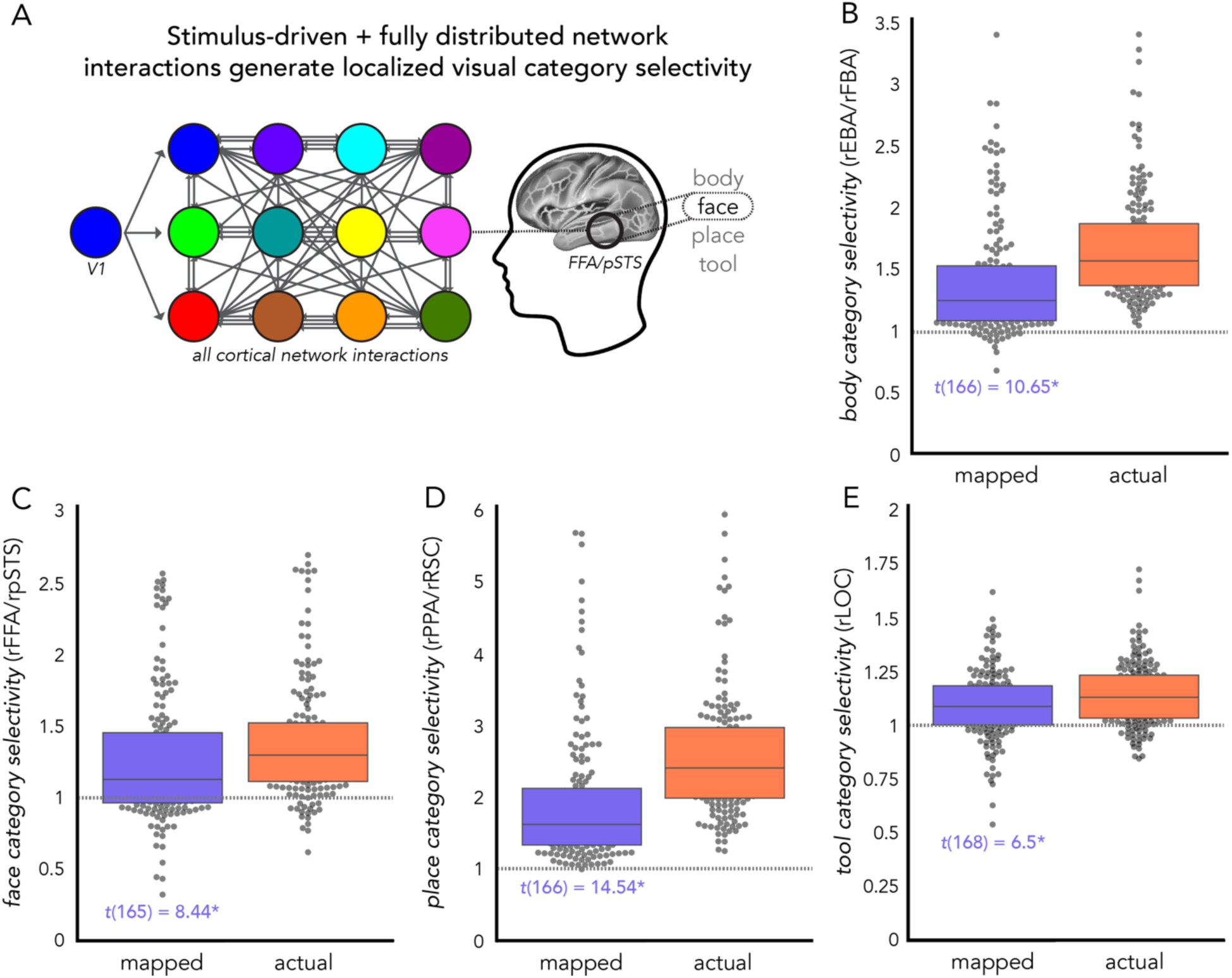
Adding fully distributed network interactions to stimulus-driven activity flows further enhances category selectivity. (**A**) A schematic of V1-initialized activity flow processes that are further propagated across all cortical network interactions (as in Fig 1D). Here, fully distributed network interactions are initially established by activity flowing over the connectivity fingerprints of each functional complex with V1. We used the multistep mapping procedure of Fig 11B (but with only two steps), except for at the region level (Methods). (**B**) Mapped (purple) and actual (coral) body selectivity exhibited by the right hemisphere EBA/FBA (gray dots: individual participants; boxplot line: median). Statistical significance is reported in the main text. (C-E) Same as B, but: (**C**) face selectivity in FFA/pSTS, (**D**) place selectivity in PPA/RSC, and (**E**) tool selectivity in LOC. Across all functional complexes, stimulus-driven + fully distributed mappings generated visual category selectivity that was greater than the stimulus-driven alone model of Fig 10 and closely matched (and in some cases improved upon) the fully distributed alone model of Figs 5-8. This suggests that activity flow processes initialized in V1, that are further processed over all cortical network interactions, are capable of generating highly accurate visual category selectivity, with reduced causal confounds (see main text for full rationale). Panels B-D: significant t-statistics are indicated with an asterisk (*p*<0.00001; see Methods); see Figs. 5-8 for significance of actual category selectivity of functional complexes.

Body selectivity in the EBA/FBA was statistically significant (mapped body selectivity = 1.41, *t*(166) = 10.65, Cohen’s *d* = 0.83, *p* < 0.00001; Fig 12B). Here, activity-flow-mapped body selectivity generated by stimulus-driven + fully distributed network interactions was an improvement upon selectivity generated from V1 only (Fig. 10) (related samples t-tests across participants’ activity-flow-mapped body category selectivity scores: *t*(175) = 4.75, *p* = 4.18 x 10^-6^). Category selectivity in the FFA/pSTS (mean mapped mean face selectivity = 1.26, *t*(165) = 8.44, Cohen’s *d* = 0.57, *p* < 0.0000; Fig. 12C), the PPA/RSC (mapped place selectivity = 1.94, *t*(166) = 14.54, Cohen’s *d* = 1.0, *p* < 0.00001; Fig 12D), and the LOC (mapped tool selectivity = 1.09, *t*(168) = 6.5, Cohen’s *d* = 0.58, *p* < 0.00001; Fig 12E) were all also significantly improved from the stimulus-driven (V1 only) mappings (faces: *t*(175) = 3.41, *p* = 8.04 x 10^-4^; places: *t*(175) = 8.41, *p* = 7.05 x 10^-15^; tools: *t*(175) = 2.16, *p* = 0.03).

## Discussion

The foregoing results suggest that visual category selectivity in visual cortex is primarily generated by distributed activity flowing over intrinsic functional connections (Figs 5A-8A), even when activity flows were restricted to stimulus-driven network interactions (i.e., excluding top-down task set influences and/or influences from other category-responsive regions) (Fig 12). We observed significant visual category selectivity in four functional complexes for both actual responses and responses mapped via fully distributed activity flow processes (Figs 1A and 5C-8C). Further, distributed network interactions accounted for the majority of variance in visual category selectivity (Figs 5D-8D). VIS2 interactions most prominently contributed to both category-specific and response profile activations across complexes, with additional contributions by other networks for select models: body-EBA/FBA activity from DAN; face-FFA/pSTS from DAN and DMN; and place-PPA/RSC from DAN, VIS1, and DMN (Figs 5E-F-8E-F). Null network architectures revealed that each region’s unique intrinsic connectivity fingerprint was a key component in enabling activity flow mapping to generate category selectivity. Likewise, when source sets excluded regions otherwise known to be responsive to visual categories, the fully-distributed activity flow processes remained capable of generating category selectivity. Lastly, we found that V1-initialized network interactions (Figs 1B-D) were sufficient to generate category selectivity (Fig 10), with additional increases in category selectivity from fully distributed network interactions (Figs 1D and 12) . Altogether, these findings support the hypothesis that distributed activity flow processes, specified by intrinsic connectivity fingerprints of visual cortex regions, chiefly shape their category selective responses.

These findings build upon prior observations of distributed and overlapping representations in the visual cortex [11], as well as the general hypothesis that connectivity fingerprints determine a region’s functional repertoire [13,15]. Results here also extend observations that structural connectivity fingerprints (of the same regions studied here) can be used to predict visual category responses [21], as well as observations that responses to visual stimuli can be predicted by the spatial topography of activity in other visual cortex regions [14]. Here we go beyond these predictive results by modeling mechanistic processes – localized convergence of activity flows over intrinsic connectivity fingerprints [27,28] – contributing to the generation of localized activations [20]. We focused on visual regions here so we could leverage the long-standing observations of category selectivity in visual cortex regions to test various network interaction models, revealing how localized activations are (at least for the regions tested) predominantly generated by distributed activity flow processes shaped by intrinsic functional connectivity fingerprints.

Importantly, fully distributed activity flow mappings (Figs 5-8) did not account for all task-evoked activity variance, leaving room for within-region mechanisms to carry out critical computations. Here we adopted a definition of local computation consistent with fMRI and electroencephalography research: local activity reflects local field potentials, which are thought to reflect inputs and within-region processing [183]. Therefore, it was not a guarantee that distributed processes would exhibit the predominant influence in this fMRI dataset. However, the present study provides two major lines of evidence in support of this claim. Firstly, fully distributed activity flow processes mapped visual category responses with high accuracy in all functional complexes. If within-region mechanisms strictly accounted for these responses we would expect lower accuracy, given prior simulation-based findings demonstrating that successful activity flow mapping requires high distributed processing (global coupling) and low local processing (self-coupling) (Fig 2B [27,28]). Mapped category selectivity within the fully distributed model (Fig 1A) is thus a proxy for the degree of distributed processes underlying category selectivity. Secondly, the estimated contribution of distributed processes to category selectivity was significantly greater than 50% for all functional complexes assessed. We additionally observed that activity flow processes explained the majority of variance in response profiles (across 24 diverse task conditions) exhibited by each complex (Figs 5F-8F), suggesting that activity flowing over global, intrinsic connectivity patterns is a general mechanism in the emergence of fMRI responses. However, large-scale functional networks had differential influence depending on the cognitive domain and target complex. For example, LOC responses exhibited a strong general influence from VIS2 interactions (Figs 8E-F); in contrast to FFA/pSTS, which exhibited strong VIS2 influence in the face-specific model (Fig 6E) with additional influence from DAN and DMN in the general response profile model (Fig 6F).

Given this, along with the well-known observation that visual inputs from the retina arrive in cortex (by way of the lateral geniculate nucleus) in V1 [40,41], we adapted activity flow mapping to test refined hypotheses about the possible stimulus-driven network interactions that generate visual category selectivity (Figs 1B-D). We found evidence that when solely initialized by V1 source regions, activity flow mappings generated significant (though less than the fully distributed model) category selectivity (Figs 1B and 10). This demonstrated that category selectivity is generated initially by activity flows directly from V1. Category selectivity inconsistently improved when mappings were further shaped by visual system interactions (Figs 1C and 11), but markedly (and consistently) improved when further shaped by fully distributed cortical network interactions (Figs 1D and 12). These results suggest that fully distributed network interactions that were initially established in V1 are sufficient to generate visual category selectivity (without, e.g., local computations). This approach allowed for improved inferential validity by largely ruling out a potential causal confound wherein the non-V1-initiated fully distributed model (Fig 1A) interactions might receive information from the functional complex itself through feedback. It is possible that an extended step of fully distributed interactions outperformed extended steps of visual system interactions because of the influence of top-down modulations thought to be critical for visual processing [166–168,170,171]. We encourage future work that systematically tests which non-visual-network interactions are most critical to further shaping stimulus-driven activity flow processes.

The present results support a framework of neurocognitive processing that emphasizes the mechanistic influence of distributed network interactions, even in the extreme case of localized visual category selectivity. This follows from our demonstrations that category selectivity in functional complexes can be generated via an empirically-estimated distributed processing model: task-evoked activity flowing over connectivity fingerprints. However, while our resting-state connectivity estimates address causal confounds better than standard FC measures (Fig 3A), they lack directional information. For instance, if ongoing within-region process variance strongly influenced downstream regions, then this distributed output activity would contribute (in the incorrect causal direction) to the mapping of the localized processes. This possibility is unlikely, however, since such strongly category selective output would likely drive downstream regions to also exhibit robust category selectivity, and the best-established category selective regions were included in each functional complex in our analyses, removing them from prediction sources. We also conducted a control analysis that specifically excluded other known visual-category-responsive regions from source sets, and results were remarkably consistent. In addition, we more systematically ruled out this possibility for circularity in the stimulus-driven network interaction models (Figs 1B-D, see Methods) by restricting all activity flows to those originating (directly or indirectly) from visual input region V1. Altogether, our results largely rule out the alternative hypothesis that localized visual category selectivity is solely driven by within-region computations. For instance, our results rule out the hypothesis that elevated face-image-evoked activity in FFA solely reflects within-FFA computations to detect or identify faces. Instead, the present results suggest elevated activity in FFA primarily reflects connectivity-specified activity flows summing together weakly category-selective distributed responses to generate highly category-selective localized responses. It will be important for future research to disentangle input and output processes to and from visual category selective brain regions in order to further specify the time course and directionality of network interactions that support visual category selectivity. One compelling approach that aligns with the broad inferences made in the present study is connective field modeling [14], which predicts responses in visual cortex regions from activity patterns in other brain regions. This approach does not implement functional and/or structural connectivity estimation standard to neuroimaging, but instead a spatially-linked Gaussian model-fit procedure that future researchers may integrate as part of the activity flow mapping to develop estimates of direction of information flow. Additionally, following suggestions by Poldrack et al. [61] and the accessibility of functional regions assessed in Osher et al. [21], we selected functional complexes of interest *a priori* and uniformly across participants. However, we encourage future work to probe individual differences in category-selective topography (as in face localizer tasks [184], or with individualized network parcellations [185]). More broadly, in order to formalize a rigorous testbed for our hypotheses, we treated the end point of activity-flow-mapped visual category selectivity as a localized response. However, future work that builds upon the present study will likely make greater contact with neurobiological reality if dispersed representation of visual categories across the cortex are probed.

In summary, we observed that distributed network interactions, specified by intrinsic connectivity fingerprints, are likely the primary contributor to the emergence of category selectivity in visual cortex regions. This finding builds upon and integrates a history of theories in vision neuroscience and network neuroscience emphasizing the importance of distributed and overlapping activation patterns [11], connectivity fingerprints specifying regional functioning [13,15], and models predicting visual responsiveness [14]. We leveraged activity flow mapping’s inherent sensitivity to global processing features [27] to estimate the contribution of distributed processes upon visual category selectivity. Looking forward, the present findings can facilitate examinations of the generative capacity of distributed neural processing mechanisms by constraining and contextualizing hypotheses.

## Acknowledgements

Data were provided [in part] by the Human Connectome Project, WU-Minn Consortium (Principal Investigators: David Van Essen and Kamil Ugurbil; 1U54MH091657) funded by the 16 NIH Institutes and Centers that support the NIH Blueprint for Neuroscience Research; and by the McDonnell Center for Systems Neuroscience at Washington University. We thank the Office of Advanced Research Computing (OARC) at Rutgers, The State University of New Jersey for providing access to the Amarel cluster and associated research computing resources. We also thank our colleagues at the Cole Neurocognition Lab for offering words of wisdom and overall support.

## Supporting Information Legends

**S1 Fig. Whole-cortex activity-flow-mapped activations for four semantic visual categories.** This assessment demonstrates that activity flow mapping of cross-cortex activations to each visual semantic category exhibits high accuracy. (**A**) Left: Cross-participant average actual (empirical) task activations to body categories projected onto the MMP cortical atlas [54]. Right: Cross-participant average activity-flow-mapped task activations to body categories projected onto the MMP cortical atlas. The mapped and actual activations exhibited a high degree of overlap: *r* = 0.89. (**B-D**) The same as in A, but for face (B), place (C), and tool categories (D) respectively. In all cases, accuracy was high, demonstrating that activity flow processes mapped predicted cross-cortex responses to each visual category of interest well.

**S2 Fig. Benchmarking activity in four functional complexes.** To corroborate findings in the literature that each of the four functional complexes assessed in the present study exhibit significantly higher activations to images in their respective visual categories, we conducted standard t-test contrasts (discovery dataset, right hemisphere results depicted here; left hemisphere statistics reported in S1 Table). (**A**) Box and swarm plots depicting activations to images of bodies and body parts exhibited by the EBA/FBA (dots = individual participant’s data). The actual activity is shown in coral and the activity-flow-mapped activity is shown in purple. Body versus non-body activations were contrasted with a one-tailed, paired samples t-test (hypothesizing that body activity was larger than non-body activity in the EBA/FBA), with an asterisk indicating a statistically significant difference (*p* < 0.0001 with nonparametric permutation tests, as reported in the main text Results). The EBA/FBA exhibited statistically greater activations to body versus non-body images, benchmarking the prior work in the literature. (**B**) Same as in A, but for face versus non-face images and the FFA/pSTS functional complex. (**C**) Same as in A, but for place versus non-place images and the PPA/RSC functional complex. (**D**) Same as in A, but for tool versus non-tool images and the LOC functional complex. All results were corroborated by the replication dataset (*N*=176 in each; statistics given in main Results text).

**S1 Table. Whole-cortex activity-flow-mapped responses to visual category conditions.** Across select portions of cortex (whole cortex = all 360 MMP cortical regions [54]) and all *n*=176 participants, the accuracy of activity flow mapping was estimated by comparing of mapped and actual responses to select HCP conditions (response profile = across all 24 HCP conditions) via Pearson’s *r*, mean absolute error (MAE), and the coefficient of determination (R^2^).

**S2 Table. Benchmark contrasts in four functional complexes.** Number of permutations in max-T nonparametric permutation testing: 10,000. Disc. = discovery; repl. = replication; hemi. = hemisphere.

**S3 Table. Category selectivity scores and estimated percent distributed processing contribution to category selectivity in four functional complexes.** Number of permutations in max-T nonparametric permutation testing: 100,000. *d.f.* = degrees of freedom (see Methods for details on outlier removal procedure). Disc. = discovery; repl. = replication; thresh. = threshold; hemi. = hemisphere.

**S4 Table. Large-scale functional network activity flows contributing to category-specific responses in four functional complexes.** Number of permutations in max-T nonparametric permutation testing: 10,000. sig. = significant. VIS1 = primary visual network; VIS2 = secondary visual network; DAN = dorsal attention network. These results corroborate results presented in Figs 4E-7E (right hemisphere discovery data; statistics reported in main text). Disc. = discovery; repl. = replication; thresh. = threshold; hemi. = hemisphere.

**S5 Table. Discovery dataset: variance explained per network in predicting cross-condition response profiles in left hemisphere complexes.** Source network = network-based source of explained variance in activity-flow-mapped activations across 24 conditions (i.e., the response profile). VIS1 = primary visual network; VIS2 = secondary visual network; SMN = somatomotor network; CON = cingulo-opercular network; DAN = dorsal attention network; LAN = language network; FPN = frontoparietal network; AUD = auditory network; DMN = default mode network; PMM = posterior multimodal network; VMM = ventral multimodal network; OAN = orbito-affective network. rel. % = percent of relative importance to the full model. Asterisks = statistically significant network contributions (*p* < 0.0001, number of permutations = 10,000). EBA/FBA max-T(175) = 3.41; FFA/pSTS max-T(175) = 3.42; PPA/RSC max-T(175) = 3.42; LOC max-T(175) = 3.39. Statistical results listed in the bottom two rows refer to 1 sample t-testing of the total R^2^ value for each model versus 0.5, which assesses whether the mapped response profile for a given functional complex explains more than 50% of the variance in the actual response profile. This provides evidence that distributed processes (as captured by activity flow mapping) are the dominant influence in generating a given functional complexes activations to a diverse set of cognitive domains. n/a = not applicable. These results corroborate results presented in Figs 4F-7F (right hemisphere discovery data; statistics reported in main text).

**S6 Table. Replication dataset: variance explained per network in predicting cross-condition response profiles in left hemisphere complexes.** Source network = network-based source of explained variance in activity-flow-mapped activations across 24 conditions (i.e., the response profile). VIS1 = primary visual network; VIS2 = secondary visual network; SMN = somatomotor network; CON = cingulo-opercular network; DAN = dorsal attention network; LAN = language network; FPN = frontoparietal network; AUD = auditory network; DMN = default mode network; PMM = posterior multimodal network; VMM = ventral multimodal network; OAN = orbito-affective network. rel. % = percent of relative importance to the full model. Asterisks = statistically significant network contributions (*p* < 0.0001, number of permutations = 10,000). EBA/FBA max-T(175) = 3.34; FFA/pSTS max-T(175) = 3.4; PPA/RSC max-T(175) = 3.35; LOC max-T(175) = 3.38. Statistical results listed in the bottom two rows refer to 1 sample t-testing of the total R^2^ value for each model versus 0.5, which assesses whether the mapped response profile for a given functional complex explains more than 50% of the variance in the actual response profile. This provides evidence that distributed processes (as captured by activity flow mapping) are the dominant influence in generating a given functional complexes activations to a diverse set of cognitive domains. n/a = not applicable. These results corroborate results presented in Figs 4F-7F (right hemisphere discovery data; statistics reported in main text).

**S7 Table. Replication dataset: variance explained per network in predicting cross-condition response profiles in right hemisphere complexes.** Source network = network-based source of explained variance in activity-flow-mapped activations across 24 conditions (i.e., the response profile). VIS1 = primary visual network; VIS2 = secondary visual network; SMN = somatomotor network; CON = cingulo-opercular network; DAN = dorsal attention network; LAN = language network; FPN = frontoparietal network; AUD = auditory network; DMN = default mode network; PMM = posterior multimodal network; VMM = ventral multimodal network; OAN = orbito-affective network. rel. % = percent of relative importance to the full model. Asterisks = statistically significant network contributions (*p* < 0.0001, number of permutations = 10,000). EBA/FBA max-T(175) = 3.41; FFA/pSTS max-T(175) = 3.39; PPA/RSC max-T(175) = 3.42; LOC max-T(175) = 3.38. Statistical results listed in the bottom two rows refer to 1 sample t-testing of the total R^2^ value for each model versus 0.5, which assesses whether the mapped response profile for a given functional complex explains more than 50% of the variance in the actual response profile. This provides evidence that distributed processes (as captured by activity flow mapping) are the dominant influence in generating a given functional complexes activations to a diverse set of cognitive domains. n/a = not applicable. These results corroborate results presented in Figs 4F-7F (right hemisphere discovery data; statistics reported in main text).

**S8 Table. Null connectivity fingerprint models, right hemisphere functional complexes and replication dataset.** In each analysis, the true model category selectivity scores (i.e., activity-flow-mapped with true connectivity fingerprint) were compared to the null model category selectivity scores (i.e., activity-flow-mapped with substituted connectivity fingerprints) across participants (paired samples t-test). Repl. = replication.

**S9 Table. Null connectivity fingerprint models, left hemisphere functional complexes.** In each analysis, the true model category selectivity scores (i.e., activity-flow-mapped with true connectivity fingerprint) were compared to the null model body selectivity scores (i.e., activity-flow-mapped with substituted connectivity fingerprints) across participants (paired samples t-test). Disc. = discovery; repl. = replication.

**S10 Table. Category selectivity generated from stimulus-driven activity flow processes for left hemisphere functional complexes and the replication dataset.** Number of permutations in max-T nonparametric permutation testing: 100,000. *d.f.* = degrees of freedom (see Methods for details on outlier removal procedure). This analysis corresponds to category selectivity generated from V1-initialized, stimulus-driven activity flow processes (Figs 1B and 9; Step 1 of Fig 11B). Disc. = discovery; repl. = replication; hemi. = hemisphere; thresh. = threshold; n.s. = not significant.

**S11 Table. Actual category selectivity exhibited by V1 for the replication dataset.** As in the main text (reporting the discovery dataset results), bilateral V1 (vertex-level data, see Methods) itself did not exhibit significant category selectivity for any of the four visual categories. Given this observation, it is likely that intrinsic connectivity patterns (between V1 and functional complexes of interest), and not source activation patterns in V1, were key to stimulus-driven activity flow processes being able to generate visual category selectivity.

**S12 Table. Stimulus-driven category selectivity generated via activity flow mapping is significantly greater than with a null network architecture.** In each analysis, the true model category selectivity scores (i.e., via stimulus-driven activity flow mapping with true connectivity fingerprint with V1) were compared to the null model (Fig 10C) category selectivity scores (i.e., activity flow mapping with randomly shuffled connectivity fingerprints, see Methods) across participants (paired samples t-test). N.s. = not significant.

**S13 Table. Category selectivity via stimulus-driven processes further shaped by VIS subsystem network interactions (step 2), left hemisphere functional complexes and the replication dataset.** Number of permutations in max-T nonparametric permutation testing: 100,000. *d.f.* = degrees of freedom (see Methods for details on outlier removal procedure). This analysis corresponds to category selectivity generated from V1-initialized, stimulus-driven activity flow processes that are further shaped by one step of VIS subsystem network interactions (Figs 1C and 11). Disc. = discovery; repl. = replication; hemi. = hemisphere; thresh. = threshold; n.s. = not significant.

**S14 Table. Category selectivity via stimulus-driven processes further shaped by VIS subsystem network interactions (step 3), left hemisphere functional complexes and the replication dataset.** Number of permutations in max-T nonparametric permutation testing: 100,000. *d.f.* = degrees of freedom (see Methods for details on outlier removal procedure). This analysis corresponds to category selectivity generated from V1-initialized, stimulus-driven activity flow processes that are further shaped by two steps of VIS subsystem network interactions (Figs 1C and 11). Disc. = discovery; repl. = replication; hemi. = hemisphere; thresh. = threshold; n.s. = not significant.

**S15 Table. Category selectivity via stimulus-driven activity flow processes further shaped by whole-cortex, fully distributed network interactions.** Number of permutations in max-T nonparametric permutation testing: 100,000. *d.f.* = degrees of freedom (see Methods for details on outlier removal procedure). This analysis corresponds to category selectivity generated from V1-initialized, stimulus-driven activity flow processes that are further shaped by fully distributed network interactions (Figs 1D and 12). Disc. = discovery; repl. = replication; hemi. = hemisphere; thresh. = threshold.

## Notes

https://humanconnectome.org/study/hcp-young-adult

### Competing Interest Statement

The authors have declared no competing interest.

### Summary of Updates

We updated the Introduction and Discussion to provide clarity on framing. We added a control analysis (see Fig. 9). Finally, we updated the Methods to provide necessary clarifying details.

## References

1. Borra, E., Ichinohe, N., Sato, T., Tanifuji, M., & Rockland, K. S. (2010). Cortical connections to area TE in monkey: hybrid modular and distributed organization. Cerebral Cortex , 20(2), 257–270.

2. Downing, P. E., Jiang, Y., Shuman, M., & Kanwisher, N. (2001). A cortical area selective for visual processing of the human body. Science, 293(5539), 2470–2473.

3. Taylor, J. C., Wiggett, A. J., & Downing, P. E. (2007). Functional MRI analysis of body and body part representations in the extrastriate and fusiform body areas. Journal of Neurophysiology, 98(3), 1626–1633.

4. Kanwisher, N., & Yovel, G. (2006). The fusiform face area: a cortical region specialized for the perception of faces. Philosophical Transactions of the Royal Society of London. Series B, Biological Sciences, 361(1476), 2109–2128.

5. Park, S., & Chun, M. M. (2009). Different roles of the parahippocampal place area (PPA) and retrosplenial cortex (RSC) in panoramic scene perception. NeuroImage, 47(4), 1747–1756.

6. Peelen, M. V., & Downing, P. E. (2005). Within-subject reproducibility of category-specific visual activation with functional MRI. Human Brain Mapping, 25(4), 402–408.

7. Kanwisher, N. (2010). Functional specificity in the human brain: a window into the functional architecture of the mind. Proceedings of the National Academy of Sciences of the United States of America, 107(25), 11163–11170.

8. Tsao, D. Y., Moeller, S., & Freiwald, W. A. (2008). Comparing face patch systems in macaques and humans. Proceedings of the National Academy of Sciences of the United States of America, 105(49), 19514–19519.

9. Corrow, S. L., Dalrymple, K. A., & Barton, J. J. (2016). Prosopagnosia: current perspectives. Eye and Brain, 8, 165–175.

10. Schalk, G., Kapeller, C., Guger, C., Ogawa, H., Hiroshima, S., Lafer-Sousa, R., Saygin, Z. M., Kamada, K., & Kanwisher, N. (2017). Facephenes and rainbows: Causal evidence for functional and anatomical specificity of face and color processing in the human brain. Proceedings of the National Academy of Sciences of the United States of America, 114(46), 12285–12290.

11. Haxby, J. V., Gobbini, M. I., Furey, M. L., Ishai, A., Schouten, J. L., & Pietrini, P. (2001). Distributed and overlapping representations of faces and objects in ventral temporal cortex. Science, 293(5539), 2425–2430.

12. Finn, E. S., Shen, X., Scheinost, D., Rosenberg, M. D., Huang, J., Chun, M. M., Papademetris, X., & Constable, R. T. (2015). Functional connectome fingerprinting: identifying individuals using patterns of brain connectivity. Nature Neuroscience, 18(11), 1664–1671.

13. Passingham, R. E., Stephan, K. E., & Kötter, R. (2002). The anatomical basis of functional localization in the cortex. Nature Reviews. Neuroscience, 3(8), 606–616.

14. Haak, K. V., Winawer, J., Harvey, B. M., Renken, R., Dumoulin, S. O., Wandell, B. A., & Cornelissen, F. W. (2013). Connective field modeling. NeuroImage, 66, 376–384.

15. Mars, R. B., Passingham, R. E., & Jbabdi, S. (2018). Connectivity Fingerprints: From Areal Descriptions to Abstract Spaces. Trends in Cognitive Sciences, 22(11), 1026–1037

16. Sporns, O. (2014). Contributions and challenges for network models in cognitive neuroscience. Nature Neuroscience, 17(5), 652–660.

17. Friston, K. J. (2011). Functional and effective connectivity: a review. Brain Connectivity, 1(1), 13– 36.

18. Reid, A. T., Headley, D. B., Mill, R. D., Sanchez-Romero, R., Uddin, L. Q., Marinazzo, D., Lurie, D. J., Valdés-Sosa, P. A., Hanson, S. J., Biswal, B. B., Calhoun, V., Poldrack, R. A., & Cole, M. W. (2019). Advancing functional connectivity research from association to causation. Nature Neuroscience, 22(11), 1751–1760.

19. Cole, M. W., Ito, T., Cocuzza, C., & Sanchez-Romero, R. (2021). The Functional Relevance of Task-State Functional Connectivity. The Journal of Neuroscience: The Official Journal of the Society for Neuroscience, 41(12), 2684–2702

20. Ito, T., Hearne, L., Mill, R., Cocuzza, C., & Cole, M. W. (2020c). Discovering the Computational Relevance of Brain Network Organization. Trends in Cognitive Sciences, 24(1), 25–38.

21. Osher, D. E., Saxe, R. R., Koldewyn, K., Gabrieli, J. D. E., Kanwisher, N., & Saygin, Z. M. (2016). Structural Connectivity Fingerprints Predict Cortical Selectivity for Multiple Visual Categories across Cortex. Cerebral Cortex , 26(4), 1668–1683.

22. Biswal, B., Yetkin, F. Z., Haughton, V. M., & Hyde, J. S. (1995). Functional connectivity in the motor cortex of resting human brain using echo-planar MRI. Magnetic Resonance in Medicine: Official Journal of the Society of Magnetic Resonance in Medicine / Society of Magnetic Resonance in Medicine, 34(4), 537–541.

23. Fox, M. D., & Raichle, M. E. (2007). Spontaneous fluctuations in brain activity observed with functional magnetic resonance imaging. Nature Reviews. Neuroscience, 8(9), 700–711.

24. Power, J. D., Cohen, A. L., Nelson, S. M., Wig, G. S., Barnes, K. A., Church, J. A., Vogel, A. C., Laumann, T. O., Miezin, F. M., Schlaggar, B. L., & Petersen, S. E. (2011). Functional network organization of the human brain. Neuron, 72(4), 665–678.

25. Ji, J. L., Spronk, M., Kulkarni, K., Repovš, G., Anticevic, A., & Cole, M. W. (2019). Mapping the human brain’s cortical-subcortical functional network organization. NeuroImage, 185, 35–57.

26. Saygin, Z. M., Osher, D. E., Koldewyn, K., Reynolds, G., Gabrieli, J. D. E., & Saxe, R. R. (2011). Anatomical connectivity patterns predict face selectivity in the fusiform gyrus. Nature Neuroscience, 15(2), 321–327

27. Cole, M. W., Ito, T., Bassett, D. S., & Schultz, D. H. (2016). Activity flow over resting-state networks shapes cognitive task activations. Nature Neuroscience, 19(12), 1718–1726.

28. Cocuzza, C. V., Sanchez-Romero, R., & Cole, M. W. (2022). Protocol for activity flow mapping of neurocognitive computations using the Brain Activity Flow Toolbox. STAR Protocols, 3(1), 101094.

29. 29. Pajani, A., Kouider, S., Roux, P., & de Gardelle, V. (2017). Unsuppressible Repetition Suppression and exemplar-specific Expectation Suppression in the Fusiform Face Area. Scientific Reports, 7(1), 160.

30. O’Reilly, R. C. (2001). Generalization in interactive networks: the benefits of inhibitory competition and Hebbian learning. Neural Computation, 13(6), 1199–1241.

31. Kriegeskorte, N., & Douglas, P. K. (2019). Interpreting encoding and decoding models. Current Opinion in Neurobiology, 55, 167–179.

32. Cole, M. W., Bassett, D. S., Power, J. D., Braver, T. S., & Petersen, S. E. (2014). Intrinsic and task-evoked network architectures of the human brain. Neuron, 83(1), 238–251.

33. NEW Krienen, F. M., Yeo, B. T. T., & Buckner, R. L. (2014). Reconfigurable task-dependent functional coupling modes cluster around a core functional architecture. *Philosophical Transactions of the Royal Society of London. Series B*, Biological Sciences, 369(1653).

34. NEW Gratton, C., Laumann, T. O., Nielsen, A. N., Greene, D. J., Gordon, E. M., Gilmore, A. W., Nelson, S. M., Coalson, R. S., Snyder, A. Z., Schlaggar, B. L., Dosenbach, N. U. F., & Petersen, S. E. (2018). Functional Brain Networks Are Dominated by Stable Group and Individual Factors, Not Cognitive or Daily Variation. Neuron, 98(2), 439–452.e5.

35. NEW Liégeois, R., Santos, A., Matta, V., Van De Ville, D., & Sayed, A. H. (2020). Revisiting correlation-based functional connectivity and its relationship with structural connectivity. *Network Neuroscience (Cambridge*, Mass.), 4(4), 1235–1251.

36. NEW Wodeyar, A., Cassidy, J. M., Cramer, S. C., & Srinivasan, R. (2020). Damage to the structural connectome reflected in resting-state fMRI functional connectivity. *Network Neuroscience (Cambridge*, Mass.), 4(4), 1197–1218.

37. NEW Peterson, K. L., Sanchez-Romero, R., Mill, R. D., & Cole, M. W. (2023). Regularized partial correlation provides reliable functional connectivity estimates while correcting for widespread confounding. In bioRxiv (p. 2023.09.16.558065).

38. NEW Sanchez-Romero, R., Ito, T., Mill, R. D., Hanson, S. J., & Cole, M. W. (2023). Causally informed activity flow models provide mechanistic insight into network-generated cognitive activations. NeuroImage, 278, 120300.

39. Van Essen, D. C., Smith, S. M., Barch, D. M., Behrens, T. E. J., Yacoub, E., Ugurbil, K., & WU-Minn HCP Consortium. (2013). The WU-Minn Human Connectome Project: an overview. NeuroImage, 80, 62–79.

40. Carandini, M., Demb, J. B., Mante, V., Tolhurst, D. J., Dan, Y., Olshausen, B. A., Gallant, J. L., & Rust, N. C. (2005). Do we know what the early visual system does? The Journal of Neuroscience: The Official Journal of the Society for Neuroscience, 25(46), 10577–10597.

41. Wandell, B. A., & Winawer, J. (2011). Imaging retinotopic maps in the human brain. Vision Research, 51(7), 718–737.

42. Glasser, M. F., Sotiropoulos, S. N., Wilson, J. A., Coalson, T. S., Fischl, B., Andersson, J. L., Xu, J., Jbabdi, S., Webster, M., Polimeni, J. R., Van Essen, D. C., Jenkinson, M., & WU-Minn HCP Consortium. (2013). The minimal preprocessing pipelines for the Human Connectome Project. NeuroImage, 80, 105–124.

43. Ito, T., Brincat, S. L., Siegel, M., Mill, R. D., He, B. J., Miller, E. K., Rotstein, H. G., & Cole, M. W. (2020a). Task-evoked activity quenches neural correlations and variability across cortical areas. PLoS Computational Biology, 16(8), e1007983.

44. Anderson, M. L., & Magruder, J. (2017). Split-Sample Strategies for Avoiding False Discoveries (No. 23544). National Bureau of Economic Research.

45. Barch, D. M., Burgess, G. C., Harms, M. P., Petersen, S. E., Schlaggar, B. L., Corbetta, M., Glasser, M. F., Curtiss, S., Dixit, S., Feldt, C., Nolan, D., Bryant, E., Hartley, T., Footer, O., Bjork, J. M., Poldrack, R., Smith, S., Johansen-Berg, H., Snyder, A. Z., … WU-Minn HCP Consortium. (2013). Function in the human connectome: task-fMRI and individual differences in behavior. NeuroImage, 80, 169–189.

46. Drobyshevsky, A., Baumann, S. B., & Schneider, W. (2006). A rapid fMRI task battery for mapping of visual, motor, cognitive, and emotional function. NeuroImage, 31(2), 732–744.

47. Wierenga, C. E., Perlstein, W. M., Benjamin, M., Leonard, C. M., Rothi, L. G., Conway, T., Cato, M. A., Gopinath, K., Briggs, R., & Crosson, B. (2009). Neural substrates of object identification: Functional magnetic resonance imaging evidence that category and visual attribute contribute to semantic knowledge. Journal of the International Neuropsychological Society: JINS, 15(2), 169– 181.

48. 48. Op de Beeck, H. P., Pillet, I., & Ritchie, J. B. (2019). Factors Determining Where Category-Selective Areas Emerge in Visual Cortex. Trends in Cognitive Sciences, 23(9), 784–797.

49. Kanwisher, N., McDermott, J., & Chun, M. M. (1997). The fusiform face area: a module in human extrastriate cortex specialized for face perception. The Journal of Neuroscience: The Official Journal of the Society for Neuroscience, 17(11), 4302–4311.

50. Iidaka, T. (2014). Role of the fusiform gyrus and superior temporal sulcus in face perception and recognition: An empirical review: Neuroimaging of face recognition. The Japanese Psychological Research, 56(1), 33–45.

51. Weiner, K. S., Barnett, M. A., Witthoft, N., Golarai, G., Stigliani, A., Kay, K. N., Gomez, J., Natu, V. S., Amunts, K., Zilles, K., & Grill-Spector, K. (2018). Defining the most probable location of the parahippocampal place area using cortex-based alignment and cross-validation. NeuroImage, 170, 373–384.

52. Grill-Spector, K., Kourtzi, Z., & Kanwisher, N. (2001). The lateral occipital complex and its role in object recognition. Vision Research, 41(10-11), 1409–1422.

53. Matić, K., Op de Beeck, H., & Bracci, S. (2020). It’s not all about looks: The role of object shape in parietal representations of manual tools. Cortex; a Journal Devoted to the Study of the Nervous System and Behavior.

54. Glasser, M. F., Coalson, T. S., Robinson, E. C., Hacker, C. D., Harwell, J., Yacoub, E., Ugurbil, K., Andersson, J., Beckmann, C. F., Jenkinson, M., Smith, S. M., & Van Essen, D. C. (2016). A multi-modal parcellation of human cerebral cortex. Nature, 536(7615), 171–178.

55. Ciric, R., Wolf, D. H., Power, J. D., Roalf, D. R., Baum, G. L., Ruparel, K., Shinohara, R. T., Elliott, M. A., Eickhoff, S. B., Davatzikos, C., Gur, R. C., Gur, R. E., Bassett, D. S., & Satterthwaite, T. D. (2017). Benchmarking of participant-level confound regression strategies for the control of motion artifact in studies of functional connectivity. NeuroImage, 154, 174–187.

56. Behzadi, Y., Restom, K., Liau, J., & Liu, T. T. (2007). A component based noise correction method (CompCor) for BOLD and perfusion based fMRI. NeuroImage, 37(1), 90–101.

57. Murphy, K., Birn, R. M., Handwerker, D. A., Jones, T. B., & Bandettini, P. A. (2009). The impact of global signal regression on resting state correlations: are anti-correlated networks introduced? NeuroImage, 44(3), 893–905.

58. Power, J. D., Mitra, A., Laumann, T. O., Snyder, A. Z., Schlaggar, B. L., & Petersen, S. E. (2014). Methods to detect, characterize, and remove motion artifact in resting state fMRI. NeuroImage, 84, 320–341.

59. Power, J. D., Plitt, M., Gotts, S. J., Kundu, P., Voon, V., Bandettini, P. A., & Martin, A. (2018). Ridding fMRI data of motion-related influences: Removal of signals with distinct spatial and physical bases in multiecho data. Proceedings of the National Academy of Sciences of the United States of America, 115(9), E2105–E2114.

60. Friston, K. J., Holmes, A. P., Worsley, K. J., Poline, J. P., Frith, C. D., & Frackowiak, R. S. J. (1994). Statistical parametric maps in functional imaging: A general linear approach. Human Brain Mapping, 2(4), 189–210.

61. Poldrack, R. A., Huckins, G., & Varoquaux, G. (2019). Establishment of Best Practices for Evidence for Prediction: A Review. JAMA Psychiatry.

62. Malikovic, A., Amunts, K., Schleicher, A., Mohlberg, H., Eickhoff, S. B., Wilms, M., Palomero-Gallagher, N., Armstrong, E., & Zilles, K. (2007). Cytoarchitectonic analysis of the human extrastriate cortex in the region of V5/MT+: a probabilistic, stereotaxic map of area hOc5. Cerebral Cortex , 17(3), 562–574.

63. Fischl, B., Rajendran, N., Busa, E., Augustinack, J., Hinds, O., Yeo, B. T. T., Mohlberg, H., Amunts, K., & Zilles, K. (2008). Cortical folding patterns and predicting cytoarchitecture. Cerebral Cortex , 18(8), 1973–1980.

64. Kolster, H., Peeters, R., & Orban, G. A. (2010). The retinotopic organization of the human middle temporal area MT/V5 and its cortical neighbors. The Journal of Neuroscience: The Official Journal of the Society for Neuroscience, 30(29), 9801–9820.

65. Abdollahi, R. O., Kolster, H., Glasser, M. F., Robinson, E. C., Coalson, T. S., Dierker, D., Jenkinson, M., Van Essen, D. C., & Orban, G. A. (2014). Correspondences between retinotopic areas and myelin maps in human visual cortex. NeuroImage, 99, 509–524.

66. Peelen, M. V., & Downing, P. E. (2005). Selectivity for the human body in the fusiform gyrus. Journal of Neurophysiology, 93(1), 603–608.

67. Spiridon, M., Fischl, B., & Kanwisher, N. (2006). Location and spatial profile of category-specific regions in human extrastriate cortex. Human Brain Mapping, 27(1), 77–89.

68. Vocks, S., Busch, M., Grönemeyer, D., Schulte, D., Herpertz, S., & Suchan, B. (2010). Differential neuronal responses to the self and others in the extrastriate body area and the fusiform body area. Cognitive, Affective & Behavioral Neuroscience, 10(3), 422–429.

69. Shultz, S., Lee, S. M., Pelphrey, K., & McCarthy, G. (2011). The posterior superior temporal sulcus is sensitive to the outcome of human and non-human goal-directed actions. Social Cognitive and Affective Neuroscience, 6(5), 602–611.

70. Ross, P. D., de Gelder, B., Crabbe, F., & Grosbras, M.-H. (2014). Body-selective areas in the visual cortex are less active in children than in adults. Frontiers in Human Neuroscience, 8, 941.

71. Orgs, G., Dovern, A., Hagura, N., Haggard, P., Fink, G. R., & Weiss, P. H. (2016). Constructing Visual Perception of Body Movement with the Motor Cortex. Cerebral Cortex , 26(1), 440–449.

72. Zimmermann, M., Mars, R. B., de Lange, F. P., Toni, I., & Verhagen, L. (2018). Is the extrastriate body area part of the dorsal visuomotor stream? Brain Structure & Function, 223(1), 31–46.

73. von Economo, C. F., & Koskinas, G. N. (1925). Die cytoarchitektonik der hirnrinde des erwachsenen menschen. J. Springer.

74. Triarhou, L. C. (2007). The Economo-Koskinas atlas revisited: cytoarchitectonics and functional context. Stereotactic and Functional Neurosurgery, 85(5), 195–203.

75. Larsson, J., & Heeger, D. J. (2006). Two retinotopic visual areas in human lateral occipital cortex. The Journal of Neuroscience: The Official Journal of the Society for Neuroscience, 26(51), 13128– 13142.

76. Hodzic, A., Kaas, A., Muckli, L., Stirn, A., & Singer, W. (2009). Distinct cortical networks for the detection and identification of human body. NeuroImage, 45(4), 1264–1271.

77. Taylor, J. C., Wiggett, A. J., & Downing, P. E. (2010). fMRI-adaptation studies of viewpoint tuning in the extrastriate and fusiform body areas. Journal of Neurophysiology, 103(3), 1467–1477.

78. Ewbank, M. P., Lawson, R. P., Henson, R. N., Rowe, J. B., Passamonti, L., & Calder, A. J. (2011). Changes in “top-down” connectivity underlie repetition suppression in the ventral visual pathway. The Journal of Neuroscience: The Official Journal of the Society for Neuroscience, 31(15), 5635– 5642.

79. Hashimoto, T., & Iriki, A. (2013). Dissociations between the horizontal and dorsoventral axes in body-size perception. The European Journal of Neuroscience, 37(11), 1747–1753.

80. Kim, N. Y., Lee, S. M., Erlendsdottir, M. C., & McCarthy, G. (2014). Discriminable spatial patterns of activation for faces and bodies in the fusiform gyrus. Frontiers in Human Neuroscience, 8, 632.

81. Ross, P., de Gelder, B., Crabbe, F., & Grosbras, M.-H. (2020). A dynamic body-selective area localizer for use in fMRI. MethodsX, 7, 100801.

82. Glasser, M. F., & Van Essen, D. C. (2011). Mapping human cortical areas in vivo based on myelin content as revealed by T1- and T2-weighted MRI. The Journal of Neuroscience: The Official Journal of the Society for Neuroscience, 31(32), 11597–11616.

83. Caspers, J., Zilles, K., Eickhoff, S. B., Schleicher, A., Mohlberg, H., & Amunts, K. (2013). Cytoarchitectonical analysis and probabilistic mapping of two extrastriate areas of the human posterior fusiform gyrus. Brain Structure & Function, 218(2), 511–526.

84. Weiner, K. S., Golarai, G., Caspers, J., Chuapoco, M. R., Mohlberg, H., Zilles, K., Amunts, K., & Grill-Spector, K. (2014). The mid-fusiform sulcus: a landmark identifying both cytoarchitectonic and functional divisions of human ventral temporal cortex. NeuroImage, 84, 453–465.

85. Turk-Browne, N. B., Norman-Haignere, S. V., & McCarthy, G. (2010). Face-Specific Resting Functional Connectivity between the Fusiform Gyrus and Posterior Superior Temporal Sulcus. Frontiers in Human Neuroscience, 4, 176.

86. Nagy, K., Greenlee, M. W., & Kovács, G. (2012). The lateral occipital cortex in the face perception network: an effective connectivity study. Frontiers in Psychology, 3, 141.

87. Baseler, H. A., Harris, R. J., Young, A. W., & Andrews, T. J. (2014). Neural responses to expression and gaze in the posterior superior temporal sulcus interact with facial identity. Cerebral Cortex , 24(3), 737–744.

88. Flack, T. R., Andrews, T. J., Hymers, M., Al-Mosaiwi, M., Marsden, S. P., Strachan, J. W. A., Trakulpipat, C., Wang, L., Wu, T., & Young, A. W. (2015). Responses in the right posterior superior temporal sulcus show a feature-based response to facial expression. Cortex; a Journal Devoted to the Study of the Nervous System and Behavior, 69, 14–23.

89. Axelrod, V., & Yovel, G. (2015). Successful decoding of famous faces in the fusiform face area. PloS One, 10(2), e0117126.

90. Schobert, A.-K., Corradi-Dell’Acqua, C., Frühholz, S., van der Zwaag, W., & Vuilleumier, P. (2018). Functional organization of face processing in the human superior temporal sulcus: a 7T high-resolution fMRI study. Social Cognitive and Affective Neuroscience, 13(1), 102–113.

91. Li, J., Song, Y., & Liu, J. (2019). Functional connectivity pattern in the core face network reflects different mechanisms of holistic face processing measured by the whole-part effect and composite-face effect. Neuroscience, 408, 248–258.

92. Pitcher, D., Pilkington, A., Rauth, L., Baker, C., Kravitz, D. J., & Ungerleider, L. G. (2020). The Human Posterior Superior Temporal Sulcus Samples Visual Space Differently From Other Face-Selective Regions. Cerebral Cortex, 30(2), 778–785.

93. Engell, A. D., & Haxby, J. V. (2007). Facial expression and gaze-direction in human superior temporal sulcus. Neuropsychologia, 45(14), 3234–3241.

94. von dem Hagen, E. A. H., Nummenmaa, L., Yu, R., Engell, A. D., Ewbank, M. P., & Calder, A. J. (2011). Autism spectrum traits in the typical population predict structure and function in the posterior superior temporal sulcus. Cerebral Cortex, 21(3), 493–500.

95. Watson, R., Latinus, M., Noguchi, T., Garrod, O., Crabbe, F., & Belin, P. (2014). Crossmodal adaptation in right posterior superior temporal sulcus during face-voice emotional integration. The Journal of Neuroscience: The Official Journal of the Society for Neuroscience, 34(20), 6813–6821.

96. Thurman, S. M., van Boxtel, J. J. A., Monti, M. M., Chiang, J. N., & Lu, H. (2016). Neural adaptation in pSTS correlates with perceptual aftereffects to biological motion and with autistic traits. NeuroImage, 136, 149–161.

97. Epstein, R. A., & Higgins, J. S. (2007). Differential parahippocampal and retrosplenial involvement in three types of visual scene recognition. Cerebral Cortex , 17(7), 1680–1693.

98. Henderson, J. M., Larson, C. L., & Zhu, D. C. (2008). Full scenes produce more activation than close-up scenes and scene-diagnostic objects in parahippocampal and retrosplenial cortex: an fMRI study. Brain and Cognition, 66(1), 40–49.

99. Pourtois, G., Schwartz, S., Spiridon, M., Martuzzi, R., & Vuilleumier, P. (2009). Object representations for multiple visual categories overlap in lateral occipital and medial fusiform cortex. Cerebral Cortex, 19(8), 1806–1819.

100. Bastin, J., Vidal, J. R., Bouvier, S., Perrone-Bertolotti, M., Bénis, D., Kahane, P., David, O., Lachaux, J.-P., & Epstein, R. A. (2013). Temporal components in the parahippocampal place area revealed by human intracerebral recordings. The Journal of Neuroscience: The Official Journal of the Society for Neuroscience, 33(24), 10123–10131.

101. Sulpizio, V., Committeri, G., & Galati, G. (2014). Distributed cognitive maps reflecting real distances between places and views in the human brain. Frontiers in Human Neuroscience, 8, 716.

102. Marchette, S. A., Vass, L. K., Ryan, J., & Epstein, R. A. (2015). Outside Looking In: Landmark Generalization in the Human Navigational System. The Journal of Neuroscience: The Official Journal of the Society for Neuroscience, 35(44), 14896–14908.

103. Bonner, M. F., Price, A. R., Peelle, J. E., & Grossman, M. (2016). Semantics of the Visual Environment Encoded in Parahippocampal Cortex. Journal of Cognitive Neuroscience, 28(3), 361– 378.

104. Bellana, B., Liu, Z.-X., Diamond, N. B., Grady, C. L., & Moscovitch, M. (2017). Similarities and differences in the default mode network across rest, retrieval, and future imagining. Human Brain Mapping, 38(3), 1155–1171.

105. Wang, L., Mruczek, R. E. B., Arcaro, M. J., & Kastner, S. (2015). Probabilistic Maps of Visual Topography in Human Cortex. Cerebral Cortex , 25(10), 3911–3931.

106. Arcaro, M. J., McMains, S. A., Singer, B. D., & Kastner, S. (2009). Retinotopic organization of human ventral visual cortex. The Journal of Neuroscience: The Official Journal of the Society for Neuroscience, 29(34), 10638–10652.

107. Wolbers, T., & Büchel, C. (2005). Dissociable retrosplenial and hippocampal contributions to successful formation of survey representations. The Journal of Neuroscience: The Official Journal of the Society for Neuroscience, 25(13), 3333–3340.

108. Sherrill, K. R., Erdem, U. M., Ross, R. S., Brown, T. I., Hasselmo, M. E., & Stern, C. E. (2013). Hippocampus and retrosplenial cortex combine path integration signals for successful navigation. The Journal of Neuroscience: The Official Journal of the Society for Neuroscience, 33(49), 19304– 19313.

109. Burles, F., Slone, E., & Iaria, G. (2017). Dorso-medial and ventro-lateral functional specialization of the human retrosplenial complex in spatial updating and orienting. Brain Structure & Function, 222(3), 1481–1493.

110. Rottschy, C., Eickhoff, S. B., & Schleicher, A. (2007). Ventral visual cortex in humans: cytoarchitectonic mapping of two extrastriate areas. Human Brain Mapping.

111. Hansen, K. A., Kay, K. N., & Gallant, J. L. (2007). Topographic organization in and near human visual area V4. The Journal of Neuroscience: The Official Journal of the Society for Neuroscience, 27(44), 11896–11911.

112. Malikovic, A., Amunts, K., Schleicher, A., Mohlberg, H., Kujovic, M., Palomero-Gallagher, N., Eickhoff, S. B., & Zilles, K. (2016). Cytoarchitecture of the human lateral occipital cortex: mapping of two extrastriate areas hOc4la and hOc4lp. Brain Structure & Function, 221(4), 1877–1897.

113. Grill-Spector, K., Kushnir, T., Hendler, T., Edelman, S., Itzchak, Y., & Malach, R. (1998). A sequence of object-processing stages revealed by fMRI in the human occipital lobe. Human Brain Mapping, 6(4), 316–328.

114. Niemeier, M., Goltz, H. C., Kuchinad, A., Tweed, D. B., & Vilis, T. (2005). A contralateral preference in the lateral occipital area: sensory and attentional mechanisms. Cerebral Cortex , 15(3), 325– 331.

115. Silvanto, J., Schwarzkopf, D. S., Gilaie-Dotan, S., & Rees, G. (2010). Differing causal roles for lateral occipital cortex and occipital face area in invariant shape recognition. The European Journal of Neuroscience, 32(1), 165–171.

116. Cant, J. S., & Goodale, M. A. (2011). Scratching beneath the surface: new insights into the functional properties of the lateral occipital area and parahippocampal place area. The Journal of Neuroscience: The Official Journal of the Society for Neuroscience, 31(22), 8248–8258.

117. McFadden, K. L., Hepburn, S., Winterrowd, E., Schmidt, G. L., & Rojas, D. C. (2012). Abnormalities in gamma-band responses to language stimuli in first-degree relatives of children with autism spectrum disorder: an MEG study. BMC Psychiatry, 12, 213.

118. Karanian, J. M., & Slotnick, S. D. (2015). Memory for shape reactivates the lateral occipital complex. Brain Research, 1603, 124–132.

119. Chiou, R., & Lambon Ralph, M. A. (2016). Task-Related Dynamic Division of Labor Between Anterior Temporal and Lateral Occipital Cortices in Representing Object Size. The Journal of Neuroscience: The Official Journal of the Society for Neuroscience, 36(17), 4662–4668.

120. Erdogan, G., Chen, Q., Garcea, F. E., Mahon, B. Z., & Jacobs, R. A. (2016). Multisensory Part-based Representations of Objects in Human Lateral Occipital Cortex. Journal of Cognitive Neuroscience, 28(6), 869–881.

121. Chouinard, P. A., Meena, D. K., Whitwell, R. L., Hilchey, M. D., & Goodale, M. A. (2017). A TMS Investigation on the Role of Lateral Occipital Complex and Caudal Intraparietal Sulcus in the Perception of Object Form and Orientation. Journal of Cognitive Neuroscience, 29(5), 881–895.

122. Bahram, M., Münte, T. F., Cole, D. M., Amir, S., Melanie, B., & Jascha, R. (2020). Changed functional connectivity at rest in functional illiterates after extensive literacy training. Berlin, 2(1), s42466–020.

123. Hadjikhani, N., Liu, A. K., Dale, A. M., Cavanagh, P., & Tootell, R. B. (1998). Retinotopy and color sensitivity in human visual cortical area V8. Nature Neuroscience, 1(3), 235–241.

124. Lacadie, C. M., Fulbright, R. K., Rajeevan, N., Constable, R. T., & Papademetris, X. (2008). More accurate Talairach coordinates for neuroimaging using non-linear registration. NeuroImage, 42(2), 717–725.

125. Sanchez-Romero, R., & Cole, M. W. (2021). Combining Multiple Functional Connectivity Methods to Improve Causal Inferences. Journal of Cognitive Neuroscience, 33(2), 180–194.

126. Spirtes, P., Glymour, C. N., Scheines, R., & Heckerman, D. (2000). Causation, Prediction, and Search. MIT Press.

127. Norman-Haignere, S. V., McCarthy, G., Chun, M. M., & Turk-Browne, N. B. (2012). Category-selective background connectivity in ventral visual cortex. Cerebral Cortex , 22(2), 391–402.

128. Hermundstad, A. M., Bassett, D. S., Brown, K. S., Aminoff, E. M., Clewett, D., Freeman, S., Frithsen, A., Johnson, A., Tipper, C. M., Miller, M. B., Grafton, S. T., & Carlson, J. M. (2013). Structural foundations of resting-state and task-based functional connectivity in the human brain. Proceedings of the National Academy of Sciences of the United States of America, 110(15), 6169– 6174.

129. Betzel, R. F., & Bassett, D. S. (2017). Generative models for network neuroscience: prospects and promise. *Journal of the Royal Society*, Interface / the Royal Society, 14(136).

130. Leys, C., Ley, C., Klein, O., Bernard, P., & Licata, L. (2013). Detecting outliers: Do not use standard deviation around the mean, use absolute deviation around the median. Journal of Experimental Social Psychology, 49(4), 764–766.

131. Azen, R., & Budescu, D. V. (2003). The dominance analysis approach for comparing predictors in multiple regression. Psychological Methods, 8(2), 129–148.

132. Kraha, A., Turner, H., Nimon, K., Zientek, L. R., & Henson, R. K. (2012). Tools to support interpreting multiple regression in the face of multicollinearity. Frontiers in Psychology, 3, 44.

133. Wiedermann, M., Donges, J. F., Kurths, J., & Donner, R. V. (2016). Spatial network surrogates for disentangling complex system structure from spatial embedding of nodes. Physical Review. E, 93, 042308.

134. Markello, R. D., & Misic, B. (2021). Comparing spatial null models for brain maps. NeuroImage, 236, 118052.

135. Zimmermann, M., Mars, R. B., de Lange, F. P., Toni, I., & Verhagen, L. (2018). Is the extrastriate body area part of the dorsal visuomotor stream? Brain Structure & Function, 223(1), 31–46.

136. Wang, X., Zhen, Z., Song, Y., Huang, L., Kong, X., & Liu, J. (2016). The Hierarchical Structure of the Face Network Revealed by Its Functional Connectivity Pattern. The Journal of Neuroscience: The Official Journal of the Society for Neuroscience, 36(3), 890–900.

137. Liu, J., Harris, A., & Kanwisher, N. (2010). Perception of face parts and face configurations: an FMRI study. Journal of Cognitive Neuroscience, 22(1), 203–211.

138. Zhen, Z., Fang, H., & Liu, J. (2013). The hierarchical brain network for face recognition. PloS One, 8(3), e59886.

139. Pitcher, D., Walsh, V., & Duchaine, B. (2011). The role of the occipital face area in the cortical face perception network. Experimental Brain Research. Experimentelle Hirnforschung. Experimentation Cerebrale, 209(4), 481–493.

140. Engell, A. D., & McCarthy, G. (2013). Probabilistic atlases for face and biological motion perception: an analysis of their reliability and overlap. NeuroImage, 74, 140–151.

141. Ishai, A. (2008). Let’s face it: it’s a cortical network. NeuroImage, 40(2), 415–419.

142. Troiani, V., Dougherty, C. C., Michael, A. M., & Olson, I. R. (2016). Characterization of Face-Selective Patches in Orbitofrontal Cortex. Frontiers in Human Neuroscience, 10, 279.

143. Barat, E., Wirth, S., & Duhamel, J.-R. (2018). Face cells in orbitofrontal cortex represent social categories. Proceedings of the National Academy of Sciences of the United States of America, 115(47), E11158–E11167.

144. Bettencourt, K. C., & Xu, Y. (2013). The role of transverse occipital sulcus in scene perception and its relationship to object individuation in inferior intraparietal sulcus. Journal of Cognitive Neuroscience, 25(10), 1711–1722.

145. Welbourne, L. E., Jonnalagadda, A., Giesbrecht, B., & Eckstein, M. P. (2021). The transverse occipital sulcus and intraparietal sulcus show neural selectivity to object-scene size relationships. Communications Biology, 4(1), 768.

146. Tobia, M. J., & Madan, C. R. (2017). Tool selection and the ventral-dorsal organization of tool-related knowledge. Physiological Reports, 5(3).

147. Grabowski, T. J., Damasio, H., & Damasio, A. R. (1998). Premotor and prefrontal correlates of category-related lexical retrieval. NeuroImage, 7(3), 232–243.

148. Chen, J., Snow, J. C., Culham, J. C., & Goodale, M. A. (2018). What Role Does “Elongation” Play in “Tool-Specific” Activation and Connectivity in the Dorsal and Ventral Visual Streams? Cerebral Cortex , 28(4), 1117–1131.

149. Mruczek, R. E. B., von Loga, I. S., & Kastner, S. (2013). The representation of tool and non-tool object information in the human intraparietal sulcus. Journal of Neurophysiology, 109(12), 2883– 2896.

150. Newman, M. E. J., & Girvan, M. (2004). Finding and evaluating community structure in networks. *Physical Review. E, Statistical*, Nonlinear, and Soft Matter Physics, 69(2 Pt 2), 026113.

151. Fosdick, B. K., Larremore, D. B., Nishimura, J., & Ugander, J. (2018). Configuring Random Graph Models with Fixed Degree Sequences. SIAM Review, 60(2), 315–355.

152. Nichols, T. E., & Holmes, A. P. (2002). Nonparametric permutation tests for functional neuroimaging: a primer with examples. Human Brain Mapping, 15(1), 1–25.

153. Carey, M., Knight, R., & Preston, C. (2019). Distinct neural response to visual perspective and body size in the extrastriate body area. Behavioural Brain Research, 372, 112063.

154. Bratch, A., Engel, S., Burton, P., & Kersten, D. (2018). The Fusiform Body Area Represents Spatial Relationships Between Pairs of Body Parts. Journal of Vision, 18(10), 408–408.

155. Puce, A., Allison, T., Bentin, S., Gore, J. C., & McCarthy, G. (1998). Temporal cortex activation in humans viewing eye and mouth movements. The Journal of Neuroscience: The Official Journal of the Society for Neuroscience, 18(6), 2188–2199.

156. Epstein, R., Graham, K. S., & Downing, P. E. (2003). Viewpoint-specific scene representations in human parahippocampal cortex. Neuron, 37(5), 865–876.

157. Aminoff, E. M., Kveraga, K., & Bar, M. (2013). The role of the parahippocampal cortex in cognition. Trends in Cognitive Sciences, 17(8), 379–390.

158. Kravitz, D. J., Saleem, K. S., Baker, C. I., & Mishkin, M. (2011). A new neural framework for visuospatial processing. Nature Reviews. Neuroscience, 12(4), 217–230.

159. Vann, S. D., Aggleton, J. P., & Maguire, E. A. (2009). What does the retrosplenial cortex do? Nature Reviews. Neuroscience, 10(11), 792–804.

160. Bar, M. (2004). Visual objects in context. Nature Reviews. Neuroscience, 5(8), 617–629.

161. Beauchamp, M. S., Lee, K. E., Haxby, J. V., & Martin, A. (2002). Parallel visual motion processing streams for manipulable objects and human movements. Neuron.

162. Kourtzi, Z., & Kanwisher, N. (2001). Representation of perceived object shape by the human lateral occipital complex. Science, 293(5534), 1506–1509.

163. Victoria, L. W., Pyles, J. A., & Tarr, M. J. (2019). The relative contributions of visual and semantic information in the neural representation of object categories. Brain and Behavior, 9(10), e01373.

164. St-Yves, G., Allen, E. J., Wu, Y., Kay, K., & Naselaris, T. (2022). Brain-optimized neural networks learn non-hierarchical models of representation in human visual cortex. In bioRxiv (p. 2022.01.21.477293).

165. Sexton, N. J., & Love, B. C. (2022). Reassessing hierarchical correspondences between brain and deep networks through direct interface. Science Advances, 8(28), eabm2219.

166. Wojciulik, E., Kanwisher, N., & Driver, J. (1998). Covert visual attention modulates face-specific activity in the human fusiform gyrus: fMRI study. Journal of Neurophysiology, 79(3), 1574–1578.

167. Saalmann, Y. B., Pigarev, I. N., & Vidyasagar, T. R. (2007). Neural mechanisms of visual attention: how top-down feedback highlights relevant locations. Science, 316(5831), 1612–1615.

168. Kay, K. N., & Yeatman, J. D. (2017). Bottom-up and top-down computations in word- and face-selective cortex. eLife, 6.

169. 169. Summerfield, C., & de Lange, F. P. (2014). Expectation in perceptual decision making: neural and computational mechanisms. Nature Reviews. Neuroscience, 15(11), 745–756.

170. Vaziri-Pashkam, M., & Xu, Y. (2017). Goal-Directed Visual Processing Differentially Impacts Human Ventral and Dorsal Visual Representations. The Journal of Neuroscience: The Official Journal of the Society for Neuroscience, 37(36), 8767–8782.

171. Bracci, S., Daniels, N., & Op de Beeck, H. (2017). Task Context Overrules Object- and Category-Related Representational Content in the Human Parietal Cortex. Cerebral Cortex , 27(1), 310–321.

172. De Cesarei, A., Cavicchi, S., Micucci, A., & Codispoti, M. (2019). Categorization Goals Modulate the Use of Natural Scene Statistics. Journal of Cognitive Neuroscience, 31(1), 109–125.

173. von Helmholtz, H. (1867). Handbuch der physiologischen Optik: mit 213 in den Text eingedruckten Holzschnitten und 11 Tafeln. Voss.

174. Yuille, A., & Kersten, D. (2006). Vision as Bayesian inference: analysis by synthesis? Trends in Cognitive Sciences, 10(7), 301–308.

175. Fang, M., Aglinskas, A., Li, Y., & Anzellotti, S. (2019). Identifying hubs that integrate responses across multiple category-selective regions.

176. Vuilleumier, P., & Driver, J. (2007). Modulation of visual processing by attention and emotion: windows on causal interactions between human brain regions. *Philosophical Transactions of the Royal Society of London. Series B*, Biological Sciences, 362(1481), 837–855.

177. Bryan, P. B., Julian, J. B., & Epstein, R. A. (2016). Rectilinear Edge Selectivity Is Insufficient to Explain the Category Selectivity of the Parahippocampal Place Area. Frontiers in Human Neuroscience, 10, 137.

178. Coggan, D. D., Allen, L. A., Farrar, O. R. H., Gouws, A. D., Morland, A. B., Baker, D. H., & Andrews, T. J. (2017). Differences in selectivity to natural images in early visual areas (V1–V3). Scientific Reports, 7(1), 1–8.

179. Poltoratski, S., Kay, K., Finzi, D., & Grill-Spector, K. (2021). Holistic face recognition is an emergent phenomenon of spatial processing in face-selective regions. Nature Communications, 12(1), 4745.

180. Grill-Spector, K., Weiner, K. S., Kay, K., & Gomez, J. (2017). The Functional Neuroanatomy of Human Face Perception. Annual Review of Vision Science, 3, 167–196.

181. Kietzmann, T. C., Spoerer, C. J., Sörensen, L. K. A., Cichy, R. M., Hauk, O., & Kriegeskorte, N. (2019). Recurrence is required to capture the representational dynamics of the human visual system. Proceedings of the National Academy of Sciences of the United States of America, 116(43), 21854–21863.

182. Keane, B. P., Barch, D. M., Mill, R. D., Silverstein, S. M., Krekelberg, B., & Cole, M. W. (2021). Brain network mechanisms of visual shape completion. NeuroImage, 236, 118069.

183. Logothetis, N. K. (2003). The underpinnings of the BOLD functional magnetic resonance imaging signal. The Journal of Neuroscience: The Official Journal of the Society for Neuroscience, 23(10), 3963–3971.

184. Kanwisher, N. (2017). The Quest for the FFA and Where It Led. The Journal of Neuroscience: The Official Journal of the Society for Neuroscience, 37(5), 1056–1061.

185. Kong, R., Li, J., Orban, C., Sabuncu, M. R., Liu, H., Schaefer, A., Sun, N., Zuo, X.-N., Holmes, A. J., Eickhoff, S. B., & Yeo, B. T. T. (2019). Spatial Topography of Individual-Specific Cortical Networks Predicts Human Cognition, Personality, and Emotion. Cerebral Cortex , 29(6), 2533–2551.

