## Supplementary material for "Distributed network flows generate localized category selectivity in human visual cortex": S1 Fig

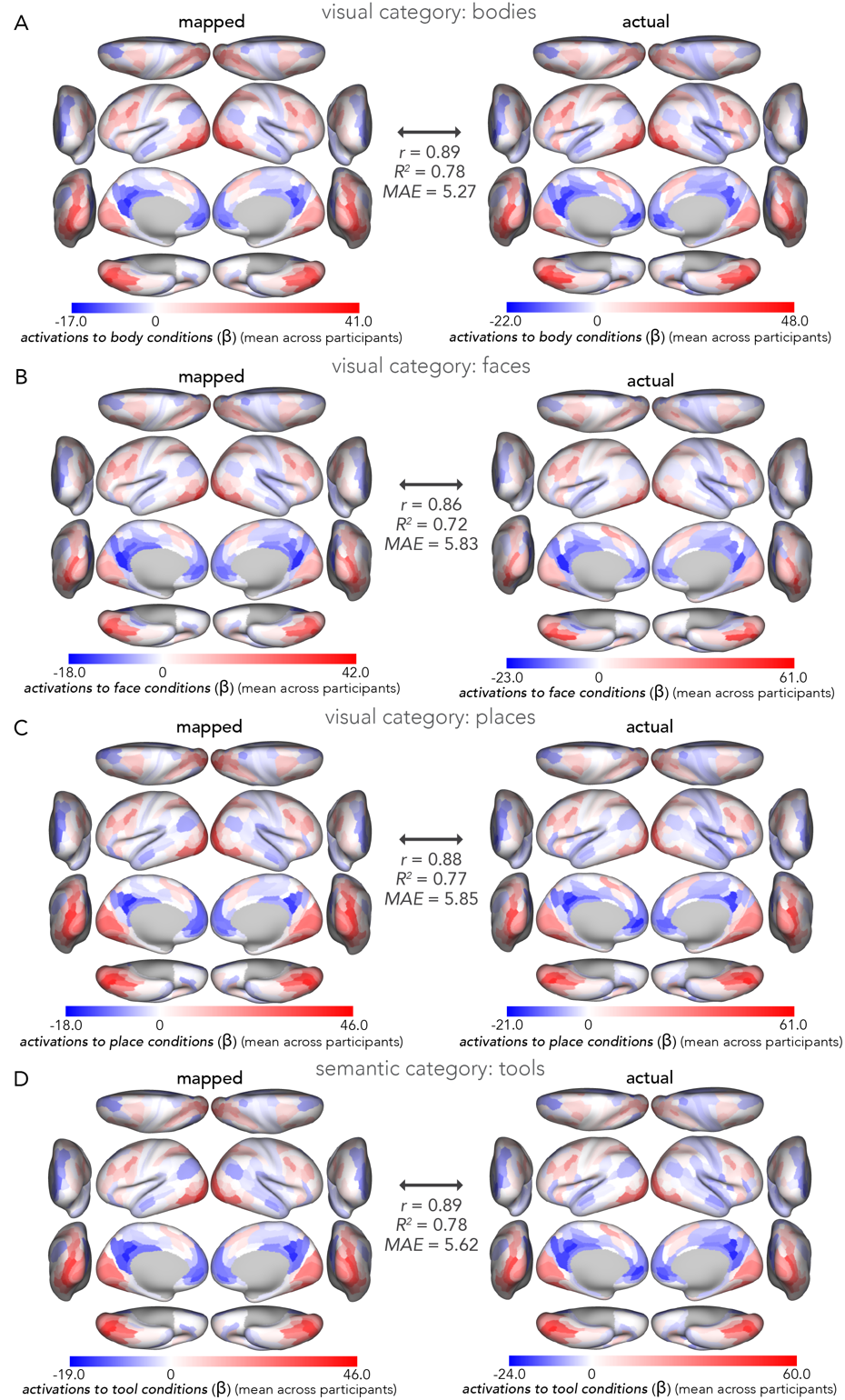

**S1 Fig. Whole-cortex activity-flow-mapped activations for four semantic visual categories.** This assessment demonstrates that activity flow mapping of cross-cortex activations to each visual semantic category exhibits high accuracy. (**A**) Left: Cross-participant average actual (empirical) task activations to body categories projected onto the MMP cortical atlas [54]. Right: Cross-participant average activity-flow-mapped task activations to body categories projected onto the MMP cortical atlas. The mapped and actual activations exhibited a high degree of overlap: *r* = 0.89. (**B-D**) The same as in A, but for face (B), place (C), and tool categories (D) respectively. In all cases, accuracy was high, demonstrating that activity flow processes mapped predicted cross-cortex responses to each visual category of interest well.
