## Supplementary material for "Distributed network flows generate localized category selectivity in human visual cortex": S1 Table

#### **S1 Table. Whole-cortex activity-flow-mapped responses to visual category conditions.**

| Analysis | Dataset | *r* | MAE | R^2^ |
| --- | --- | --- | --- | --- |
| Response profiles of four functional complexes: |  |  |  |  |
| Mapping accuracy: right hemisphere | Replication | 0.92 | 4.09 | 0.79 |
| Mapping accuracy: left hemisphere | Replication | 0.92 | 4.01 | 0.79 |
| Category-specific responses across the whole cortex: |  |  |  |  |
| Mapping accuracy: body image categories | Replication | 0.89 | 5.29 | 0.77 |
| Mapping accuracy: face image categories | Replication | 0.85 | 5.29 | 0.71 |
| Mapping accuracy: place image categories | Replication | 0.88 | 6.11 | 0.76 |
| Mapping accuracy: tools image categories | Replication | 0.89 | 5.68 | 0.78 |

Across select portions of cortex (whole cortex = all 360 MMP cortical regions [54]) and all *n*=176 participants, the accuracy of activity flow mapping was estimated by comparing of mapped and actual responses to select HCP conditions (response profile = across all 24 HCP conditions) via Pearson’s *r*, mean absolute error (MAE), and the coefficient of determination (R^2^).
