## Supplementary material for "Distributed network flows generate localized category selectivity in human visual cortex": S2 Fig

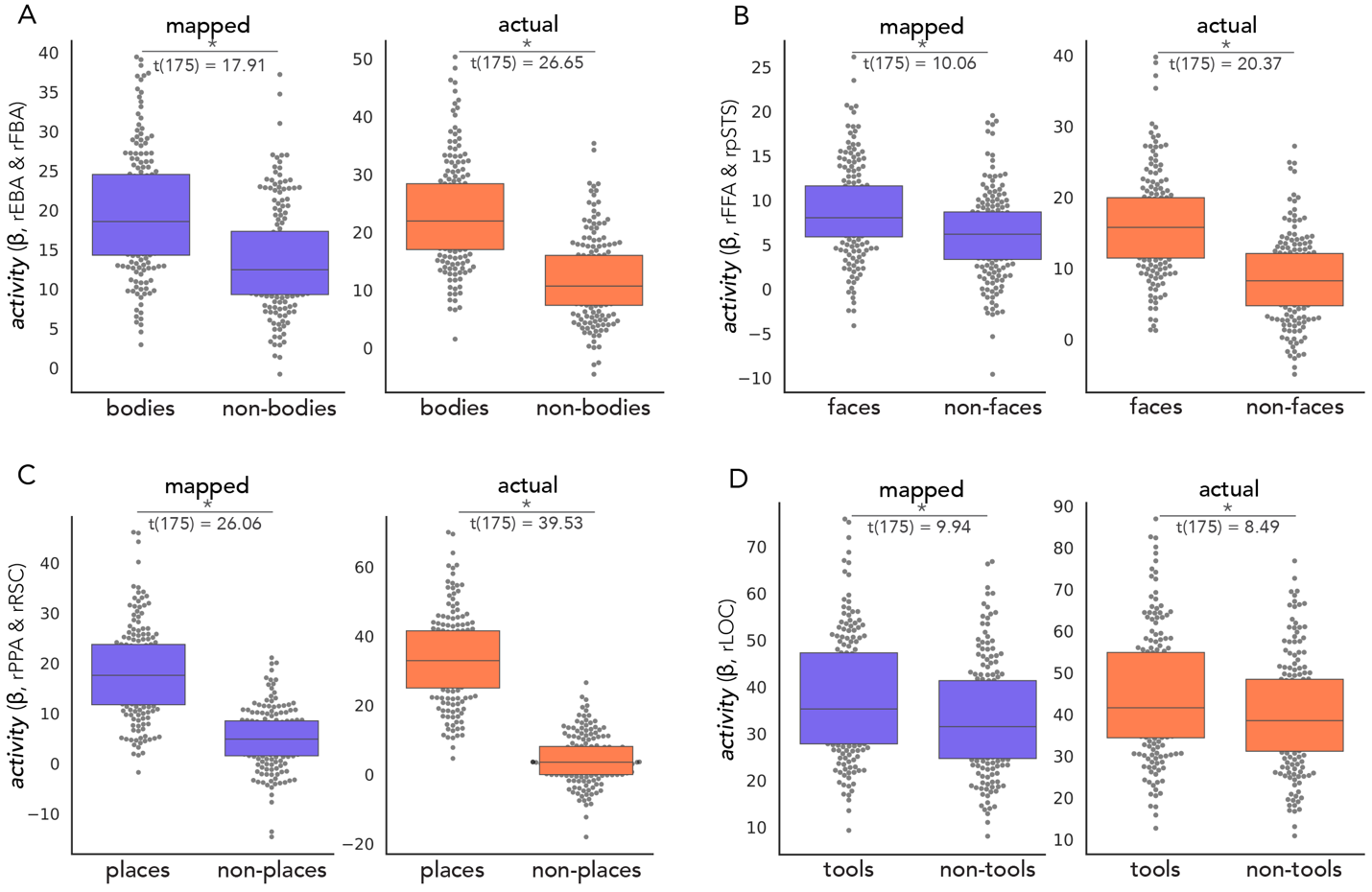

**S2 Fig. Benchmarking activity in four functional complexes.**To corroborate findings in the literature that each of the four functional complexes assessed in the present study exhibit significantly higher activations to images in their respective visual categories, we conducted standard t-test contrasts (discovery dataset, right hemisphere results depicted here; left hemisphere statistics reported in S1 Table). (**A**) Box and swarm plots depicting activations to images of bodies and body parts exhibited by the EBA/FBA (dots = individual participant’s data). The actual activity is shown in coral and the activity-flow-mapped activity is shown in purple. Body versus non-body activations were contrasted with a one-tailed, paired samples t-test (hypothesizing that body activity was larger than non-body activity in the EBA/FBA), with an asterisk indicating a statistically significant difference (*p* < 0.0001 with nonparametric permutation tests, as reported in the main text Results). The EBA/FBA exhibited statistically greater activations to body versus non-body images, benchmarking the prior work in the literature. (**B**) Same as in A, but for face versus non-face images and the FFA/pSTS functional complex. (**C**) Same as in A, but for place versus non-place images and the PPA/RSC functional complex. (**D**) Same as in A, but for tool versus non-tool images and the LOC functional complex. All results were corroborated by the replication dataset (*N*=176 in each; statistics given in main Results text).
