## Supplementary material for "Distributed network flows generate localized category selectivity in human visual cortex": S2 Table

#### **S2 Table. Benchmark contrasts in four functional complexes.**

| Analysis | Dataset | Hemi. | *t*(175) | *p*-value | Cohen’s *d* |
| --- | --- | --- | --- | --- | --- |
| Actual category activity > non-category activity |  |  |  |  |  |
| EBA/FBA: body > non-body | Disc. | left | 21.03 | < 0.0001 | 1.59 |
| FFA/pSTS: face > non-face | Disc. | left | 18.42 | < 0.0001 | 1.39 |
| PPA/RSC: place > non-place | Disc. | left | 37.52 | < 0.0001 | 2.71 |
| LOC: tool > non-tool | Disc. | left | 13.9 | < 0.0001 | 1.05 |
| EBA/FBA: body > non-body | Repl. | left | 22.64 | < 0.0001 | 1.71 |
| FFA/pSTS: face > non-face | Repl. | left | 20.06 | < 0.0001 | 1.52 |
| PPA/RSC: place > non-place | Repl. | left | 34.26 | < 0.0001 | 2.49 |
| LOC: tool > non-tool | Repl. | left | 14.9 | < 0.0001 | 1.13 |
| EBA/FBA: body > non-body | Repl. | right | 28.39 | < 0.0001 | 2.15 |
| FFA/pSTS: face > non-face | Repl. | right | 23.08 | < 0.0001 | 1.74 |
| PPA/RSC: place > non-place | Repl. | right | 36.15 | < 0.0001 | 2.71 |
| LOC: tool > non-tool | Repl. | right | 8.61 | < 0.0001 | 0.65 |
| Mapped category activity > non-category activity |  |  |  |  |  |
| EBA/FBA: body > non-body | Disc. | left | 15.12 | < 0.0001 | 1.14 |
| FFA/pSTS: face > non-face | Disc. | left | 8.23 | < 0.0001 | 0.62 |
| PPA/RSC: place > non-place | Disc. | left | 25.49 | < 0.0001 | 1.82 |
| LOC: tool > non-tool | Disc. | left | 13.13 | < 0.0001 | 0.99 |
| EBA/FBA: body > non-body | Repl. | left | 15.47 | < 0.0001 | 1.17 |
| FFA/pSTS: face > non-face | Repl. | left | 9.62 | < 0.0001 | 0.73 |
| PPA/RSC: place > non-place | Repl. | left | 24.04 | < 0.0001 | 1.73 |
| LOC: tool > non-tool | Repl. | left | 13.48 | < 0.0001 | 1.02 |
| EBA/FBA: body > non-body | Repl. | right | 18.43 | < 0.0001 | 1.39 |
| FFA/pSTS: face > non-face | Repl. | right | 10.97 | < 0.0001 | 0.83 |
| PPA/RSC: place > non-place | Repl. | right | 24.72 | < 0.0001 | 1.82 |
| LOC: tool > non-tool | Repl. | right | 10.09 | < 0.0001 | 0.76 |

Number of permutations in max-T nonparametric permutation testing: 10,000. Disc. = discovery; repl. = replication; hemi. = hemisphere.
