## Supplementary material for "Distributed network flows generate localized category selectivity in human visual cortex": S3 Table

#### **S3 Table. Category selectivity scores and estimated percent distributed processing contribution to category selectivity in four functional complexes.**

| Analysis | Dataset | Hemi. | Score | max-T thresh. (*d.f.*) | *p*-value | Cohen’s *d* |
| --- | --- | --- | --- | --- | --- | --- |
| Mapped category selectivity  (null mean = 1.0) |  |  |  |  |  |  |
| EBA/FBA: body selectivity | Disc. | left | 1.28 | 6.69 (163) | < 0.00001 | 1.20 |
| FFA/pSTS: face selectivity | Disc. | left | 1.31 | 7.34 (165) | < 0.00001 | 0.75 |
| PPA/RSC: place selectivity | Disc. | left | 1.74 | 3.17 (167) | < 0.00001 | 1.56 |
| LOC: tool selectivity | Disc. | left | 1.16 | 9.19 (173) | < 0.00001 | 1.04 |
| EBA/FBA: body selectivity | Repl. | left | 1.31 | 6.32 (170) | < 0.00001 | 1.10 |
| FFA/pSTS: face selectivity | Repl. | left | 1.28 | 7.7 (164) | < 0.00001 | 0.83 |
| PPA/RSC: place selectivity | Repl. | left | 1.72 | 3.35 (161) | < 0.00001 | 1.55 |
| LOC: tool selectivity | Repl. | left | 1.14 | 9.24 (173) | < 0.00001 | 1.17 |
| EBA/FBA: body selectivity | Repl. | right | 1.39 | 5.78 (168) | < 0.00001 | 1.18 |
| FFA/pSTS: face selectivity | Repl. | right | 1.29 | 7.49 (162) | < 0.00001 | 0.98 |
| PPA/RSC: place selectivity | Repl. | right | 1.77 | 3.52 (164) | < 0.00001 | 1.46 |
| LOC: tool selectivity | Repl. | right | 1.11 | 9.67 (173) | < 0.00001 | 0.95 |
| Actual category selectivity  (null mean = 1.0) |  |  |  |  |  |  |
| EBA/FBA: body selectivity | Disc. | left | 1.5 | 6.69 (163) | < 0.00001 | 1.43 |
| FFA/pSTS: face selectivity | Disc. | left | 1.36 | 7.34 (165) | < 0.00001 | 0.95 |
| PPA/RSC: place selectivity | Disc. | left | 2.55 | 3.17 (167) | < 0.00001 | 1.75 |
| LOC: tool selectivity | Disc. | left | 1.2 | 9.19 (173) | < 0.00001 | 1.28 |
| EBA/FBA: body selectivity | Repl. | left | 1.59 | 6.32 (170) | < 0.00001 | 1.38 |
| FFA/pSTS: face selectivity | Repl. | left | 1.32 | 7.7 (164) | < 0.00001 | 1.10 |
| PPA/RSC: place selectivity | Repl. | left | 2.42 | 3.35 (161) | < 0.00001 | 1.71 |
| LOC: tool selectivity | Repl. | left | 1.2 | 9.24 (173) | < 0.00001 | 1.26 |
| EBA/FBA: body selectivity | Repl. | right | 1.7 | 5.78 (168) | < 0.00001 | 1.60 |
| FFA/pSTS: face selectivity | Repl. | right | 1.35 | 7.49 (162) | < 0.00001 | 1.16 |
| PPA/RSC: place selectivity | Repl. | right | 1.87 | 3.52 (164) | < 0.00001 | 1.87 |
| LOC: tool selectivity | Repl. | right | 1.15 | 9.67 (173) | < 0.00001 | 1.05 |
| Percent distributed contribution  (null mean = 50%) |  |  |  |  |  |  |
| EBA/FBA: body selectivity | Disc. | left | 85% | 5.66 (163) | < 0.00001 | 2.85 |
| FFA/pSTS: face selectivity | Disc. | left | 96% | 4.78 (165) | < 0.00001 | 2.03 |
| PPA/RSC: place selectivity | Disc. | left | 68% | 7.11 (167) | < 0.00001 | 1.23 |
| LOC: tool selectivity | Disc. | left | 97% | 5.2 (173) | < 0.00001 | 10.17 |
| EBA/FBA: body selectivity | Repl. | left | 82% | 6.04 (170) | < 0.00001 | 2.41 |
| FFA/pSTS: face selectivity | Repl. | left | 97% | 4.87 (164) | < 0.00001 | 2.64 |
| PPA/RSC: place selectivity | Repl. | left | 71% | 6.63 (161) | < 0.00001 | 1.32 |
| LOC: tool selectivity | Repl. | left | 95% | 5.26 (173) | < 0.00001 | 9.40 |
| EBA/FBA: body selectivity | Repl. | right | 82% | 6.01 (168) | < 0.00001 | 2.12 |
| FFA/pSTS: face selectivity | Repl. | right | 95% | 4.9 (162) | < 0.00001 | 2.91 |
| PPA/RSC: place selectivity | Repl. | right | 74% | 6.61 (164) | < 0.00001 | 1.58 |
| LOC: tool selectivity | Repl. | right | 97% | 5.14 (173) | < 0.00001 | 10.32 |

Number of permutations in max-T nonparametric permutation testing: 100,000. *d.f.* = degrees of freedom (see Methods for details on outlier removal procedure). Disc. = discovery; repl. = replication; thresh. = threshold; hemi. = hemisphere.
