## Supplementary material for "Distributed network flows generate localized category selectivity in human visual cortex": S4 Table

#### **S4 Table. Large-scale functional network activity flows contributing to category-specific responses in four functional complexes.**

| Analysis | Dataset | Hemi. | Contributing networks (sig.) | max-T thresh. (175) | *p*-value |
| --- | --- | --- | --- | --- | --- |
| Contributing network-mean activity flow products to: |  |  |  |  |  |
| EBA/FBA body responses | Disc. | left | VIS2, DAN | 3.41 | <0.0001 |
| FFA/pSTS face responses | Disc. | left | VIS2 | 3.4 | <0.0001 |
| PPA/RSC place responses | Disc. | left | VIS1, VIS2, DAN | 3.38 | <0.0001 |
| LOC tool responses | Disc. | left | VIS1, VIS2 | 3.4 | <0.0001 |
| EBA/FBA body responses | Repl. | left | VIS2, DAN | 3.39 | <0.0001 |
| FFA/pSTS face responses | Repl. | left | VIS2 | 3.36 | <0.0001 |
| PPA/RSC place responses | Repl. | left | VIS1, VIS2, DAN | 3.4 | <0.0001 |
| LOC tool responses | Repl. | left | VIS1, VIS2 | 3.37 | <0.0001 |
| EBA/FBA body responses | Repl. | right | VIS2, DAN | 3.41 | <0.0001 |
| FFA/pSTS face responses | Repl. | right | VIS2 | 3.35 | <0.0001 |
| PPA/RSC place responses | Repl. | right | VIS1, VIS2, DAN | 3.41 | <0.0001 |
| LOC tool responses | Repl. | right | VIS2 | 3.35 | <0.0001 |

Number of permutations in max-T nonparametric permutation testing: 10,000. sig. = significant. VIS1 = primary visual network; VIS2 = secondary visual network; DAN = dorsal attention network. These results corroborate results presented in Figs 4E-7E (right hemisphere discovery data; statistics reported in main text). Disc. = discovery; repl. = replication; thresh. = threshold; hemi. = hemisphere.
