## Supplementary material for "Distributed network flows generate localized category selectivity in human visual cortex": S5 Table

#### **S5 Table. Discovery dataset: variance explained per network in predicting cross-condition response profiles in left hemisphere complexes.**

| Source network | EBA/FBA partial R^2^ | EBA/FBA  rel. % | FFA/pSTS partial R^2^ | FFA/pSTS rel. % | PPA/RSC partial R^2^ | PPA/RSC  rel. % | LOC  partial R^2^ | LOC  rel. % |
| --- | --- | --- | --- | --- | --- | --- | --- | --- |
| VIS1 | 0.035 | 4.2% | 0.052 | 5.8% | 0.038 | 5.6% | 0.059 | 6.5% |
| VIS2 | 0.464 | 55.2%* | 0.362 | 40.0%* | 0.292 | 42.4%* | 0.706 | 77.6%* |
| SMN | 0.018 | 2.2% | 0.020 | 2.2% | 0.010 | 1.5% | 0.011 | 1.2% |
| CON | 0.021 | 2.5% | 0.031 | 3.4% | 0.015 | 2.2% | 0.015 | 1.7% |
| DAN | 0.134 | 15.9%* | 0.092 | 10.1%* | 0.073 | 10.6%* | 0.030 | 3.3% |
| LAN | 0.029 | 3.4% | 0.085 | 9.5% | 0.012 | 1.7% | 0.015 | 1.6% |
| FPN | 0.039 | 4.6% | 0.035 | 3.8% | 0.030 | 4.4% | 0.020 | 2.2% |
| AUD | 0.013 | 1.6% | 0.027 | 3.0% | 0.010 | 1.5% | 0.011 | 1.2% |
| DMN | 0.014 | 1.6% | 0.073 | 8.1% | 0.180 | 26.2%* | 0.013 | 1.5% |
| PMM | 0.046 | 5.4% | 0.070 | 7.7% | 0.012 | 1.8% | 0.014 | 1.6% |
| VMM | 0.024 | 2.9% | 0.052 | 5.8% | 0.011 | 1.6% | 0.012 | 1.3% |
| OAN | 0.005 | 0.6% | 0.005 | 0.6% | 0.004 | 0.5% | 0.003 | 0.3% |
| total | 0.841 | 100% | 0.904 | 100% | 0.688 | 100% | 0.910 | 100% |
| *t*(175) vs. 0.5 | 69.93 | n/a | 119.53 | n/a | 23.45 | n/a | 135.51 | n/a |
| *p*-value | 8.0x10^-130^ | n/a | 1.1x10^-169^ | n/a | 6.6x10^-56^ | n/a | 3.9x10^-179^ | n/a |

Source network = network-based source of explained variance in activity-flow-mapped activations across 24 conditions (i.e., the response profile). VIS1 = primary visual network; VIS2 = secondary visual network; SMN = somatomotor network; CON = cingulo-opercular network; DAN = dorsal attention network; LAN = language network; FPN = frontoparietal network; AUD = auditory network; DMN = default mode network; PMM = posterior multimodal network; VMM = ventral multimodal network; OAN = orbito-affective network. rel. % = percent of relative importance to the full model. Asterisks = statistically significant network contributions (*p* < 0.0001, number of permutations = 10,000). EBA/FBA max-T(175) = 3.41; FFA/pSTS max-T(175) = 3.42; PPA/RSC max-T(175) = 3.42; LOC max-T(175) = 3.39. Statistical results listed in the bottom two rows refer to 1 sample t-testing of the total R^2^ value for each model versus 0.5, which assesses whether the mapped response profile for a given functional complex explains more than 50% of the variance in the actual response profile. This provides evidence that distributed processes (as captured by activity flow mapping) are the dominant influence in generating a given functional complexes activations to a diverse set of cognitive domains. n/a = not applicable. These results corroborate results presented in Figs 4F-7F (right hemisphere discovery data; statistics reported in main text).
