## Supplementary material for "Distributed network flows generate localized category selectivity in human visual cortex": S6 Table

#### **S6 Table. Replication dataset: variance explained per network in predicting cross-condition response profiles in left hemisphere complexes.**

| Source network | EBA/FBA partial R^2^ | EBA/FBA  rel. % | FFA/pSTS partial R^2^ | FFA/pSTS rel. % | PPA/RSC partial R^2^ | PPA/RSC  rel. % | LOC  partial R^2^ | LOC  rel. % |
| --- | --- | --- | --- | --- | --- | --- | --- | --- |
| VIS1 | 0.039 | 4.6% | 0.044 | 4.9% | 0.041 | 5.9% | 0.062 | 6.9% |
| VIS2 | 0.452 | 54.0%* | 0.349 | 38.9%* | 0.287 | 42.1%* | 0.698 | 77.0%* |
| SMN | 0.019 | 2.3% | 0.018 | 2.0% | 0.011 | 1.6% | 0.009 | 1.0% |
| CON | 0.017 | 2.0% | 0.037 | 4.1% | 0.016 | 2.4% | 0.014 | 1.5% |
| DAN | 0.133 | 15.9%* | 0.105 | 11.7%* | 0.075 | 11.0%* | 0.031 | 3.4% |
| LAN | 0.029 | 3.4% | 0.089 | 9.9%* | 0.012 | 1.8% | 0.016 | 1.7% |
| FPN | 0.035 | 4.2% | 0.035 | 3.8% | 0.028 | 4.1% | 0.019 | 2.1% |
| AUD | 0.017 | 2.0% | 0.023 | 2.6% | 0.012 | 1.7% | 0.012 | 1.4% |
| DMN | 0.015 | 1.8% | 0.084 | 9.3% | 0.174 | 25.5%* | 0.013 | 1.4% |
| PMM | 0.054 | 6.4% | 0.053 | 5.9% | 0.011 | 1.6% | 0.014 | 1.6% |
| VMM | 0.024 | 2.8% | 0.056 | 6.2% | 0.009 | 1.2% | 0.014 | 1.5% |
| OAN | 0.004 | 0.5% | 0.006 | 0.6% | 0.007 | 1.0% | 0.004 | 0.5% |
| total | 0.837 | 100% | 0.899 | 100% | 0.682 | 100% | 0.906 | 100% |
| *t*(175) vs. 0.5 | 68.96 | n/a | 97.88 | n/a | 22.93 | n/a | 124.21 | n/a |
| *p*-value | 8.5x10^-129^ | n/a | 9.7x10^-155^ | n/a | 1.3x10^-54^ | n/a | 1.4x10^-172^ | n/a |

Source network = network-based source of explained variance in activity-flow-mapped activations across 24 conditions (i.e., the response profile). VIS1 = primary visual network; VIS2 = secondary visual network; SMN = somatomotor network; CON = cingulo-opercular network; DAN = dorsal attention network; LAN = language network; FPN = frontoparietal network; AUD = auditory network; DMN = default mode network; PMM = posterior multimodal network; VMM = ventral multimodal network; OAN = orbito-affective network. rel. % = percent of relative importance to the full model. Asterisks = statistically significant network contributions (*p* < 0.0001, number of permutations = 10,000). EBA/FBA max-T(175) = 3.34; FFA/pSTS max-T(175) = 3.4; PPA/RSC max-T(175) = 3.35; LOC max-T(175) = 3.38. Statistical results listed in the bottom two rows refer to 1 sample t-testing of the total R^2^ value for each model versus 0.5, which assesses whether the mapped response profile for a given functional complex explains more than 50% of the variance in the actual response profile. This provides evidence that distributed processes (as captured by activity flow mapping) are the dominant influence in generating a given functional complexes activations to a diverse set of cognitive domains. n/a = not applicable. These results corroborate results presented in Figs 4F-7F (right hemisphere discovery data; statistics reported in main text).
