## Supplementary material for "Distributed network flows generate localized category selectivity in human visual cortex": S7 Table

#### **S7 Table. Replication dataset: variance explained per network in predicting cross-condition response profiles in right hemisphere complexes.**

| Source network | EBA/FBA partial R^2^ | EBA/FBA  rel. % | FFA/pSTS partial R^2^ | FFA/pSTS rel. % | PPA/RSC partial R^2^ | PPA/RSC  rel. % | LOC  partial R^2^ | LOC  rel. % |
| --- | --- | --- | --- | --- | --- | --- | --- | --- |
| VIS1 | 0.0333 | 4.06% | 0.0331 | 3.73% | 0.0492 | 7.14% | 0.0494 | 5.51% |
| VIS2 | 0.4791 | 58.48%* | 0.3688 | 41.58%* | 0.3239 | 47.00%* | 0.7077 | 78.91%* |
| SMN | 0.0213 | 2.60% | 0.0205 | 2.31% | 0.0091 | 1.32% | 0.0097 | 1.08% |
| CON | 0.0307 | 3.75% | 0.0328 | 3.70% | 0.0148 | 2.15% | 0.0157 | 1.75% |
| DAN | 0.0869 | 10.61%* | 0.0902 | 10.17%* | 0.0922 | 13.38%* | 0.0305 | 3.40% |
| LAN | 0.0265 | 3.23% | 0.0666 | 7.51% | 0.0108 | 1.57% | 0.0127 | 1.42% |
| FPN | 0.0212 | 2.59% | 0.0401 | 4.52% | 0.0253 | 3.67% | 0.0208 | 2.32% |
| AUD | 0.0142 | 1.73% | 0.0228 | 2.57% | 0.0115 | 1.67% | 0.0111 | 1.24% |
| DMN | 0.0145 | 1.77% | 0.0773 | 8.71%* | 0.1277 | 18.53%* | 0.0107 | 1.19% |
| PMM | 0.0657 | 8.02%* | 0.072 | 8.12%* | 0.0092 | 1.33% | 0.0114 | 1.27% |
| VMM | 0.0216 | 2.64% | 0.0565 | 6.37% | 0.0118 | 1.71% | 0.0123 | 1.37% |
| OAN | 0.0042 | 0.51% | 0.0063 | 0.71% | 0.0037 | 0.54% | 0.0048 | 0.54% |
| total | 0.819 | 100% | 0.887 | 100% | 0.689 | 100% | 0.897 | 100% |
| *t*(175) vs. 0.5 | 52.25 | n/a | 80.42 | n/a | 22.54 | n/a | 104.55 | n/a |
| *p*-value | 1.1x10^-108^ | n/a | 3.9x10^-140^ | n/a | 1.3x10^-53^ | n/a | 1.2x10^-159^ | n/a |

Source network = network-based source of explained variance in activity-flow-mapped activations across 24 conditions (i.e., the response profile). VIS1 = primary visual network; VIS2 = secondary visual network; SMN = somatomotor network; CON = cingulo-opercular network; DAN = dorsal attention network; LAN = language network; FPN = frontoparietal network; AUD = auditory network; DMN = default mode network; PMM = posterior multimodal network; VMM = ventral multimodal network; OAN = orbito-affective network. rel. % = percent of relative importance to the full model. Asterisks = statistically significant network contributions (*p* < 0.0001, number of permutations = 10,000). EBA/FBA max-T(175) = 3.41; FFA/pSTS max-T(175) = 3.39; PPA/RSC max-T(175) = 3.42; LOC max-T(175) = 3.38. Statistical results listed in the bottom two rows refer to 1 sample t-testing of the total R^2^ value for each model versus 0.5, which assesses whether the mapped response profile for a given functional complex explains more than 50% of the variance in the actual response profile. This provides evidence that distributed processes (as captured by activity flow mapping) are the dominant influence in generating a given functional complexes activations to a diverse set of cognitive domains. n/a = not applicable. These results corroborate results presented in Figs 4F-7F (right hemisphere discovery data; statistics reported in main text).
