## Supplementary material for "Distributed network flows generate localized category selectivity in human visual cortex": S8 Table

#### **S8 Table. Null connectivity fingerprint models, right hemisphere functional complexes and replication dataset.**

| Analysis | Dataset | *t*(175) | *p*-value | Cohen’s *d* |
| --- | --- | --- | --- | --- |
| True model body selectivity > null model body selectivity, when EBA/FBA rsFC substituted with: |  |  |  |  |
| face complex connectivity fingerprint: FFA/pSTS rsFC | Repl. | 5.92 | 1.7x10^-08^ | 0.63 |
| place complex connectivity fingerprint: PPA/RSC rsFC | Repl. | 6.47 | 9.7x10^-10^ | 0.69 |
| tool complex connectivity fingerprint: LOC rsFC | Repl. | 9.27 | 6.8x10^-17^ | 0.98 |
| True model face selectivity > null model face selectivity, when FFA/pSTS rsFC substituted with: |  |  |  |  |
| body complex connectivity fingerprint: EBA/FBA rsFC | Repl. | 7.2 | 1.7x10^-11^ | 0.78 |
| place complex connectivity fingerprint: PPA/RSC rsFC | Repl. | 12.6 | 2.3x10^-26^ | 1.35 |
| tool complex connectivity fingerprint: LOC rsFC | Repl. | 9.15 | 1.5x10^-16^ | 1.00 |
| True model place selectivity > null model place selectivity, when PPA/RSC rsFC substituted with: |  |  |  |  |
| body complex connectivity fingerprint: EBA/FBA rsFC | Repl. | 15.0 | 2.8x10^-33^ | 1.61 |
| face complex connectivity fingerprint: FFA/pSTS rsFC | Repl. | 15.5 | 1.2x10^-34^ | 1.61 |
| tool complex connectivity fingerprint: LOC rsFC | Repl. | 12.5 | 5.7x10^-26^ | 1.33 |
| True model tool selectivity > null model tool selectivity, when LOC rsFC substituted with: |  |  |  |  |
| body complex connectivity fingerprint: EBA/FBA rsFC | Repl. | 5.06 | 1.1x10^-06^ | 0.55 |
| face complex connectivity fingerprint: FFA/pSTS rsFC | Repl. | 7.62 | 1.6x10^-12^ | 0.82 |
| place complex connectivity fingerprint: PPA/RSC rsFC | Repl. | 3.62 | 0.0004 | 0.39 |

In each analysis, the true model category selectivity scores (i.e., activity-flow-mapped with true connectivity fingerprint) were compared to the null model category selectivity scores (i.e., activity-flow-mapped with substituted connectivity fingerprints) across participants (paired samples t-test). Repl. = replication.
