## Supplementary material for "Distributed network flows generate localized category selectivity in human visual cortex": S9 Table

#### **S9 Table. Null connectivity fingerprint models, left hemisphere functional complexes.**

| Analysis | Dataset | *t*(175) | *p*-value | Cohen’s *d* |
| --- | --- | --- | --- | --- |
| True model body selectivity > null model body selectivity, when EBA/FBA rsFC substituted with: |  |  |  |  |
| face complex connectivity fingerprint: FFA/pSTS rsFC | Disc. | 3.78 | 0.0002 | 0.36 |
| place complex connectivity fingerprint: PPA/RSC rsFC | Disc. | 2.77 | 0.006 | 0.28 |
| tool complex connectivity fingerprint: LOC rsFC | Disc. | 11.16 | 3.5x10^-22^ | 1.12 |
| face complex connectivity fingerprint: FFA/pSTS rsFC | Repl. | 4.55 | 1.0x10^-05^ | 0.48 |
| place complex connectivity fingerprint: PPA/RSC rsFC | Repl. | 6.07 | 7.8x10^-09^ | 0.64 |
| tool complex connectivity fingerprint: LOC rsFC | Repl. | 10.47 | 3.1x10^-20^ | 1.10 |
| True model face selectivity > null model face selectivity, when FFA/pSTS rsFC substituted with: |  |  |  |  |
| body complex connectivity fingerprint: EBA/FBA rsFC | Disc. | 9.27 | 6.7x10^-17^ | 0.94 |
| place complex connectivity fingerprint: PPA/RSC rsFC | Disc. | 13.52 | 5.5x10^-29^ | 1.39 |
| tool complex connectivity fingerprint: LOC rsFC | Disc. | 9.84 | 1.9x10^-18^ | 1.03 |
| body complex connectivity fingerprint: EBA/FBA rsFC | Repl. | 6.61 | 4.5x10^-10^ | 0.73 |
| place complex connectivity fingerprint: PPA/RSC rsFC | Repl. | 13.15 | 6.6x10^-28^ | 1.40 |
| tool complex connectivity fingerprint: LOC rsFC | Repl. | 9.53 | 1.3x10^-17^ | 1.03 |
| True model place selectivity > null model place selectivity, when PPA/RSC rsFC substituted with: |  |  |  |  |
| body complex connectivity fingerprint: EBA/FBA rsFC | Disc. | 13.37 | 1.5x10-^28^ | 1.4 |
| face complex connectivity fingerprint: FFA/pSTS rsFC | Disc. | 14.43 | 1.3x10^-31^ | 1.44 |
| tool complex connectivity fingerprint: LOC rsFC | Disc. | 11.1 | 5.1x10^-22^ | 1.14 |
| body complex connectivity fingerprint: EBA/FBA rsFC | Repl. | 14.5 | 8.3x10^-32^ | 1.51 |
| face complex connectivity fingerprint: FFA/pSTS rsFC | Repl. | 15.61 | 5.7x10^-35^ | 1.56 |
| tool complex connectivity fingerprint: LOC rsFC | Repl. | 11.9 | 2.6x10^-24^ | 1.26 |
| True model tool selectivity > null model tool selectivity, when LOC rsFC substituted with: |  |  |  |  |
| body complex connectivity fingerprint: EBA/FBA rsFC | Disc. | 4.67 | 6.0x10^-06^ | 0.24 |
| face complex connectivity fingerprint: FFA/pSTS rsFC | Disc. | 3.68 | 0.0003 | 0.35 |
| place complex connectivity fingerprint: PPA/RSC rsFC | Disc. | 2.44 | 0.02 | 0.25 |
| body complex connectivity fingerprint: EBA/FBA rsFC | Repl. | 2.19 | 0.03 | 0.23 |
| face complex connectivity fingerprint: FFA/pSTS rsFC | Repl. | 4.43 | 1.7x10^-05^ | 0.48 |
| place complex connectivity fingerprint: PPA/RSC rsFC | Repl. | 3.09 | 0.002 | 0.33 |

In each analysis, the true model category selectivity scores (i.e., activity-flow-mapped with true connectivity fingerprint) were compared to the null model body selectivity scores (i.e., activity-flow-mapped with substituted connectivity fingerprints) across participants (paired samples t-test). Disc. = discovery; repl. = replication.
