## Supplementary material for "Distributed network flows generate localized category selectivity in human visual cortex": S10 Table

#### **S10 Table. Category selectivity generated from stimulus-driven activity flow processes for left hemisphere functional complexes and the replication dataset.**

| Analysis | Dataset | Hemi. | Score | max-T thresh. (*d.f.*) | *p*-value | Cohen’s *d* |
| --- | --- | --- | --- | --- | --- | --- |
| Mapped category selectivity  (null mean = 1.0) |  |  |  |  |  |  |
| EBA/FBA: body selectivity | Disc. | left | 1.21 | 8.9 (174) | <0.00001 | 0.71 |
| FFA/pSTS: face selectivity | Disc. | left | 1.05 | 8.48 (172) | n.s. | 0.23 |
| PPA/RSC: place selectivity | Disc. | left | 1.07 | 5.02 (172) | <0.00001 | 0.39 |
| LOC: tool selectivity | Disc. | left | 1.04 | 9.38 (175) | n.s. | 0.16 |
| EBA/FBA: body selectivity | Repl. | right | 1.32 | 7.27 (174) | <0.00001 | 1.63 |
| FFA/pSTS: face selectivity | Repl. | right | 1.16 | 8.19 (175) | <0.00001 | 0.83 |
| PPA/RSC: place selectivity | Repl. | right | 1.27 | 5.4 (173) | <0.00001 | 1.44 |
| LOC: tool selectivity | Repl. | right | 1.36 | 7.87 (171) | <0.00001 | 1.73 |
| EBA/FBA: body selectivity | Repl. | left | 1.3 | 8.47 (174) | <0.00001 | 1.96 |
| FFA/pSTS: face selectivity | Repl. | left | 1.12 | 8.58 (175) | <0.00001 | 0.74 |
| PPA/RSC: place selectivity | Repl. | left | 1.27 | 5.68 (174) | <0.00001 | 2.0 |
| LOC: tool selectivity | Repl. | left | 1.31 | 8.37 (174) | <0.00001 | 2.29 |
| Actual category selectivity  (null mean = 1.0) |  |  |  |  |  |  |
| EBA/FBA: body selectivity | Disc. | left | 1.17 | 8.9 (174) | <0.00001 | 1.19 |
| FFA/pSTS: face selectivity | Disc. | left | 1.28 | 8.48 (172) | <0.00001 | 1.53 |
| PPA/RSC: place selectivity | Disc. | left | 1.78 | 5.02 (172) | <0.00001 | 2.79 |
| LOC: tool selectivity | Disc. | left | 1.19 | 9.38 (175) | <0.00001 | 1.33 |
| EBA/FBA: body selectivity | Repl. | right | 1.45 | 7.27 (174) | <0.00001 | 1.88 |
| FFA/pSTS: face selectivity | Repl. | right | 1.33 | 8.19 (175) | <0.00001 | 1.68 |
| PPA/RSC: place selectivity | Repl. | right | 1.83 | 5.4 (173) | <0.00001 | 2.29 |
| LOC: tool selectivity | Repl. | right | 1.24 | 7.87 (171) | <0.00001 | 1.59 |
| EBA/FBA: body selectivity | Repl. | left | 1.18 | 8.47 (174) | <0.00001 | 1.23 |
| FFA/pSTS: face selectivity | Repl. | left | 1.28 | 8.58 (175) | <0.00001 | 1.53 |
| PPA/RSC: place selectivity | Repl. | left | 1.77 | 5.68 (174) | <0.00001 | 2.36 |
| LOC: tool selectivity | Repl. | left | 1.19 | 8.37 (174) | <0.00001 | 1.29 |

Number of permutations in max-T nonparametric permutation testing: 100,000. *d.f.* = degrees of freedom (see Methods for details on outlier removal procedure). This analysis corresponds to category selectivity generated from V1-initialized, stimulus-driven activity flow processes (Figs 1B and 9; Step 1 of Fig 11B). Disc. = discovery; repl. = replication; hemi. = hemisphere; thresh. = threshold; n.s. = not significant.
