## Supplementary material for "Distributed network flows generate localized category selectivity in human visual cortex": S11 Table

#### **S11 Table. Actual category selectivity exhibited by V1 for the replication dataset.**

| Analysis | Dataset | Score | *t(*175) | *p*-value | Cohen’s *d* |
| --- | --- | --- | --- | --- | --- |
| EBA/FBA: body selectivity | Replication | 0.62 | -16.28 | n.s. | -1.24 |
| FFA/pSTS: face selectivity | Replication | 0.56 | -27.93 | n.s. | -2.14 |
| PPA/RSC: place selectivity | Replication | 0.61 | -19.28 | n.s. | -1.47 |
| LOC: tool selectivity | Replication | 064 | -14.45 | n.s. | -1.1 |

As in the main text (reporting the discovery dataset results), bilateral V1 (vertex-level data, see Methods) itself did not exhibit significant category selectivity for any of the four visual categories. Given this observation, it is likely that intrinsic connectivity patterns (between V1 and functional complexes of interest), and not source activation patterns in V1, were key to stimulus-driven activity flow processes being able to generate visual category selectivity.
