## Supplementary material for "Distributed network flows generate localized category selectivity in human visual cortex": S12 Table

#### **S12 Table. Stimulus-driven category selectivity generated via activity flow mapping is significantly greater than with a null network architecture.**

| Analysis | Dataset | Hemisphere | *t*(175) | *p*-value | Cohen’s *d* |
| --- | --- | --- | --- | --- | --- |
| EBA/FBA: body selectivity | Discovery | left | 3.84 | 8.8x10^-5^ | 0.29 |
| FFA/pSTS: face selectivity | Discovery | left | 10.8 | 2.3x10^-21^ | 0.82 |
| PPA/RSC: place selectivity | Discovery | left | 3.27 | 6.6x10^-4^ | 0.25 |
| LOC: tool selectivity | Discovery | left | -0.68 | n.s. | -0.05 |
| EBA/FBA: body selectivity | Replication | right | 11.93 | 1.3x10^-24^ | 0.91 |
| FFA/pSTS: face selectivity | Replication | right | 8.49 | 4.5x10^-15^ | 0.65 |
| PPA/RSC: place selectivity | Replication | right | 10.74 | 3x10^-21^ | 0.82 |
| LOC: tool selectivity | Replication | right | 12.28 | 1.3x10^-25^ | 0.93 |
| EBA/FBA: body selectivity | Replication | left | 7.65 | 6.7x10^-13^ | 0.58 |
| FFA/pSTS: face selectivity | Replication | left | 8.79 | 7.4x10^-16^ | 0.67 |
| PPA/RSC: place selectivity | Replication | left | 13.58 | 2.4x10^-29^ | 1.03 |
| LOC: tool selectivity | Replication | left | 12.19 | 2.2x10^-25^ | 0.93 |

In each analysis, the true model category selectivity scores (i.e., via stimulus-driven activity flow mapping with true connectivity fingerprint with V1) were compared to the null model (Fig 10C) category selectivity scores (i.e., activity flow mapping with randomly shuffled connectivity fingerprints, see Methods) across participants (paired samples t-test). N.s. = not significant.
