## Supplementary material for "Distributed network flows generate localized category selectivity in human visual cortex": S13 Table

#### **S13 Table. Category selectivity via stimulus-driven processes further shaped by VIS subsystem network interactions (step 2), left hemisphere functional complexes and the replication dataset.**

| Analysis | Dataset | Hemi. | Score | max-T thresh. (*d.f.*) | *p*-value | Cohen’s *d* |
| --- | --- | --- | --- | --- | --- | --- |
| Mapped category selectivity  (null mean = 1.0) |  |  |  |  |  |  |
| EBA/FBA: body selectivity | Disc. | left | 1.13 | 9.46 (174) | <0.00001 | 0.84 |
| FFA/pSTS: face selectivity | Disc. | left | 0.96 | 8.49 (172) | n.s. | -0.26 |
| PPA/RSC: place selectivity | Disc. | left | 1.25 | 5.68 (170) | <0.00001 | 1.75 |
| LOC: tool selectivity | Disc. | left | 1.24 | 8.68 (170) | <0.00001 | 1.7 |
| EBA/FBA: body selectivity | Repl. | right | 1.1 | 7.2 (170) | <0.00001 | 0.59 |
| FFA/pSTS: face selectivity | Repl. | right | 0.97 | 8.17 (175) | n.s. | -0.18 |
| PPA/RSC: place selectivity | Repl. | right | 1.22 | 5.33 (168) | <0.00001 | 1.42 |
| LOC: tool selectivity | Repl. | right | 1.3 | 8.12 (167) | <0.00001 | 1.48 |
| EBA/FBA: body selectivity | Repl. | left | 1.07 | 9.42 (172) | n.s. | 0.47 |
| FFA/pSTS: face selectivity | Repl. | left | 0.92 | 8.56 (175) | n.s. | -0.65 |
| PPA/RSC: place selectivity | Repl. | left | 1.22 | 6.63 (172) | <0.00001 | 1.37 |
| LOC: tool selectivity | Repl. | left | 1.24 | 8.66 (170) | <0.00001 | 1.54 |
| Actual category selectivity  (null mean = 1.0) |  |  |  |  |  |  |
| EBA/FBA: body selectivity | Disc. | left | 1.16 | 9.46 (174) | <0.00001 | 1.18 |
| FFA/pSTS: face selectivity | Disc. | left | 1.28 | 8.49 (172) | <0.00001 | 1.53 |
| PPA/RSC: place selectivity | Disc. | left | 1.76 | 5.68 (170) | <0.00001 | 2.87 |
| LOC: tool selectivity | Disc. | left | 1.18 | 8.68 (170) | <0.00001 | 1.31 |
| EBA/FBA: body selectivity | Repl. | right | 1.46 | 7.2 (170) | <0.00001 | 1.91 |
| FFA/pSTS: face selectivity | Repl. | right | 1.33 | 8.17 (175) | <0.00001 | 1.68 |
| PPA/RSC: place selectivity | Repl. | right | 1.83 | 5.33 (168) | <0.00001 | 2.32 |
| LOC: tool selectivity | Repl. | right | 1.24 | 8.12 (167) | <0.00001 | 1.6 |
| EBA/FBA: body selectivity | Repl. | left | 1.18 | 9.42 (172) | <0.00001 | 1.2 |
| FFA/pSTS: face selectivity | Repl. | left | 1.28 | 8.56 (175) | <0.00001 | 1.53 |
| PPA/RSC: place selectivity | Repl. | left | 1.78 | 6.63 (172) | <0.00001 | 2.37 |
| LOC: tool selectivity | Repl. | left | 1.19 | 8.66 (170) | <0.00001 | 1.28 |

Number of permutations in max-T nonparametric permutation testing: 100,000. *d.f.* = degrees of freedom (see Methods for details on outlier removal procedure). This analysis corresponds to category selectivity generated from V1-initialized, stimulus-driven activity flow processes that are further shaped by one step of VIS subsystem network interactions (Figs 1C and 11). Disc. = discovery; repl. = replication; hemi. = hemisphere; thresh. = threshold; n.s. = not significant.
