## Supplementary material for "Distributed network flows generate localized category selectivity in human visual cortex": S14 Table

#### **S14 Table. Category selectivity via stimulus-driven processes further shaped by VIS subsystem network interactions (step 3), left hemisphere functional complexes and the replication dataset.**

| Analysis | Dataset | Hemi. | Score | max-T thresh. (*d.f.*) | *p*-value | Cohen’s *d* |
| --- | --- | --- | --- | --- | --- | --- |
| Mapped category selectivity  (null mean = 1.0) |  |  |  |  |  |  |
| EBA/FBA: body selectivity | Disc. | left | 1.03 | 9.62 (175) | n.s. | 0.23 |
| FFA/pSTS: face selectivity | Disc. | left | 0.89 | 8.47 (172) | n.s. | -0.83 |
| PPA/RSC: place selectivity | Disc. | left | 1.15 | 5.68 (172) | <0.00001 | 1.02 |
| LOC: tool selectivity | Disc. | left | 1.17 | 9.14 (173) | <0.00001 | 1.2 |
| EBA/FBA: body selectivity | Repl. | right | 0.99 | 7.26 (172) | n.s. | -0.1 |
| FFA/pSTS: face selectivity | Repl. | right | 0.86 | 8.18 (175) | n.s. | -0.13 |
| PPA/RSC: place selectivity | Repl. | right | 1.15 | 5.39 (170) | <0.00001 | 0.95 |
| LOC: tool selectivity | Repl. | right | 1.2 | 8.62 (168) | <0.00001 | 1.32 |
| EBA/FBA: body selectivity | Repl. | left | 1.0 | 9.43 (171) | n.s. | -0.02 |
| FFA/pSTS: face selectivity | Repl. | left | 0.86 | 8.58 (175) | n.s. | -0.99 |
| PPA/RSC: place selectivity | Repl. | left | 1.16 | 5.66 (172) | <0.00001 | 1.01 |
| LOC: tool selectivity | Repl. | left | 1.2 | 8.91 (168) | <0.00001 | 1.38 |
| Actual category selectivity  (null mean = 1.0) |  |  |  |  |  |  |
| EBA/FBA: body selectivity | Disc. | left | 1.16 | 9.62 (175) | <0.00001 | 1.18 |
| FFA/pSTS: face selectivity | Disc. | left | 1.28 | 8.47 (172) | <0.00001 | 1.53 |
| PPA/RSC: place selectivity | Disc. | left | 1.77 | 5.68 (172) | <0.00001 | 2.81 |
| LOC: tool selectivity | Disc. | left | 1.19 | 9.14 (173) | <0.00001 | 1.33 |
| EBA/FBA: body selectivity | Repl. | right | 1.45 | 7.26 (172) | <0.00001 | 1.89 |
| FFA/pSTS: face selectivity | Repl. | right | 1.33 | 8.18 (175) | <0.00001 | 1.68 |
| PPA/RSC: place selectivity | Repl. | right | 1.82 | 5.39 (170) | <0.00001 | 2.34 |
| LOC: tool selectivity | Repl. | right | 1.24 | 8.62 (168) | <0.00001 | 1.58 |
| EBA/FBA: body selectivity | Repl. | left | 1.17 | 9.43 (171) | <0.00001 | 1.23 |
| FFA/pSTS: face selectivity | Repl. | left | 1.28 | 8.58 (175) | <0.00001 | 1.53 |
| PPA/RSC: place selectivity | Repl. | left | 1.77 | 5.66 (172) | <0.00001 | 2.38 |
| LOC: tool selectivity | Repl. | left | 1.19 | 8.91 (168) | <0.00001 | 1.28 |

Number of permutations in max-T nonparametric permutation testing: 100,000. *d.f.* = degrees of freedom (see Methods for details on outlier removal procedure). This analysis corresponds to category selectivity generated from V1-initialized, stimulus-driven activity flow processes that are further shaped by two steps of VIS subsystem network interactions (Figs 1C and 11). Disc. = discovery; repl. = replication; hemi. = hemisphere; thresh. = threshold; n.s. = not significant.
