## Supplementary material for "Distributed network flows generate localized category selectivity in human visual cortex": S15 Table

#### **S15 Table. Category selectivity via stimulus-driven activity flow processes further shaped by whole-cortex, fully distributed network interactions.**

| Analysis | Dataset | Hemi. | Score | max-T thresh. (*d.f.*) | *p*-value | Cohen’s *d* |
| --- | --- | --- | --- | --- | --- | --- |
| Mapped category selectivity  (null mean = 1.0) |  |  |  |  |  |  |
| EBA/FBA: body selectivity | Disc. | left | 1.32 | 6.29 (169) | <0.00001 | 0.74 |
| FFA/pSTS: face selectivity | Disc. | left | 1.29 | 8.52 (165) | <0.00001 | 0.67 |
| PPA/RSC: place selectivity | Disc. | left | 1.77 | 4.45 (167) | <0.00001 | 1.13 |
| LOC: tool selectivity | Disc. | left | 1.1 | 10.4 (165) | <0.00001 | 0.81 |
| EBA/FBA: body selectivity | Repl. | right | 1.41 | 5.67 (168) | <0.00001 | 0.94 |
| FFA/pSTS: face selectivity | Repl. | right | 1.31 | 8.01 (163) | <0.00001 | 0.63 |
| PPA/RSC: place selectivity | Repl. | right | 1.76 | 4.86 (160) | <0.00001 | 1.13 |
| LOC: tool selectivity | Repl. | right | 1.09 | 6.36 (167) | <0.00001 | 0.49 |
| EBA/FBA: body selectivity | Repl. | left | 1.26 | 6.12 (165) | <0.00001 | 0.82 |
| FFA/pSTS: face selectivity | Repl. | left | 1.28 | 8.22 (163) | <0.00001 | 0.66 |
| PPA/RSC: place selectivity | Repl. | left | 1.75 | 4.63 (162) | <0.00001 | 1.12 |
| LOC: tool selectivity | Repl. | left | 1.11 | 10.4 (164) | <0.00001 | 0.83 |
| Actual category selectivity  (null mean = 1.0) |  |  |  |  |  |  |
| EBA/FBA: body selectivity | Disc. | left | 1.57 | 6.29 (169) | <0.00001 | 1.16 |
| FFA/pSTS: face selectivity | Disc. | left | 1.39 | 8.52 (165) | <0.00001 | 0.89 |
| PPA/RSC: place selectivity | Disc. | left | 2.67 | 4.45 (167) | <0.00001 | 1.5 |
| LOC: tool selectivity | Disc. | left | 1.2 | 10.4 (165) | <0.00001 | 1.32 |
| EBA/FBA: body selectivity | Repl. | right | 1.72 | 5.67 (168) | <0.00001 | 1.61 |
| FFA/pSTS: face selectivity | Repl. | right | 1.36 | 8.01 (163) | <0.00001 | 1.19 |
| PPA/RSC: place selectivity | Repl. | right | 2.44 | 4.86 (160) | <0.00001 | 1.65 |
| LOC: tool selectivity | Repl. | right | 1.15 | 6.36 (167) | <0.00001 | 1.06 |
| EBA/FBA: body selectivity | Repl. | left | 1.61 | 6.12 (165) | <0.00001 | 1.39 |
| FFA/pSTS: face selectivity | Repl. | left | 1.33 | 8.22 (163) | <0.00001 | 1.11 |
| PPA/RSC: place selectivity | Repl. | left | 2.52 | 4.63 (162) | <0.00001 | 1.41 |
| LOC: tool selectivity | Repl. | left | 1.19 | 10.4 (164) | <0.00001 | 1.29 |

Number of permutations in max-T nonparametric permutation testing: 100,000. *d.f.* = degrees of freedom (see Methods for details on outlier removal procedure). This analysis corresponds to category selectivity generated from V1-initialized, stimulus-driven activity flow processes that are further shaped by fully distributed network interactions (Figs 1D and 12). Disc. = discovery; repl. = replication; hemi. = hemisphere; thresh. = threshold.
